# Non-self RNA rewires IFNβ signaling: A mathematical model of the innate immune response

**DOI:** 10.1101/2022.01.30.478391

**Authors:** Zbigniew Korwek, Maciej Czerkies, Joanna Jaruszewicz-Błońska, Wiktor Prus, Ilona Kosiuk, Marek Kochańczyk, Tomasz Lipniacki

## Abstract

Type I interferons (IFNs) are key coordinators of the innate immune response to viral infection, which through activation of STAT1/2 in bystander cells induce the expression of IFN-stimulated genes (ISGs). The complex system-level mechanisms of IFN signaling are however not well understood. Here, we show that in cells transfected with an analog of viral RNA, poly(I:C), transcriptional activity of STAT1/2 is terminated due to depletion of the interferon β (IFNβ) receptor, IFNAR. Two ISGs, RNase L and PKR, not only hinder replenishment of IFNAR, but also suppress negative regulators of IRF3 and NF-κB, consequently promoting IFNβ transcription. We incorporated these findings into a comprehensive mathematical model of innate immunity. By coupling signaling through the IRF3/NF-κB and STAT1/2 pathways with the activity of RNase L and PKR, the model explains how poly(I:C) switches the transcriptional program from STAT1/2-induced to IRF3/NF-κB-induced, turning IFNβ-responding cells to IFNβ-secreting cells.

**One-sentence summary:** A computational model explains how non-self RNA turns cells from IFNβ- responders to IFNβ-producers.

## INTRODUCTION

The antiviral innate immune response is coordinated by interferon (IFN) signaling. Some virus-infected cells produce and secrete type I and type III IFNs to warn bystander cells about the threat (*1, 2*). These IFNs are expressed due to joint transcriptional activity of NF-κB and IRF3 (*3–5*), activated upon recognition of non-self RNA by intracellular receptors such as RIG-I in cooperation with MAVS (*6–9*). Secreted IFNs trigger transcriptional activity of STAT1 and STAT2 (by phosphorylation on Tyr^701^ and Tyr^690^, respectively), inducing expression of interferon-stimulated genes (ISGs), which encode potent antiviral proteins such as OAS1/2/3/L, PKR, and RIG-I (*10*). Upon detection of non-self RNA, OAS proteins activate RNase L, which degrades both viral and cellular RNA (*11*), while PKR phosphorylates eIF2α (on Ser^51^), which then inhibits translation (*12*). Of note, to prevent global shutdown of biosynthesis and enable replication, many viruses express inhibitors that counteract activation of RNase L or prevent the eIF2α phosphorylation-dependent translation arrest (*13–15*). Overall, through both autocrine and paracrine signaling, mediating, respectively, positive feedback and feedforward loops (*16, 17*), IFNs promote expression of ISGs to enhance the innate immune response to viral RNA.

When viral infection progresses in a gradual manner, the cytokine alert signal reaches most of the cells before they come into contact with the proliferating virus. This physiologically realistic scenario, recapitulated in vitro in experiments on respiratory epithelial cells challenged with a virus at a low multiplicity of infection (MOI), gives rise to three distinct subpopulations of cells: (*i*) primary infected cells that may produce IFNβ, (*ii*) not-yet-infected cells that respond to IFNβ, and (*iii*) IFNβ-primed cells that become infected due to infection spread. In Fig. 1, A and B, we show how infections with the human respiratory syncytial virus (RSV) or influenza A virus (IAV) activate the IRF3 pathway (leading to secretion of IFNs) and eventually activate (principally, in the paracrine manner) STAT1. In fig. S1A we show that activation of STAT1 and STAT2 leads to accumulation of ISGs (OAS3, PKR and RIG-I) as well as STAT1 and STAT2. Ultimately, infection of cells from the IFNβ-primed subpopulation leads to termination of STAT1 activity in these cells and activation of the IRF3 pathway (see late hours in Fig. 1, A and B, and the scheme in Fig. 1C). This is supported by population-based data showing a decrease of the p-STAT1-to-RSV fusion glycoprotein ratio as well as the p-STAT1-to-IAV nucleoprotein ratio between 24 and 48 h of infection (fig. S1B). We consider the behavior of the IFN-primed subpopulation of cells to be of crucial importance for shaping the kinetics of progression of viral infection (*18*).

**Figure 1.**
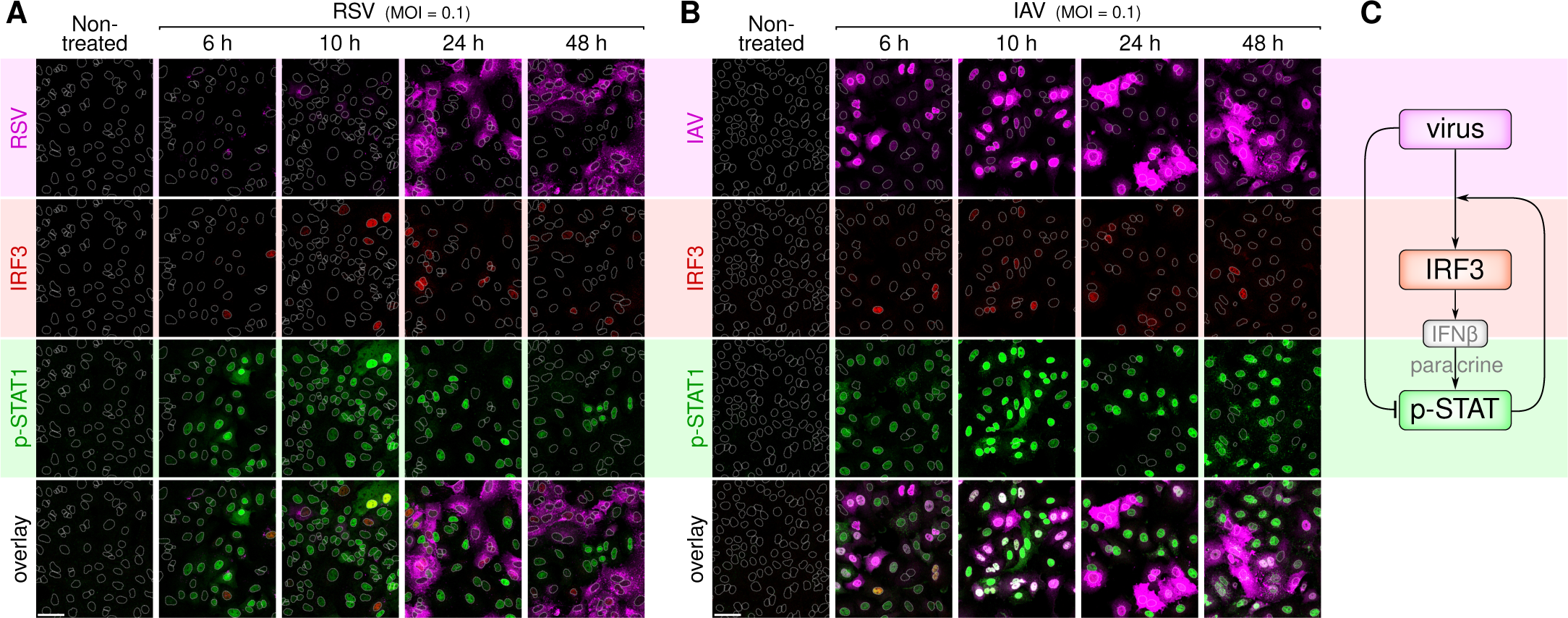
Progression of two respiratory virus infections in a monolayer of A549 cells. **A** Cells infected with RSV at MOI = 0.01 stained for RSV, IRF3, and p-STAT1 in indicated time points. **B** Cells infected with IAV at MOI = 0.1 stained for IAV, IRF3, and p-STAT1 in indicated time points. **C** Scheme of virus-initiated STAT activation (elicited by paracrine IFNβ signaling) and STAT inhibition in infected cells. Panels A, B show representative excerpts from confocal images of 3 independent experiments. Scale bar, 50 μm. Nuclear outlines (white dashed contours) were drawn based on DAPI staining (not shown). See Source Data for corresponding uncropped full-resolution immunostaining images.

Motivated by the role of priming with IFNβ, we investigated the activation of the NF-κB/IRF3 pathways by an analog of viral RNA, poly(I:C), as well as activation of the STAT1/STAT2 pathway by IFNβ. Using an alveolar epithelial cell line A549 and bronchial epithelial cell line BEAS-2B, we found that in IFNβ-stimulated cells, poly(I:C) terminates STAT signaling. Deactivation of STAT1/2 turned out to result from the depletion of the IFNβ receptor, IFNAR. This in turn results from both translation inhibition by PKR-phosphorylated eIF2α and IFNAR1 transcript degradation by RNase L. We also found that RNase L rapidly degrades STAT-regulated transcripts (including those of RIG-I, PKR, and OAS1/2/3), but does not affect transcripts of IFNβ and weakly impacts transcripts of interleukin-6 and -8, permitting propagation of paracrine signaling. Although poly(I:C) is ubiquitously used as an analog of viral RNA (*19*), one can expect that viruses, employing their non-structural proteins, interact in a more complex way with the cells they infect.

The innate immune response arises in the crosstalk of the NF-κB/IRF3 pathways and the STAT1/STAT2 pathway. Although these pathways working separately are well understood (*20–22*), a comprehensive mathematical model accounting for their interactions is still missing. Building such a model is challenging because of numerous feedback loops controlling activity of considered transcription factors and coupling the regulatory modules. For example, activity of NF-κB is controlled by NF-κB-inducible IκBα (*23*); another NF-κB-inducible inhibitor, A20, attenuates activation of both NF-κB and IRF3 (*24–26*); STAT signaling is known to be confined by proteins from the SOCS and the PIAS family (*27*) as well as USP18 (formerly UBP43) (*28*). SOCS1, SOCS3, and USP18 are of importance because they are transcriptionally regulated by STATs and thus mediate several negative feedbacks (*28, 29*). Expression of these three inhibitory proteins depends on the cell line; they are mainly expressed in immune cells (*29, 30*), additionally SOCS1 and SOCS3 are frequently silenced by hypermethylation in cancers (*31, 32*). Another challenge arises because of a large number of free parameters, that make even the models of submodules unidentifiable (*33, 34*). As the structural identifiability can be nearly algorithmically removed by reparameterization, the practical identifiability is more challenging (*35, 36*). To reach practical parameter identifiability we coarse-grained the model and performed a comprehensive set of experiments using various temporal stimulation protocols on A549 WT and PKR KO, RNase L KO, and RNase L & PKR double KO cells.

In our previous study (*17*), we dissected the NF-κB/IRF3–STAT1/STAT2 crosstalk to understand the role of priming with IFNβ. Here, based on our new experimental findings, we developed a computational model that explains how poly(I:C) converts IFNβ-responding cells into IFNβ-secreting cells, which are executing a distinct transcriptional program. As a consequence, the pairs of NF-κB/IRF3 and STAT1/STAT2 transcription factors are unlikely to be active simultaneously in the same cell.

## RESULTS

### Poly(I:C) activates NF-κB and IRF3 and terminates activity of STAT1/2

To reproduce and characterize the bulk behavior of various cell subpopulations in an infected respiratory epithelium, we treated A549 cells according to three protocols. (1) Cells that were warned about nearby infection through paracrine signaling and thus were granted time to upregulate ISG-coded proteins, achieving an antiviral state, were imitated using 24 h-long stimulation with IFNβ. (2) Cells in the antiviral state that have been challenged with a virus and are responding to non-self RNA were imitated by 24 h-long prestimulation with IFNβ followed by addition of poly(I:C) (without replacement of IFNβ-containing cell culture medium; we will refer to this protocol as the ‘IFNβ + poly(I:C) protocol’). (3) Finally, cells infected by an RNA virus and responding to viral RNA (without interference from viral proteins) were imitated by 10 h-long treatment with poly(I:C). The response to the third protocol is the most complex as a direct effect of poly(I:C) is convoluted with the response to secreted IFNβ and only with the help of the first and the second protocol are we able to distinguish individual contributions of these two factors. In the second protocol, the effect of IFNβ secreted upon stimulation with poly(I:C) can be neglected, because after 24 h of stimulation with IFNβ, the addition of fresh IFNβ does not influence phosphorylation of STAT1 and STAT2 (see fig. S2A).

Stimulation with IFNβ at 1000 U/ml leads to rapid activation of STAT1 and STAT2, which peaks at 15–30 min (fig. S2B). STAT1/2 phosphorylation measured at 30 min, 2 h, and 4 h increases gradually for IFNβ concentrations spanning from 30 to 1000 U/ml (fig. S2C). STAT1/2 phosphorylation lasts more than 30 h and after 24 h we observe accumulation of antiviral proteins — PKR, RIG-I, OAS1/2/3 — as well as STAT1 and STAT2 (Fig. 2, A and B). Since PKR, RIG-I, and OAS1/2/3 serve as cytoplasmic non-self RNA sensors, their upregulation sensitizes cells to poly(I:C) (*17*).

**Figure 2.**
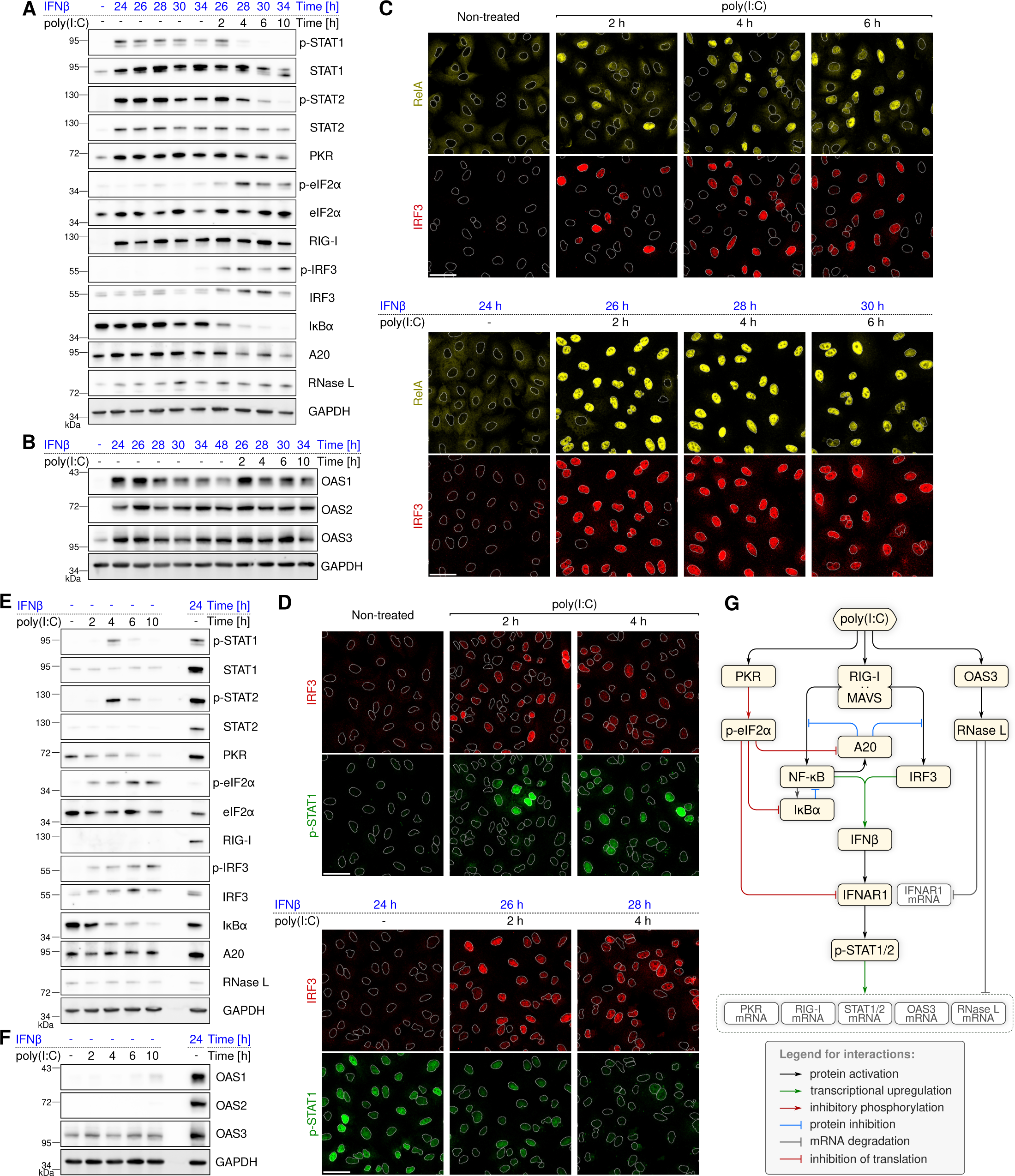
Poly(I:C) triggers and terminates STAT1 and STAT2 activation. **A, B** A549 WT cells stimulated for 34 h with IFNβ (1000 U/ml) and with poly(I:C) (0.1 μg/ml) added 24 h after beginning of stimulation with IFNβ. **C** Activation and nuclear translocation of NF-κB and IRF3 in response to poly(I:C) (0.1 μg/ml) with or without 24 h-long priming with IFNβ (1000 U/ml). A549 WT cells were stained for RelA (an NF-κB subunit) and IRF3 in indicated time points. **D** Activation and nuclear translocation of STAT1 and IRF3 in response to poly(I:C) (0.1 μg/ml) with or without priming with IFNβ (1000 U/ml). A549 WT cells were stained for p-STAT1 and IRF3 in indicated time points. **E, F** A549 WT cells stimulated with either poly(I:C) (0.1 μg/ml) for 10 h or IFNβ (1000 U/ml) for 24 h. **H** Simplified diagram of innate immune signaling in response to poly(I:C) and IFNβ. Panels A, B and E, F show representative blots from 2 independent experiments. In fig. S4, all blots from panels A, E and blots for OAS3 from panels B, F are quantified and juxtaposed with computational model predictions. Panels C, D show representative excerpts from confocal images of 3 independent experiments. In all fields of view and channels, nuclear outlines (white dashed contours) were drawn based on DAPI staining (not shown); scale bar, 50 μm. See Source Data for corresponding uncropped blots and full-resolution immunostaining images.

Using the IFNβ + poly(I:C) protocol we observed that interferon-induced phosphorylation of STAT1 and STAT2 is severely suppressed within 4 h after addition of poly(I:C), although total levels of these proteins remain relatively unchanged (Fig. 2, A and B). Simultaneously, addition of poly(I:C) induces rapid activation of IRF3 (by phosphorylation of Ser^396^) and extensive degradation of the principal NF-κB inhibitor, IκBα. This indicates that poly(I:C) abruptly converts IFNβ-responding cells, with the transcriptional program governed by STAT1 and STAT2, into IFNβ-producing cells, in which the transcriptional program is governed by IRF3 and NF-κB.

Immunofluorescence staining revealed that in A549 cells, stimulation with poly(I:C) within 2–4 h leads to activation and nuclear translocation of transcription factors IRF3 and NF-κB, although only a fraction of cells respond with a joint NF-κB and IRF3 activation (Fig. 2C). These cells synthesize and secrete IFNβ. Despite the presence of IFNβ secreted and accumulated over the course of the 24 h-long poly(I:C) treatment (see ELISA in fig. S3), phosphorylation and nuclear translocation of STAT1 is observed only briefly at 4 h (Fig. 2, D and E) and does not lead to accumulation of antiviral proteins (Fig. 2, E and F), contrary to what we observed in the case of direct IFNβ stimulation. As evidenced by staining of RelA (a subunit of NF-κB) and IRF3 in the nucleus, poly(I:C) treatment leads to a more robust activation of the NF-κB and IRF3 pathways in IFNβ-prestimulated cells (Fig. 2C). Immunostaining of IRF3 and phosphorylated STAT1 (p-STAT1) confirms that in the case of IFNβ-prestimulated cells, poly(I:C) activates IRF3 and simultaneously terminates phosphorylation of STAT1 (Fig. 2D).

From now on we will experimentally dissect the molecular basis of the observed cell state switch to support the mathematical model of the innate immune response to IFNβ and poly(I:C). The model is described in detail in a further Results subsection but, as we begin juxtaposing experimental results and predictions of the model, in the following enumeration we highlight crucial regulatory processes contained therein (see Fig. 2G):

(1) Poly(I:C) forms complexes with RIG-I and MAVS to activate transcription factors NF-κB and IRF3. Joint activation and nuclear translocation of NF-κB and IRF3 leads to transcription and translation of IFNβ. NF-κB also triggers transcription of its inhibitors, IκBα and A20, the latter being an inhibitor of both NF-κB and IRF3.
(2) Poly(I:C) activates PKR, which in turn inactivates eIF2α by phosphorylation. In the model, p-eIF2α acts as an inhibitor of translation of IFNAR1 and NF-κB-inducible inhibitors: IκBα and A20.
(3) Poly(I:C) activates OAS3 that activates RNase L, which in turn degrades transcripts of IFNAR1, IκBα and A20, STAT1, STAT2, and STAT-regulated genes.
(4) IFNβ activates IFNAR1 (in an autocrine and paracrine manner), which in turn enables phosphorylation of STAT1 and STAT2, allowing them to dimerize and translocate to the nucleus. Heterodimers of p-STAT1/2 serve as transcription factors of RIG-I, PKR, OAS3, as well as STAT1 and STAT2.

We confront simulated model trajectories (fig. S4) with quantified blots shown in Fig. 2, A, B, E and F, and their replicates. As demonstrated, the model satisfactorily reproduces profiles of protein activation and accumulation in response to stimulation with IFNβ and/or poly(I:C).

### Termination of STAT activity is caused by depletion of IFNAR

To explain the mechanism responsible for the observed termination of STAT1/2 activity upon treatment with poly(I:C), we turned our attention to the receptor of IFNβ, IFNAR. Although the JAK/STAT pathway is regulated on several levels (*37, 38*), the processes that control the abundance of IFNAR subunits, such as endocytosis, trafficking, and degradation, have been proposed to be of key importance in tuning interferon signaling (*39*). To investigate the dynamics of depletion of IFNAR1 (a subunit of IFNAR) and dephosphorylation of STAT1/2, we performed an experiment with two IFNβ pulses, the first of duration of 2 h and the second, occurring after a 2 h or 4 h-long break, lasting 30 min (Fig. 3A). IFNβ stimulation leads to a rapid depletion of IFNAR1 (observed at 30 min) followed by a decrease of STAT1 phosphorylation at 2 h. IFNβ withdrawal also leads, within 1 h, to a nearly complete termination of both STAT1 and STAT2 phosphorylation, but in this case IFNAR1 is replenished within 4 h. After 4 h (but not after 2 h) since IFNβ withdrawal, the level of IFNAR1 seems to be restored sufficiently to allow equally high STAT1 and even higher STAT2 phosphorylation in response to the second IFNβ pulse. The surge of p-STAT2 likely stems from the accumulation of STAT2 protein after the first IFNβ pulse.

**Figure 3.**
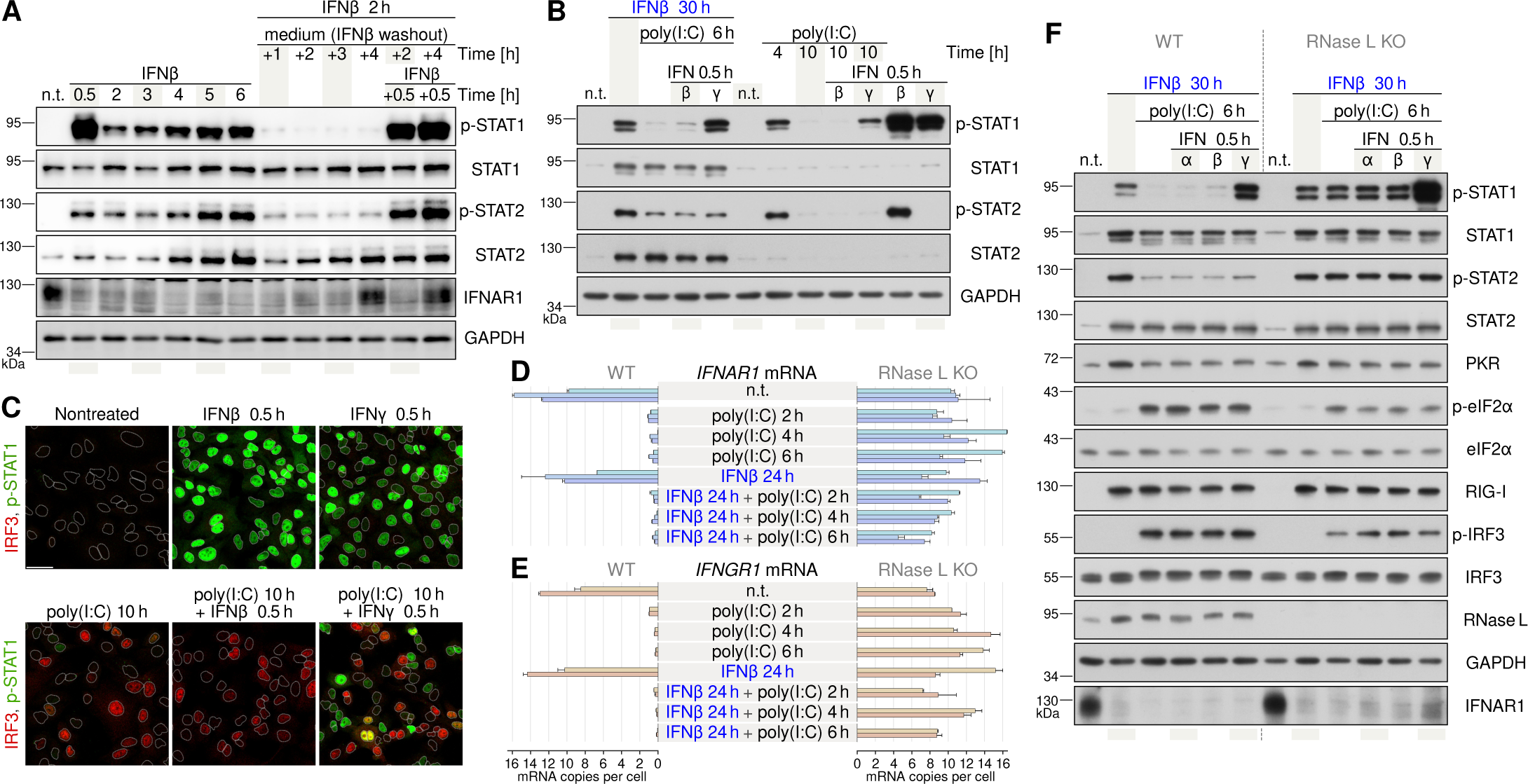
Termination of STAT activity follows RNase L-dependent depletion of IFNAR. **A** A549 WT cells were stimulated with IFNβ (1000 U/ml) according to the indicated protocols. For example, in the last column, cells were stimulated with IFNβ for 2 h, then IFNβ was washed out by medium replacement and cells were cultured in the fresh medium for subsequent 4 h, and finally IFNβ was added for 0.5 h (protocol total duration is 6.5 h). Representative blots from 3 independent experiments are shown. Quantified blots from panel A are confronted with computational model predictions in fig. S6A. **B** A549 WT cells were stimulated with IFNβ (1000 U/ml), IFNγ (20 ng/ml), and poly(I:C) (0.1 μg/ml) according to indicated protocols. For example, in the fifth column, the experiment was terminated after 30 h of IFNβ stimulation with poly(I:C) added 6 h and IFNγ added 0.5 h before cell lysis and protein extraction. Representative blots from 2 independent experiments are shown. **C** A549 WT cells were stimulated either with IFNβ (1000 U/ml) or IFNγ (20 ng/ml) for 30 min either with or without poly(I:C) (0.1 μg/ml) prestimulation. Scale bar, 50 μm. Nuclear outlines (white dashed contours) were drawn based on DAPI staining (not shown). **D, E** Levels of *IFNAR1* and *IFNGR1* transcripts measured with digital PCR in A549 WT and RNase L KO cells after poly(I:C) (1 μg/ml), either with or without 24 h-long prestimulation with IFNβ (1000 U/ml). Error bars represent s.e.m. of two technical replicates in each experiment. **F** A549 WT and RNase L KO cells were stimulated with IFNβ (1000 U/ml), IFNγ (20 ng/ml), IFNα (200 U/ml) and poly(I:C) (0.1 μg/ml) using indicated protocols. Representative blots from 3 independent experiments are shown. See Source Data for corresponding uncropped blots and full-resolution immunostaining images.

The 6 h-long stimulation with poly(I:C) preceded by a 24 h-long prestimulation with IFNβ leads to inhibition of STAT1/2 phosphorylation as evidenced by Western blotting (Fig. 3B). To corroborate this result, we expanded this protocol with a subsequent 30-min stimulation with interferon β or γ. As shown by Western blotting, additional stimulation with interferon β fails to restore phosphorylation of either STAT1 or STAT2 (Fig. 3B). Interferon γ, that activates STAT1 (but not STAT2) using a different receptor, IFNGR, is nonetheless able to restore phosphorylation of STAT1. This suggests that the JAK/STAT pathway downstream of the receptor retains its signaling capabilities. Altogether, these results indicate that the observed termination of STAT activity is tied specifically to depletion of IFNAR and not to inhibition of the downstream JAK/STAT pathway by USP18 or proteins from the SOCS or the PIAS families. Termination of STAT1 and STAT2 activity after poly(I:C) is also observed in MAVS KO cells (fig. S5), which respond to IFNβ but are not able to activate IRF3 and secrete IFNβ in response to poly(I:C) (fig. S3). This implies that depletion of IFNAR and termination of STAT activity are also not associated with MAVS signaling.

We observed a similar effect even without initial prestimulation with IFNβ. In cells treated with poly(I:C) only, STAT1 and STAT2 become briefly phosphorylated at 4 h post-treatment, but completely dephosphorylated at 10 h post-treatment. Additional stimulation with IFNβ is, again, not able to restore STAT1 and STAT2 activation, whereas stimulation with IFNγ activates STAT1, however to a much lower degree than in non-prestimulated cells (Fig. 3B). Consistently, immunostaining shows that IFNγ administered after 10 h of stimulation with poly(I:C) activates STAT1 in a smaller fraction of cells than IFNγ acting on nontreated cells (Fig. 3C). This may suggest that the level of IFNGR is also reduced during poly(I:C) stimulation, however not to the same extent as the level of IFNAR.

### RNase L and PKR are responsible for depletion of IFNAR after poly(I:C) stimulation

To assess whether depletion of IFNAR and IFNGR is accompanied by downregulation of expression of their key subunits, we evaluated the role of two antiviral proteins, PKR and RNase L, as their modes of action (respectively, inhibition of translation and degradation of RNA after recognition of non-self RNA) makes them likely suspects for downregulators of protein levels under conditions that simulate viral infection. To this end, we used A549 cell lines with genomic deletions of either *EIF2AK2* (PKR KO) or *RNASEL* (RNase L KO), or both these genes simultaneously (RNase L & PKR double KO, dKO).

Using digital PCR, we measured mRNA levels of *IFNAR1* and *IFNGR1* at the end of the above-described stimulation protocols: IFNβ, poly(I:C), and IFNβ + poly(I:C). IFNβ stimulation alone does not affect *IFNAR1* mRNA or *IFNGR1* mRNA (Fig. 3, D and E). Stimulation with poly(I:C), however, in WT cells leads to about 10-fold reduction of *IFNAR1* and *IFNGR1* mRNAs after 2 h, whereas depletion of these transcripts is not observed in RNase L KO cells. This suggests that the degradation of transcripts for both receptors by poly(I:C)-activated RNase L is responsible for the lack or lowered STAT1 activation in response to IFNβ or IFNγ following poly(I:C) treatment (Fig. 3B). In agreement with this, inhibition of phosphorylation of STAT1 and STAT2 after 6 h of poly(I:C) treatment of IFNβ-primed cells is not observed in RNase L KO cells (Fig. 3F). In the RNase L KO cell line, IFNAR1, although substantially reduced after IFNβ stimulation, remains at the level that allows continued phosphorylation of STAT1 and STAT2 (Fig. 3F). In fig. S6 we confront quantifications of blots shown in Fig. 3A (and their replicates) as well as digital PCR profiles shown in Fig. 3D with simulated model trajectories.

Next, we compare profiles of STAT1/2 phosphorylation in response to poly(I:C) in IFNβ-primed WT, RNase L KO, PKR KO and dKO cells (Fig. 4 and fig. S7). We may observe that in comparison to WT cells, the RNase L and PKR knockouts cooperatively extend phosphorylation of STAT1/2. In Fig. 4B we confront quantifications of blots shown in Fig. 4A with model predictions, while in fig. S7 we present a juxtaposition of the four A549 cell lines in selected time points (from Fig. 4A) on single blots (and likewise confront blot quantifications with model predictions). Double KO cells exhibit sustained STAT1/2 activation with some decrease observed at 10 h of poly(I:C) stimulation. This decrease is not reproduced by the model and may be a consequence of feedbacks mediated by USP18 (*28*) and SOCS1/3 (*29*) not included in the model.

**Figure 4.**
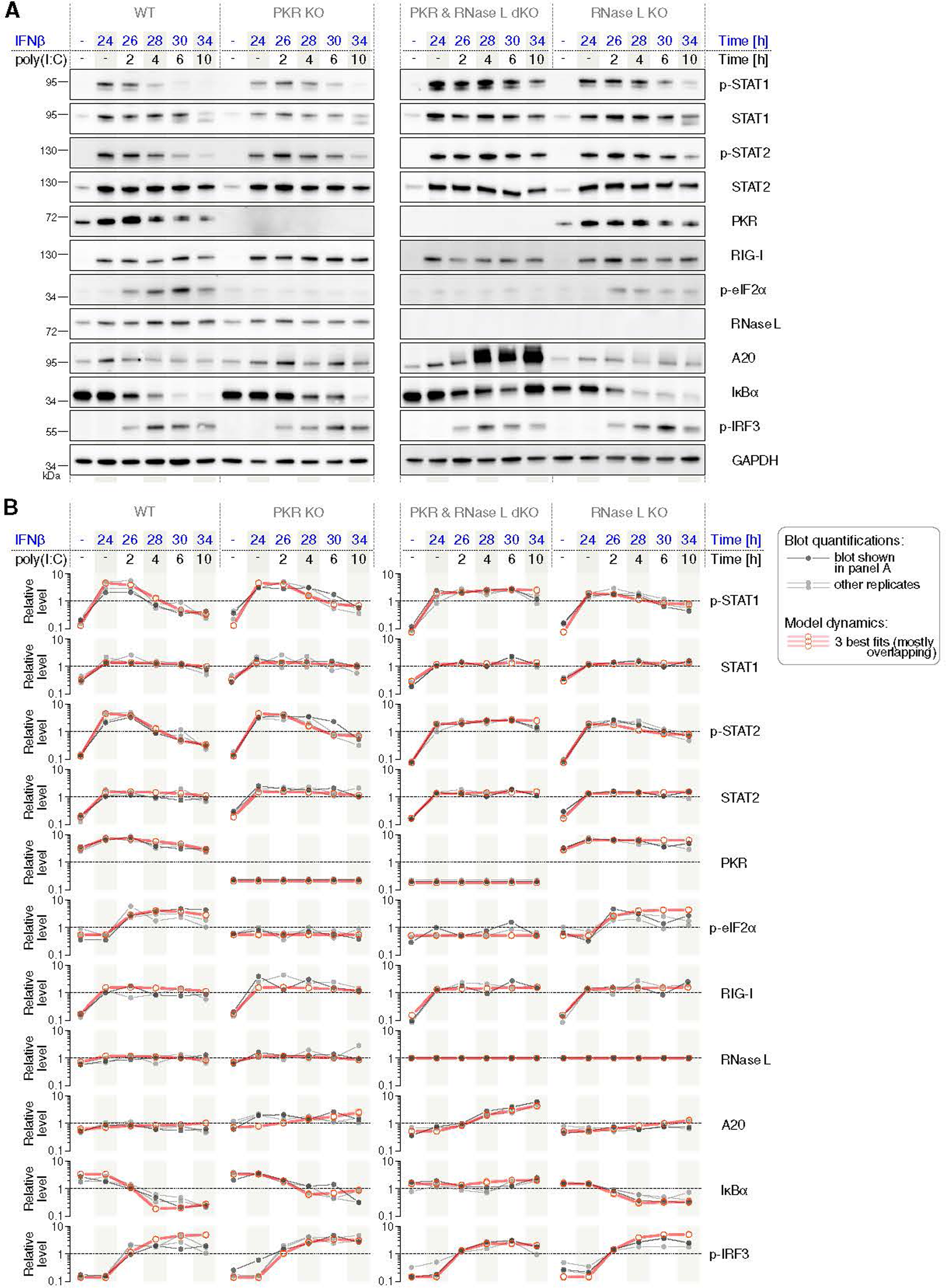
RNase L and PKR contribute to termination of STAT activity. **A** A549 WT, PKR KO, PKR & RNase L double KO, and RNase L KO cells prestimulated for 24 h with IFNβ (1000 U/ml), then costimulated with poly(I:C) (0.1 μg/ml). Representative blots from 3 independent experiments are shown. **B** Quantified Western blots juxtaposed with numerical solutions (continuous in time but, for the sake of comparison to experimental data, extracted in experimental time points and connected with straight line segments to guide the eye). See Source Data for corresponding uncropped blots.

Prompted by an anonymous Reviewer, we investigate whether the effect of termination of STAT1 activity by poly(I:C) holds also for IFNγ-mediated signaling. In an additional experiment performed for A549 WT and dKO cells we observe (as expected) that the simultaneous stimulation of WT cells with IFNβ and poly(I:C) after 6 h leads to a nearly full depletion of IFNAR1 and suppression of STAT1 phosphorylation (fig. S8A). Analogously, the simultaneous stimulation of WT cells with IFNγ and poly(I:C) leads to nearly full depletion of IFNGR1 and, correspondingly, suppression of STAT1 phosphorylation (fig. S8B). We should notice that although IFNβ alone leads to depletion of IFNAR1, IFNγ only jointly with poly(I:C) causes depletion of IFNGR1, showing that for termination of IFNγ-mediated signaling, poly(I:C) is even more critical.

Next, we demonstrate that depletion of IFNAR1 or IFNGR1 in response to joint stimulation with their respective interferon and poly(I:C), and subsequent suppression of STAT1 phosphorylation take place also in another respiratory (but non-cancerous) cell line, BEAS-2B (fig. S9). The difference between these two lines is such that, in contrast to A549 cells, the stimulation with IFNβ alone in BEAS-2B cells leads only to a modest decrease of IFNAR1, suggesting an initially higher level of the receptor. Of note, IFNβ exerts some antiproliferative effect on both A549 and BEAS-2B cells; however, since this effect is similar for A549 WT, PKR KO, RNase L KO and dKO cells (fig. S10), we do not expect it to influence our results.

Finally, we confirm (within manuscript revision) that the impact of PKR KO, RNase L KO and dKO on STAT1/2 phosphorylation also holds in cells that were not prestimulated with IFNβ/γ (and thus did not develop an antiviral state prior to poly(I:C) treatment). In contrast to WT and PKR KO cells, the RNase L KO cells and dKO cells display nearly sustained phosphorylation of STAT1/2 at least up to 8 h since poly(I:C) treatment (fig. S11).

In summary, we showed that poly(I:C) stimulation terminates IFNβ-induced phosphorylation of STAT1/2 by depletion of IFNAR caused by *IFNAR1* mRNA degradation by RNase L and PKR-regulated inhibition of translation. These two processes also diminish the level of IFNGR1 and in this way reduce STAT1 responsiveness to IFNγ. The effect is observed both in cancerous A549 and non-cancerous BEAS-2B cells.

### Poly(I:C)-activated RNase L and PKR downregulate NF-κB and IRF3 inhibitors IκBα and A20

Nuclear NF-κB translocation is enabled by degradation of its inhibitor, IκBα. In resting cells, IκBα sequesters NF-κB in an inactive form in the cytoplasm (*23*). However, despite continued NF-κB activation by poly(I:C), IκBα (being transcriptionally NF-κB-inducible) is not rebuilt in WT cells (Fig. 4). Also, the level of another NF-κB- and IRF3-inducible protein, A20, does not increase (Fig. 4). Regulation of IκBα and A20 in response to poly(I:C) is thus different than observed after TNFα stimulation, when rapid IκBα degradation and NF-κB nuclear translocation are followed by IκBα resynthesis (within 2 h after TNFα stimulation) and a significant increase of A20 (fig. S12). This may suggest that IκBα and A20 are downregulated by poly(I:C)-activated RNase L and PKR. Indeed, in RNase L & PKR dKO cells responding to poly(I:C) IκBα is rebuilt and A20 increases significantly. As a consequence of the higher level of the IRF3 inhibitor, A20, the level of IRF3 phosphorylation in the dKO cell line is lower than in the WT or single KO cell lines (Fig. 4 and fig. S7).

### RNase L differentially regulates transcripts of NF-κB/IRF3- and STAT1/2-inducible genes

Using RT-PCR we measured mRNA levels of both the NF-κB/IRF3- and the STAT1/2-regulated genes, after IFNβ and IFNβ + poly(I:C). Poly(I:C) directly activates NF-κB/IRF3, triggering the production of IFNβ and subsequent transcriptional induction of interferon-dependent genes, rendering the direct effect of poly(I:C) on levels of transcripts hard to resolve. Within the IFNβ + poly(I:C) protocol, however, in the first phase (stimulation with IFNβ) we can observe the accumulation of transcripts of ISGs, whereas in the second phase (treatment with poly(I:C)), we can observe the accumulation of transcripts of NF-κB- and IRF3-inducible genes (Fig. 5A).

**Figure 5.**
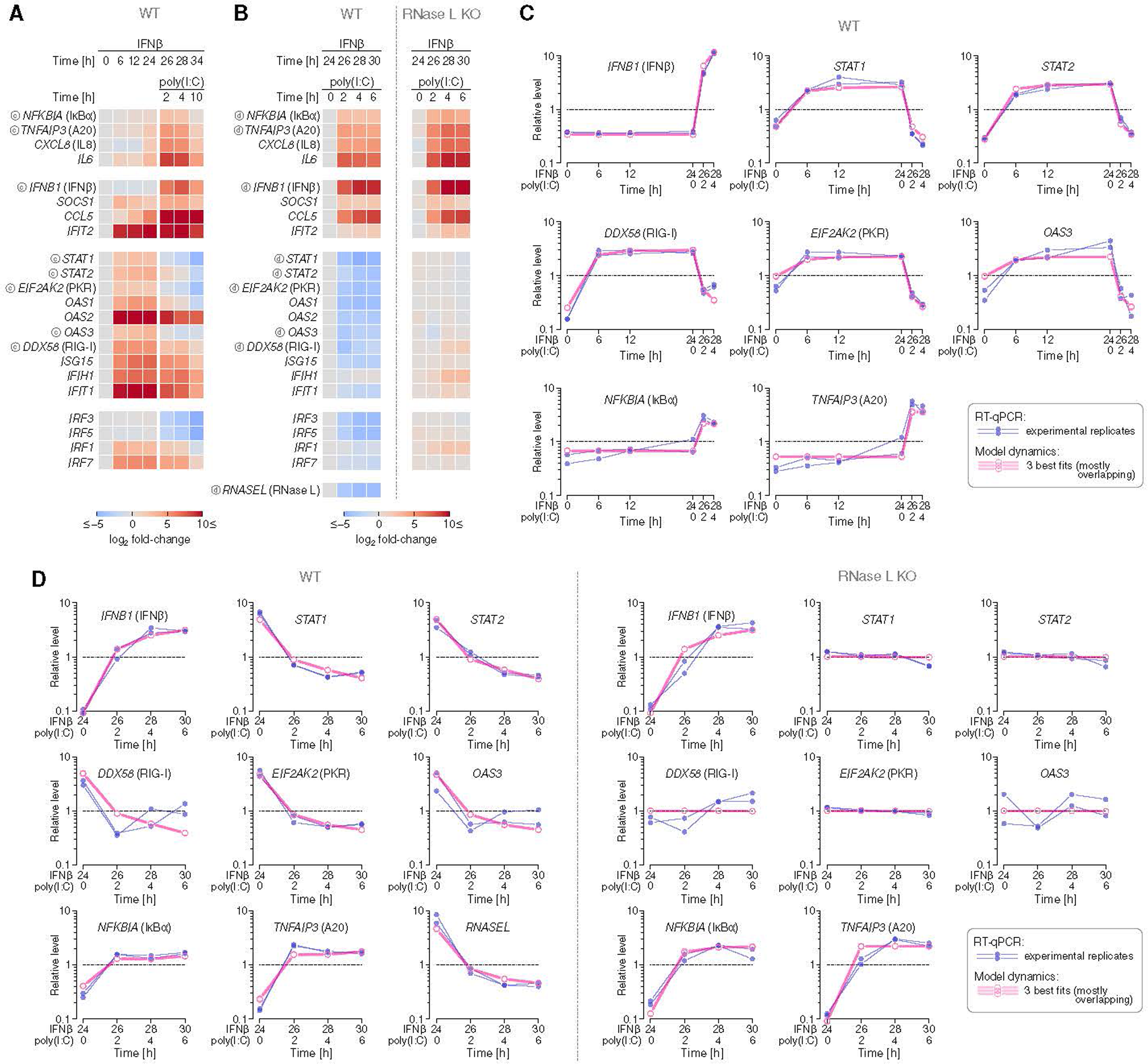
Poly(I:C) and IFNβ trigger distinct transcriptional programs. **A** Dynamics of transcripts (relative to a control) in A549 WT cells stimulated with IFNβ (1000 U/ml) for 24 h then additionally with poly(I:C) (1 μg/ml). Heatmaps show fold-change in RT-qPCR using the logarithmic scale with base 2. Encircled letter ‘c’ indicates transcripts included in the computational model, shown in panel C. **B** Dynamics of transcripts (relative to a control) in A549 WT and A549 RNase L KO cells stimulated with IFNβ (1000 U/ml) for 24 h and then additionally with poly(I:C) (1 μg/ml). Heatmaps show fold-change in RT-qPCR using the logarithmic scale with base 2. Encircled letter ‘d’ indicates transcripts included in the computational model, shown in panel D. **C** Selected RT-qPCR profiles from panel A juxtaposed with model trajectories (continuous in time but, for the sake of comparison to experimental data, extracted in experimental time points) and connected with straight line segments to guide the eye). **D** Selected RT-qPCR profiles from panel B juxtaposed with model trajectories (continuous in time but, for the sake of comparison to experimental data, extracted in experimental time points and connected with straight line segments to guide the eye).

In the course of 24 h-long IFNβ stimulation, we observed an increase of mRNAs of all measured ISGs, including PKR, RIG-I, OAS3, as well as mRNAs of STAT1 and STAT2, which explains the post-IFNβ accumulation of these proteins observed earlier (Fig 2, A and B). The highest fold increase is observed for IFIT2, OAS2, ISG15, IFIH1, IFIT1, and IRF7. After a subsequent poly(I:C) stimulation, we observe the accumulation of transcripts of NF-κB- and IRF3-inducible genes (of IκBα, A20, IL6, IL8, CCL5, IFNβ), featured by a rapid drop of the levels of all measured STAT-induced transcripts. To explain this effect, we compared responses of IFNβ-primed WT and RNase L KO cells to 6 h-long stimulation with poly(I:C) (Fig. 5B). In RNase L-KO cells, the levels of STAT-inducible genes remain almost unaffected, whereas in WT cells these levels decrease sharply.

In the IFNβ-primed cells, after stimulation with poly(I:C) the level of mRNA for IFNβ increases with the same kinetics in both WT and RNase L KO cells (Fig. 5B, see also (*40*)). The increase of mRNAs of NF-κB-inducible interleukin-6 and -8 is only slightly lower in WT than in RNase L KO cells (Fig. 5B). Expression of A20 and IκBα is however attenuated by RNase L, as evidenced by their level being 3–4 times higher in RNase L KO than WT cells in response to poly(I:C). The latter result is in agreement with resynthesis and accumulation of IκBα and A20 proteins in RNase L & PKR dKO cells after poly(I:C), that we observed previously (Fig. 4). As corroborated by the mathematical model, the relatively modest difference in expression of IκBα and A20 genes between WT and RNase L KO cells is caused by the negative feedback coupling IκBα and A20 with NF-κB. In WT cells, by degrading mRNAs for IκBα and A20, RNase L promotes NF-κB activation, which in turn induces the synthesis of mRNAs for IκBα and A20.

### RNase L and PKR play contradicting roles in regulation of the innate immune response

As demonstrated in Fig. 4, poly(I:C)-activated PKR and RNase L may promote NF-κB and IRF3 signaling and thus induce synthesis of mRNA for IFNβ by downregulation of inhibitors IκBα and A20. It requires more detailed investigations to determine whether this mechanism compensates for degradation of mRNA for IFNβ by RNase L, or whether IFNβ transcripts are resistant to RNase L-catalyzed cleavage. The latter, simplifying assumption is sufficient to reproduce profiles of mRNA for IFNβ mRNA using the mathematical model (Fig. 5C). We should notice that after poly(I:C) stimulation, IFNβ concentrations measured by ELISA are about fivefold higher in the case of PKR- or RNase L-deficient cells and more than fivefold higher in dKO cells relative to WT cells (fig. S13). This shows that RNase L is still involved in regulation of IFNβ expression in an indirect way, possibly by breaking down most cellular RNAs (including ribosomal RNAs 16S and 23S, see fig. S14 and Ref. *41*). Activation of PKR, which inhibits translation through eIF2α phosphorylation, also downregulates production of IFNβ, although it is not clear whether it acts in a direct or indirect manner. Dalet *et al.* (*42*) postulated that dephosphorylation of EIF2α by GADD34 leads to waves of IFNβ synthesis in single cells. As global effects of RNase L and PKR on cell functioning would require further analysis, such effects are not included in the computational model.

Despite the fact that activation of RNase L by poly(I:C) may promote long lasting activation of NF-κB and IRF3, it also downregulates secretion of IFNβ, terminates STAT1/2 activation induced by secreted IFNβ, and additionally prevents accumulation of ISG-encoded antiviral proteins — RIG-I, OAS1/2/3, PKR — and STAT1/2 by degradation of their transcripts. As already shown (Fig 2, E and F), the levels of these proteins remain nearly unchanged 10 h after treatment with poly(I:C) and are much lower than the levels reached after 24 h of stimulation with IFNβ.

### Computational model of the innate immune response

Based on our experimental findings, we constructed a computational model of the innate immune response to IFNβ and poly(I:C). The model consists of 5 interlinked regulatory modules (Fig. 6). A coarse view of the interactions (Fig. 6A) can be summarized as follows. Poly(I:C) activates ISG-encoded proteins: RIG-I, PKR, OAS3, RNase L. RIG-I triggers transcriptional activity of NF-κB and IRF3, which jointly trigger expression of IFNβ. IFNβ-activated STAT1/2 heterodimers promote synthesis of mRNAs of STAT1 and STAT2, creating a (topologically short) autoregulatory positive feedback loop. STAT1/2 heterodimers also promote synthesis of ISG-encoded proteins, giving rise to another, partially overlapping, positive feedback loop that couples all five modules. Both these positive feedback loops are undercut by poly(I:C)-activated ISGs. In particular, RNase L degrades transcripts of not only ISGs (that mediate the longer-range feedback loop), but also *IFNAR1* (disabling in this way STAT1/2 activation and consequently STAT1/2 accumulation), which ultimately results in full termination of STAT signaling.

**Figure 6.**
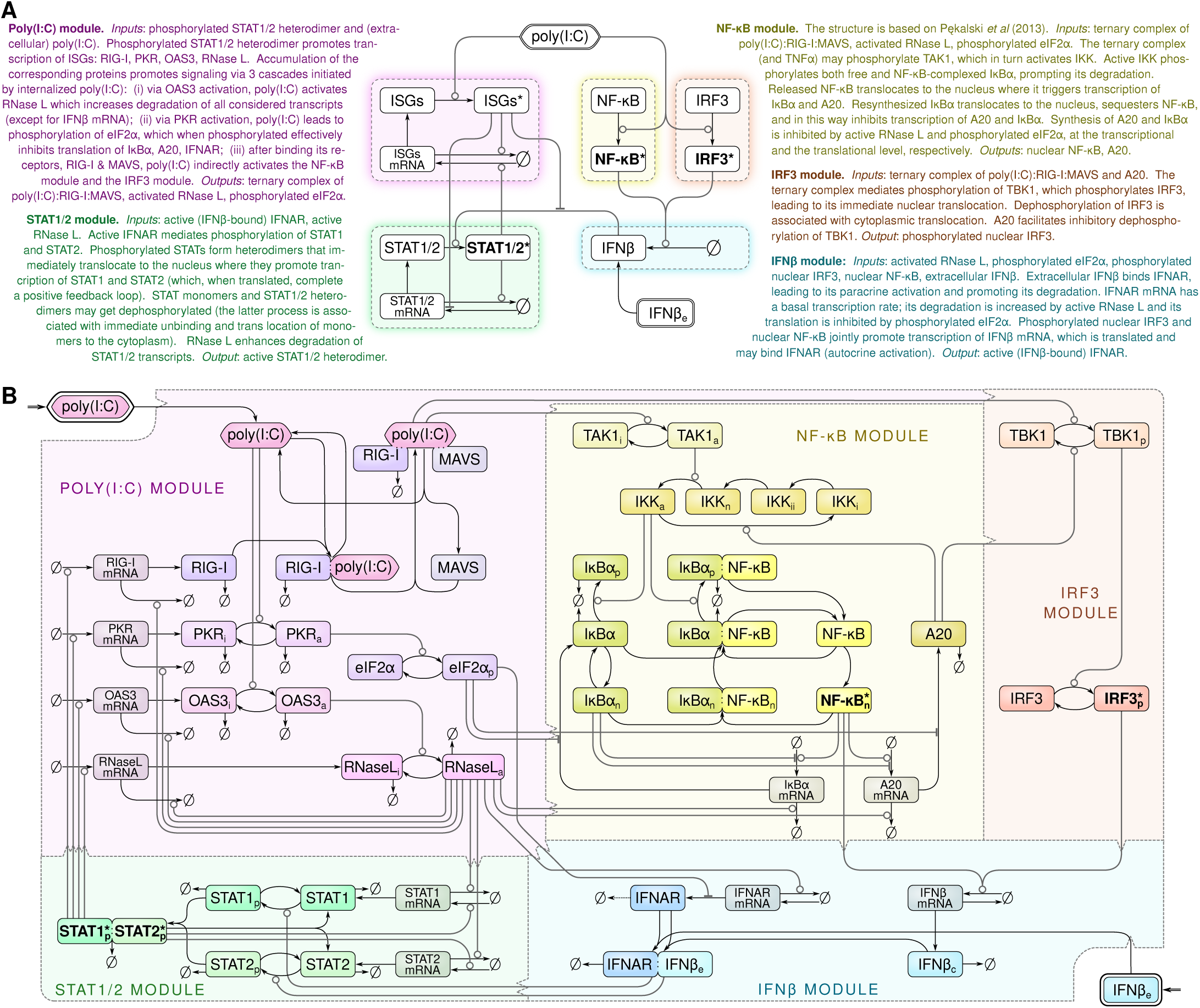
The computational model of the innate immune signaling. **A** Coarse-grained scheme of key interactions within and between five regulatory modules. Stars denote active forms of proteins. **B** Complete representation of all processes modeled within the five modules. The poly(I:C) module and the IFNβ module receive extracellular stimuli (marked with an additional thick outer border) and each module receives and generates signals from other modules (for example, the ternary complex of poly(I:C) : RIG-I : MAVS is an output from the poly(I:C) module and an input to both the NF-κB module and the IRF3 module). Lines corresponding to signals leaving and entering communicated modules are featured by directed notches on the dashed module border. Arrow-headed solid lines indicate transitions, circle-headed and hammer-headed solid lines indicate positive and negative influence, respectively. Bold font and asterisks indicate active transcription factors. Key for subscripts: e – extracellular, c – cytoplasmic, n – nuclear (neutral for IKK), p – phosphorylated, a – active, i – inactive, ii – inactive (secondary). Kinetic rate expressions and parameter values are given in Supplementary Table S1 (see Supplementary Materials).

The model has been formulated as a system of 53 ordinary differential equations generated from a system of chemical reactions defined in the BioNetGen language (*43*) (see Supplementary Materials). Our model enables the analysis of responses of wild-type and knock-out cells to IFNβ, poly(I:C), and TNFα; the latest is used to calibrate kinetic rate constants in the NF-κB pathway module, that follows Lipniacki *et al*. (*44*) and Tay *et al*. (*45*). To reach parameter identifiability, we reduced the model in two major steps: (1) nondimensionalization; (2) coarse-graining of model structure and parametrization. In the second step: (2a) we removed some intermediate steps in signal processing; (2b) to all kinetic rate constants which were not constrained from above, corresponding to relatively fast processes, we assigned an identical large value (equal to 1 s^−1^ for the sake of numerical stability); (2c) we assumed equal rate constants of processes of a given type that, based on our data, appear to have similar kinetics. In this way, we reduced the number of independent parameters to 38. We assessed parameter identifiability through linear analysis of the best log-fit by calculating singular values (which all turned out to exceed 0.31) and reported the ratio of the multiplicative parameter estimation error to the multiplicative experimental error (which turned out to be smaller than 2.06 for all parameters; see Table 1 and fig. S15, A and B). In fig. S16 we show convergence of all 38 model parameters for 5 best fits. For 10 best fits, deviations of log-fitted parameters have the mean of 0.05 and maximum of 0.19 for the worst-fitted parameter, which implies excellent parameter identifiability as for such a complex model. The model was constrained based on the experiments presented thus far as well as auxiliary experiments (figs. S17, S18 and S19). Overall, our comprehensive set of experimental data used for fitting model parameters comprised as many as 2915 data points obtained from various stimulation protocols and time points. The average discrepancy (multiplicative error) between the model and experimental data is 1.49 and is greater than the average error of experimental Western blot replicates equal 1.24, which likely results from simplifying assumptions and model coarse-graining. The main discrepancies are discussed at the end of this section. In revision, prompted by anonymous Reviewers, we performed additional experiments with the simultaneous IFNβ and poly(I:C) stimulation and with poly(I:C)-only stimulation and, without refitting the model, we used this experiment to verify model accuracy (fig. S8A, fig. S11B). The average multiplicative error between the model and data from these additional experiments was 1.34, which is below the average error obtained for experiments used to constrain the model (1.49). This confirms that the responses to the additional protocols were well predicted by the model.

**Table 1.** Summary of model complexity, data used for parameter fitting, and the fitted computational model.

| Property | Value |
| --- | --- |
| *Model complexity* |  |
| Number of variables in the model | 53 |
| Number of independent model parameters | 38 |
| *Statistics of data used for fitting* |  |
| Number of experimentally measured variables | 25 |
| Number of independent experimental data points used for parameter fitting<br>(dimensionality of the measurement vector space) | 2915 |
| Average multiplicative error between experimental replicates (for Western blots) | 1.24 |
| *Linear identifiability analysis* |  |
| Ratio of the largest parameter estimation error to the experimental error | 2.06 |
| The smallest singular value | 0.31 |
| *Parameter fitting* |  |
| Average multiplicative error between data and the best fit | 1.4845 |
| Average multiplicative error between data and the worst of 10 best fits | 1.4856 |
| Average pairwise error between 10 best fits | 1.02 |
| Max pairwise error between 10 best fits | 1.04 |
| Average s.d. of log-fitted parameters (from 10 best fits) | 0.05 |
| Maximum s.d. of log-fitted parameters (among 10 best fits) | 0.19 |

The main consequence of the discussed structure of interlinked feedbacks coupling the five modules is that a homogenous and a nonhomogenous population of cells respond to poly(I:C) in a qualitatively different manner. In agreement with the model, in a single cell or in a homogenous population of cells, poly(I:C) may elicit only transient STAT1/2 activity (Fig. 2E and fig. S4C). However, IFNβ secreted by poly(I:C)-stimulated cells may lead to long-lasting STAT1/2 activation and accumulation of ISG-encoded proteins in bystander cells without internalized poly(I:C) (Fig. 2A and fig. S4A). When these IFNβ-prestimulated cells are challenged by poly(I:C), they exhibit a rapid rewiring of signaling: termination of STAT1/2 activity (fig. S4A), degradation of ISG transcripts (Fig. 5C), activation of NF-κB and IRF3, and accumulation of NF-κB- and IRF3-induced transcripts and ISGs’ transcripts degradation (Fig. 5C).

As predicted by the model, rewiring of signaling in IFNβ-prestimulated cells responding to poly(I:C) is accomplished by RNase L and PKR. In the RNase L & PKR dKO cells, the decrease of STAT1/2 activity is much weaker than in WT or single-KO cells (Fig. 4 and fig. S7) and in RNase L-deficient cells, ISG transcripts are not degraded (Fig. 5C). As a consequence, the STAT1/2 transcriptional program is not interrupted in dKO cells. The model shows also that in the absence of RNase L and PKR, A20 (an inhibitor of both NF-κB and IRF3) may accumulate and IκBα (an inhibitor of NF-κB) is resynthesized, which attenuates NF-κB and IRF3 signaling (Fig. 4 and fig. S7).

The model implies that termination of STAT1/2 activity in IFNβ-prestimulated cells responding to poly(I:C) results from depletion of IFNAR, which is caused not only by degradation of *IFNAR1* mRNA by RNase L and inhibition of *IFNAR1* mRNA translation by PKR, but also by IFNAR endocytosis-driven decay (fig. S6). A consequence of the endocytosis of IFNAR upon stimulation with IFNβ is the observed significant peak of STAT1 phosphorylation (nonsignificant for STAT2) observed after about 30 min of IFNβ stimulation (fig. S2, A and B). IFNβ withdrawal results in a sharp decrease of STAT1/2 phosphorylation and enables recovery of IFNAR within 4 h (fig. S6A) and rapid STAT1/2 activation in response to another IFNβ pulse. Between 2 h and 24 h of IFNβ stimulation, STAT1/2 remain at about constant level, promoting accumulation of STAT1/2 proteins (fig. S2B). In agreement with the model, the accumulation of STAT2 protein results in significantly higher STAT2 (but not STAT1) activation in response to the second pulse of IFNβ (fig. S17). Since STAT monomers undergo faster dephosphorylation than STAT1/2 dimers, the initially lower level of STAT2 than STAT1 limits STAT2 phosphorylation level in non-prestimulated cells, and is in part responsible for the observed peak of STAT1 phosphorylation (fig. S2, A and B).

Finally, we should mention several discrepancies between model predictions and experimental data.

1. We observe some accumulation of transcripts for IκBα, A20, IL6 and IL8 (Fig. 5A) as well as A20 protein (Fig. 2E) in response to 24 h-long stimulation with IFNβ, which may suggest that IFNβ weakly activates NF-κB, in agreement with Ref. *46*. This regulation has not been directly confirmed in our system, and is thus not included in the model.
2. Although we observe rapid depletion of IFNAR1 due to the first pulse of IFNβ stimulation, this is not the case in response to the second pulse of IFNβ (Fig. 3A). This may suggest that a significant proportion of newly synthesized IFNAR1 remains in the cytoplasm (and thus is not depleted due to IFNβ stimulation). Because of the lack of sufficient data, this potential cytoplasmic retention of IFNAR1 is not included in the model.
3. We observe a significant depletion of PKR in non-prestimulated (and less significant in IFNβ-prestimulated) cells in response to poly(I:C), Fig. 2, E and A. We speculate that this effect may be caused by enhanced degradation of activated PKR. Since the depletion of PKR is more pronounced in non-prestimulated cells (in which the initial level of PKR is lower), we expect that this effect may be more pronounced at higher ratios of poly(I:C) to PKR.

## DISCUSSION

At the beginning of viral infection, a relatively small proportion of infected cells produce and secrete IFNβ (see Ref. *47* for ELISA after RSV and IAV), which activates the STAT pathway in an autocrine and paracrine manner (*10*). In infected cells, STAT activity appears to be transient, however in bystander not-yet-infected cells, IFNβ induces long-lasting STAT activity, enabling accumulation of antiviral proteins such as RIG-I, PKR, and OAS1/2/3 (*48*). Responses of the IFNβ-prestimulated (sub)population are crucial for containment of infection. Accordingly, we focused on responses to poly(I:C), an analog of viral RNA, to IFNβ, and to poly(I:C) after prestimulation with IFNβ. It should be noticed, however, that as viral RNA triggers the innate immune response, viral proteins, especially non-structural proteins (including these of ubiquitous respiratory viruses: RSV, IAV, and SARS-CoV- 2), compromise this response, promoting viral spread (*49*). It is thus intuitive to expect that virus–host cell interactions are more complex than in the case of poly(I:C).

Previously, we showed that IFNβ-priming enhances cellular responses to poly(I:C), resulting in faster activation of IRF3 and NF-κB, which is also more uniform across the cell population (*17*). Here, we found that in IFNβ-prestimulated cells, STAT signaling is terminated within 4 h of poly(I:C) stimulation, which reprograms cells from IFNβ-responders to IFNβ-producers. Reprogramming of IFNβ-primed cells results from depletion of IFNAR1 and is associated with activation of the PKR/EIF2α and the OAS/RNase L pathways. In RNase L KO cells, STAT1/2 activity lasts longer than in WT cells, whereas in PKR & RNase L dKO cells, STAT1/2 remain active 10 h after poly(I:C) stimulation. Because of the accumulation of PKR and OAS1/2/3 in the course of IFNβ stimulation, the process of switching off STATs is more rapid when it follows prestimulation with IFNβ. RNase L degrades transcripts of STAT1/2-induced genes, including those coding for RIG-I, PKR, and OAS1/2/3. IFNGR1 transcript is also degraded, rendering poly(I:C)-stimulated or infected cells insensitive to stimulation with either IFNβ or IFNγ. Translational and transcriptional inhibition of IκBα and A20 enhances activation of NF-κB and IRF3, inducing expression of genes coding for cytokines IFNβ, IL6, and IL8. The transcript for IFNβ is resistant to RNase L (*50, 51*) and transcripts for IL6 and IL8 are only weakly degraded.

Under physiological conditions, conversion of the IFNβ-responding to the IFNβ-producing cells, that follows the progression of infection, may be crucial in sustaining and relaying the antiviral alert ahead of propagating virus. This allows the not-yet-infected cells to enter the antiviral state. In turn, rapid RNase L activation and termination of STAT signaling in subsequently infected cells may help to override the virus program (*52*), attenuate cell metabolism, and slow down or inhibit virus replication. It was demonstrated recently that RNase L controls protein synthesis not only by degrading ribosomal RNAs (*41*), but even more rapidly by degrading tRNA and Y-RNA (*53*).

Our findings led us to a comprehensive computational model allowing us to analyze responses to combined IFNβ and poly(I:C) stimulations. The model couples five regulatory modules: poly(I:C), NF-κB, IRF3, IFNβ and STAT, which are intertwined by positive and negative feedback loops. The main positive feedback loop augments paracrine IFNβ signaling such that IFNβ-primed cells upon stimulation with poly(I:C) become IFNβ producers. The main negative feedback causes that the STAT program is triggered only transiently in cells challenged with poly(I:C), and is terminated in IFNβ-primed cells upon poly(I:C) stimulation. The model parameters were constrained based on an ample set of experiments on WT as well as KO cells. By systematic simplification of the model we were able to reduce the number of independent parameters and reach parameter identifiability.

In summary, we found that poly(I:C) turns IFNβ-responding cells into IFNβ-secreting by terminating the STAT transcriptional program and triggering the IRF3/NF-κB transcriptional program. We elucidated the molecular mechanism underlying the switch and showed that the PKR and RNase L pathways activated by poly(I:C) play a decisive role. Based on our experimental findings, we developed and constrained a comprehensive computational model of the innate immune responses to IFNβ and poly(I:C).

## MATERIALS AND METHODS

### Cell lines

A549 cells (derived from a 58-year-old Caucasian male) and BEAS-2B (derived from normal bronchial epithelium obtained from autopsy of non-cancerous individuals) were purchased from ATCC and cultured in Ham’s F-12K (Kaighn’s) Medium (Gibco), supplemented with 10% FBS (Gibco) and 100 mg/ml penicillin/streptomycin mix (Sigma-Aldrich). HeLa cells were grown on Dulbecco’s modified Eagle’s medium with 4.5 g/l of D-glucose and 0.1 mM L-glutamine (Gibco) supplemented with 10% FBS and 100 mg/ml penicillin/streptomycin mix. The *EIF2AK2* (PKR) KO, *MAVS* KO, *IRF3* KO, and *TNFAIP3* (A20) KO A549 cell lines were generated using the CRISPR/Cas9 method as described further. *RNASEL* (RNase L) KO, another *EIF2AK2* (PKR) KO (used for preliminary studies), and *EIF2AK2* (PKR) & *RNASEL* (RNase L) double KO (dKO) cell lines were gifts from Dr. Bernard Moss (*54*). All cell lines were cultured under standard conditions (37 ℃, 5% CO_2_) and were passaged upon reaching 90% confluency (every 2–3 days). All cell lines were routinely tested for mycoplasma contamination using LookOut Mycoplasma qPCR Detection Kit (Merck 200-664-3). None of the cell lines were authenticated but low passage numbers were utilized.

### Cell line generation

For genome editing, wild-type A549 cells were transfected with sgRNA and Cas9-expressing plasmids. For MAVS gene knockout, an sgRNA targeting sequence GTAGATACAACTGACCCTGT in exon 6 of MAVS gene was designed using an online tool: http://www.benchling.com. The pair of sgRNA oligonucleotides (delivered by Oligo, IBB PAS, Warsaw) were inserted into the BbsI site of the vector backbone expressing Cas9 and GFP: pSpCas9(BB)- 2A-GFP (PX458) (Addgene; plasmid number 48138). For knockouts of *EIF2AK2* (PKR)*, IRF3* and *TNFAIP3* (A20) genes, commercially available plasmids (pCMV-Cas9-RFP and pCMV-Cas9-GFP) from Sigma-Aldrich was used targeting sequence AATACATACCGTCAGAAGCAGG in exon 3 of *EIF2AK2* (PKR) gene, ACACCTCTCCGGACACCAATGG in exon 4 of *IRF3* gene, or GTCATAGCCGAGAACAATGGGG in exon 5 of *TNFAIP3* (A20) gene. After reaching 80% confluency, cells were harvested, counted, suspended in a concentration of 1.5×10^6^ in 100 µl cell line nucleofector solution T (Lonza) and nucleofected with 5 μg of a plasmid using program X-001 in the Nucleofector IIb device (Lonza) according to the manufacturer’s procedure. Transfected cells were seeded into ϕ = 60 mm dishes in full medium. After 2 days, cells were washed with PBS, trypsinized, collected in full medium, and analyzed using the FACSaria sorter (BD Biosciences). Single fluorescent cells were sorted into 96-well plates in full medium. Individual clones were expanded and the level of PKR, MAVS, IRF3, or A20 protein was analyzed by Western blotting. Selected clones devoid of PKR, MAVS, IRF3, or A20 were further validated by sequencing by an external company (Oligo, IBB PAS, Warsaw).

### Virus amplification and isolation

Respiratory Syncytial Virus A2 strain and Influenza A virus H1N1, strain A/PR/8/34, were purchased from ATCC and amplified in HeLa or MDCK cells, respectively. Cells were seeded on 225 cm^2^ Tissue Culture Flasks (Falcon) and cultured as described above for 2–3 days until reaching 90% confluency. On the day of infection, virus growth medium was prepared: DMEM + 2% FBS for RSV or MEM basal medium + 0.3% BSA and 1 μg/ml of TPCK-Trypsin for IAV. The dilutions of virus were prepared in appropriate media, with target MOI around 0.05. Culture media were removed, cells were washed once with PBS, and overlaid with 10 ml of inoculum. Virus was allowed to adsorb to cells for 2 h at 37 ℃ with occasional stirring. Then, additional virus growth medium was added to a total volume of 40 ml per flask. Infected cells were cultured at 37 ℃ until the development of cytopathic effects could be observed in at least 80% of cells (typically around 3 days for IAV and 5 days for RSV). Virus-containing culture fluid was then collected and clarified by centrifugation at 3000×*g*, 4 ℃, for 20 min. Then, virus particles were precipitated by adding 50% (w/v) PEG 6000 (Sigma-Aldrich) in NT buffer (150 mM NaCl, 50 mM Tris-HCl, pH 7.5) to a final concentration of 10% and stirred gently at 4 ℃ for 90 min. Virus was centrifuged at 3250×*g*, 4 ℃, for 20 min and re-centrifuged after removing supernatant to remove the remaining fluid. Pellet was suspended in 1 ml of NT buffer, supplemented with 20% sucrose in case of RSV, aliquoted, and stored at −80 ℃.

### Virus quantification

Virus concentration in collected samples was quantified using immunofluorescence staining of cells expressing viral proteins. HeLa or MDCK cells were seeded on microscopic cover slips and cultured upon reaching 90–100% confluency. Serial dilutions of virus samples were obtained in the virus growth medium in the 10^−3^ to 10^−6^ range. After washing with PBS, cells were overlaid in duplicates with diluted virus, which was allowed to adhere for 2 h with occasional stirring. Afterwards, the virus-containing medium was removed, cells were overlaid with fresh virus growth medium and cultured for 16 h (for IAV) or 24 h (for RSV). Then, cells were washed with PBS and fixed with 4% formaldehyde for 20 min, at room temperature. Cells containing stained viral proteins were counted using a Leica TCS SP5 confocal microscope. Virus concentration was calculated using the following formula: (avg. number of infected cells)/(dilution factor × volume containing virus added) = infectious particles/ml.

### Stimulation protocols

#### Cell treatment, stimulation with poly(I:C) and IFNs

Cells were seeded in a medium containing 10% FBS on dishes, multi-well plates or coverslips, depending on the type of experiment, and typically allowed to adhere overnight at 37 ℃. Before poly(I:C) (Sigma-Aldrich) stimulation, the medium was changed to antibiotic-free F12K with 2% FBS at least 4 h after seeding, before overnight incubation. Poly(I:C) was delivered to cells by means of lipid-based transfection, using Lipofectamine LTX with Plus reagent (Thermo Fisher Scientific) as a reagent of choice. Modified manufacturer’s protocol optimized for transfection was used; poly(I:C) was mixed with Plus reagent diluted in serum-free F12K and then mixed with F12K-diluted lipofectamine. Liposome-poly(I:C) complexes were allowed to form for 20 minutes before adding them to cells at room temperature. Per 3×10^5^ cells in 1 ml of medium on a ϕ = 30 mm dish, 1 µl or 10 µl poly(I:C) stock solution (100 µg/ml) was added together with 6 µl of lipofectamine and 6 µl of Plus reagent, diluted in 100 µl of serum-free F12K. These amounts were appropriately scaled up or down for use with different numbers of cells and medium volumes in distinct experiments. Human interferon β (PBL Assay Science) was used at a typical concentration of 1000 U/ml and, while used in conjunction with poly(I:C), added to cells up to 24 h before poly(I:C), depending on experimental requirements. To study STAT pathway activation in cells exposed to poly(I:C) for 4, 6 or 10 h (with or without IFNβ pre-stimulation), IFNα 200 U/ml (Peprotech 300-02AA), IFNβ 1000 U/ml, or IFNγ 20 ng/ml (Gibco PHC4031) were added for last 30 min before protein extraction. For IFNβ visualization, a brefeldin A solution (BD Biosciences) was added to cells for 2 h prior to fixation.

#### RSV and IAV infection

Cells were cultured as described above until reaching 90% confluency. Medium was then removed, cells were washed with PBS, and either RSV or IAV was added in F12K medium with 2% FBS. Virus was allowed to adhere for 2 h with occasional rocking to support even distribution. Cells were fixed or collected for analysis in time points as described in the Results section.

### RT-qPCR

*RNA isolation*. For gene expression analysis experiments, cells were seeded on 12- or 24-well plates (Falcon) at a density of, respectively, 1.0× or 1.5×10^5^ cells/well. Upon completed stimulation according to chosen protocol, cells were washed once with PBS and submitted to isolation of total RNA using PureLink RNA Mini Kit (Thermo Fisher Scientific), following manufacturer’s instructions: cells were harvested and vortexed in Lysis Buffer with 2-mercaptoethanol and then vortexed again with one volume of 70% ethanol. Upon transferring to the spin cartridge, cellular RNA content was bound to the column, washed with appropriate buffers and eluted, all by centrifugation at 12,000×*g*. Eluted RNA in RNase-free water was used immediately for reverse transcription or stored for later use at −80 ℃.

#### Reverse transcription

RNA concentration and quality was determined by measuring UV absorbance of samples diluted 1:100 in 10 mM Tris-HCl (pH 7.5) at 260 and 280 nm, using the Multiskan GO spectrophotometer (Thermo Fisher Scientific). Independently, RIN index for isolated RNA was checked using Agilent 2100 Bioanalyzer (see below). Around 1 µg of RNA was used as a template for reverse transcription, performed using High-Capacity cDNA Reverse Transcription Kit (Thermo Fisher Scientific). Following manufacturer’s protocol, diluted RNA samples were mixed 1:1 with freshly prepared Master Mix containing RT Buffer, RT Random Primers, dNTP Mix, MultiScribe Reverse Transcriptase and RNase Inhibitor. Reaction was performed in MasterCycler Gradient thermal cycler (Eppendorf) under following conditions: 10 min at 25 ℃, 120 min at 37 ℃, and 5 min at 85 ℃.

#### Real-Time Quantitative Polymerase Chain Reaction

Reaction was performed in a QuantStudio 12K Flex Real-Time PCR system with Array Card block (Life Technologies). In most cases, 750 ng of cDNA from reverse transcription was mixed with reaction Master Mix and loaded onto TaqMan Array Card containing probes and primers for 24 genes (in 2 replicates), including endogenous reference controls (see Supplementary Table S2). Reaction was conducted using QuantStudio “Standard” protocol, with FAM/ROX chemistry. Upon completion, expression of target genes was analyzed based on absolute Ct quantification or classical ΔΔCt method with QuantStudio 12K Flex software, normalized against 18S gene expression. We found however that absolute Ct quantification of three control genes: HPRT1, GAPDH and 18S increase substantially in all experiments after poly(I:C) stimulation in A549 WT (but not in RNase L KO) cells. This is in agreement with the observation that ribosomal RNAs are degraded by RNase L (fig. S14 and Ref. *41*). Correspondingly, the absolute Ct quantification method was used in Fig. 5.

#### Digital PCR

Digital PCR (dPCR) measurements for IFNAR1 and IFNGR1 mRNA level were performed using QuantStudio 3D system (Thermo Fisher Scientific) (Fig. 3, D and E) and TaqMan Gene Expression Assays Hs1066118_m1 and Hs00988304_m1, respectively, according to manufacturer’s protocol. cDNA samples were prepared in an identical manner as in the case of RT-PCR experiments (see above).

### RNA analysis

rRNA integrity analysis was performed using Agilent 2100 Bioanalyzer System for dual purposes of verifying quality of isolated RNA for gene expression experiments and for indirect measurement of RNase L activity within cells. To this end, RNA samples and RNA ladder were prepared according to manufacturer’s protocol, by heat denaturation at 70 ℃ for 2 min and subsequent cooling on ice. Agilent RNA 6000 Nano chips were filled with a Gel-Dye Mix using a dedicated priming station. Five μl of RNA marker and 1 μl of RNA sample or ladder were then loaded onto the chip. The chip was vortexed for 1 min and immediately analyzed using the Bioanalyzer. Results, in the form of RIN quantifications and electropherograms, were collected using the Agilent 2100 Expert software.

### ELISA

For measuring IFNβ production, cells were seeded on 96-well plates (Falcon) at the density of 20,000 cells/well. Upon requested stimulation time according to a chosen protocol, culture medium from cells was collected and stored at −20 ℃ until further analysis.

IFNβ levels were estimated using the VeriKine Human IFNβ ELISA Kit (PBL Assay Science). Standards and diluted samples in duplicates or triplicates were added to a precoated plate included in the kit and incubated for 1 h. After subsequent washing, antibody solution was prepared and added to wells for another 1 h, followed by another washing and 1 h incubation with HRP solution. Finally, TMB Substrate Solution was added to wells and developing color reaction was stopped after 15 min with the addition of Stop Solution. Optical densities of samples after resulting color development were determined using Multiskan GO plate reader (Thermo Fisher Scientific) set to 450 nm, with wavelength correction at 570 nm. IFNβ concentrations were obtained according to a standard curve obtained by fitting a 4-parameter function (containing a sigmoid and a linear term) to measurements of 8 standards.

### Immunostaining

Confocal microscopy imaging was used in order to visualize subcellular localization of studied proteins in stimulated cells. Cells were seeded on ϕ = 12 mm round glass coverslips, which were previously washed in 60% ethanol/40% HCl, thoroughly rinsed with water and sterilized. Seeding density was 10^5^ cells/coverslip 24 h prior to treatment or 7×10^4^ cells/coverslip 48 h prior to treatment (experiments with IFNβ prestimulation). After stimulation (typically in duplicates) at desired time points, cells on coverslips were washed with PBS and immediately fixed with 4% formaldehyde (20 min, room temperature). Cells were then washed three times with PBS and incubated for 20 min at −20 ℃ with 100% cold methanol. After washing with PBS, coverslips with cells were blocked for 1.5 h with 5% BSA (Sigma-Aldrich) with 0.3% Triton X-100 (Sigma-Aldrich) in PBS at room temperature. After removing the blocking solution, coverslips with cells were incubated overnight at 4 ℃ with primary antibodies diluted in PBS containing 1% BSA and 0.3% Triton X-100. After washing cells five times with PBS, appropriate secondary antibodies conjugated with fluorescent dyes were added and incubated for 1 h at room temperature. Subsequently, cells were washed and their nuclei were stained for 10 min with 200 ng/ml DAPI (Sigma-Aldrich) to enable detection of nuclear outlines (Fig. 1, A and B; Fig. 2, C and D; Fig. 3C). After final washing in miliQ water, coverslips with stained cells were mounted on microscope slides with a drop of Mowiol (Sigma-Aldrich). Cellular sublocalization of stained proteins was observed using Leica SP5 confocal microscope and the Leica Application Suite AF software. At least three images from each experimental variant, each containing no less than 50 cells, were captured.

### Western blot

Cells were washed in PBS two times, lysed in the Laemmli sample buffer containing DTT and boiled at 95 ℃ for 10 min. Even amounts of each protein sample were separated on a 8–10% polyacrylamide gel using Mini-PROTEAN Tetra Cell electrophoresis system (Bio-Rad). Upon completion of electrophoresis, proteins were transferred to the nitrocellulose membrane using wet electrotransfer in the Mini-PROTEAN apparatus, according to the modified Towbin method (400 mA, 1 h). Membrane was rinsed with TBST (TBS buffer containing 0.1% Tween-20, Sigma-Aldrich) and blocked for 1 h with 5% BSA/TBS or 5% non-fat dry milk to prevent nonspecific antibody binding. Subsequently, membranes were incubated at 4 ℃ overnight with one of the primary antibodies diluted in the 5% BSA/TBS buffer. After thorough washing with TBST, membranes were incubated with secondary antibodies conjugated with specific fluorochrome or horseradish peroxidase diluted in 5% nonfat dry milk for 1 h, RT. Chemiluminescent reaction was developed with the Clarity Western ECL system (Bio-Rad). Specific proteins were detected in the dark room on the medical X-ray film blue (Agfa) with an Agfa brand developer and fixer or using the ChemiDoc MP Imaging System (Bio-Rad). List of antibodies and all other materials and reagents is provided in Supplementary Table S2 in Supplementary Materials.

### Computational methods

We developed the computational model within RuleBender (*55*), that uses BioNetGen (*43*) as its engine for both deriving a system of ODEs from chemical reactions specified in the BioNetGen language (BNGL) and performing deterministic simulations of the resultant system (using a sparse solver). In Supplementary Materials, in addition to the source BNGL file, we provide the model definition exported to an SBML file.

We fitted kinetic parameters of the model with PyBioNetFit (*56*). To be able to use quantifications of Western blots as reference data for fitting parameters of the nondimensionalized model, we normalized each blot to its maximum. To focus on the fold change of measured observables and at the same time ignore background noise or relative change of low-level signals, we transformed both the normalized experimental data and corresponding max-normalized values extracted in experimental time points from simulated model trajectories by adding 0.03 and then taking logarithm. In the final normalization step, we subtracted the mean value of each such max-normalized and log-transformed data series. Overall, this normalization provides the best alignment of data series originating from relative experimental measurements and from trajectories of corresponding variables of the nondimensionalized model. In fitting, we started from 300 random points in the parameter space and minimized the objective function, expressed as the sum of squares of differences of such log-transformed experimental and simulated data points, by means of stochastic optimization (scatter search) followed by refinement according to the Nelder–Mead (simplex) method. We additionally imposed that before stimulation the amount of unbound cytoplasmic IκBα is < 0.2 and that at 6 h after stimulation with poly(I:C), the total amount of IκBα is < 0.3. Fitted parameter values are given in Supplementary Table S1.

The sensitivity matrix *S* was calculated based on model trajectories for all experimental stimulation protocols with the corresponding series of measurement time points transformed and normalized in the same way as for fitting. The number of columns of *S* is equal to the number of model parameters (38 in the case of the final model) and the number of rows is equal to the number of independent experimental data points used for parameter fitting (2915). We approximated the measured variable derivatives with respect to the model parameters using finite central differences (we decreased and increased each parameter value by 1% of its nominal value).

Based on the sensitivity matrix *S* we calculated singular values and ratios of the parameter estimation error *ξ* _i_ (where 1 ≤ *i ≤ p* is an index of a parameter) to the experimental error in the final fitted model (*57*). For the latter we assumed that all experimental errors (in a log-scale) are independent and identically normally distributed. Lognormal distribution of Western blot data was shown by Kreutz *et al.* (*58*). Let *R_p_* denote the matrix composed of the first *p* rows of an upper triangular matrix *R* resulting from the QR decomposition of the sensitivity matrix *S*. Then, *ξ* _i_ is the ℓ^2^-norm of the *i*-th row vector of the inverse of *R_p_* (*57*).

## Supporting information

Software: Mathematical model of the innate immune response (zipped BNGL snd SBML files).

Source Data: Uncropped Western blot images and uncropped full-resolution immunostaining images.

## Supplementary materials

- **Supplementary Figures and Tables (PDF file).** Supplementary Figures S1–S19, Supplementary Table S1 (model documentation), Supplementary Table S2 (reagents, antibodies, and other resources).
- **Source Data (PDF file).** Uncropped Western blot and full-resolution immunofluorescence images.
- **Software (ZIP file).** Mathematical model of the innate immune response (BNGL, SBML).

## Acknowledgments

We thank Dr. Bernard Moss for sharing A549 cell lines with PKR KO, RNase L KO, and PKR and RNase L double KO. We thank Bartosz Tarkowski for generating a CRISPR plasmid construct targeting MAVS gene, Frederic Grabowski, Paweł Nałęcz-Jawecki, Paulina Koza, and Karolina Tudelska for discussion. This research was performed using CePT-financed equipment. Numerical simulations were performed using PLGrid Infrastructure.

## Author contributions

ZK, MC and WP planned and performed experiments; JJB built the computational model. JJB and IK performed identifiability analysis and fitting; MK provided software tools and expertise; TL conceived and supervised the project; all authors contributed to writing the manuscript.

## Data and materials availability

The cells lines developed in this study — *EIF2AK2* (PKR) KO, *MAVS* KO, *IRF3* KO, and *TNFAIP3* (A20) KO A549 — are available from the corresponding author without restriction. All data needed to evaluate the conclusions in the paper are present in the paper or the Supplementary Materials: uncropped Western blot images and full-resolution uncropped immunostaining images are provided in Source Data; computer code produced in this study is available in this paper’s Supplementary Materials (in BNGL format and SBML format).

