## Supplementary Figures S1-S19 and Tables S1-S2. for "Non-self RNA rewires IFNβ signaling: A mathematical model of the innate immune response"

featuring the article

#### The PDF file includes:

#### Other Supplementary Material for this manuscript includes the following:

Source Data (separate PDF file)
Software (separate ZIP file)

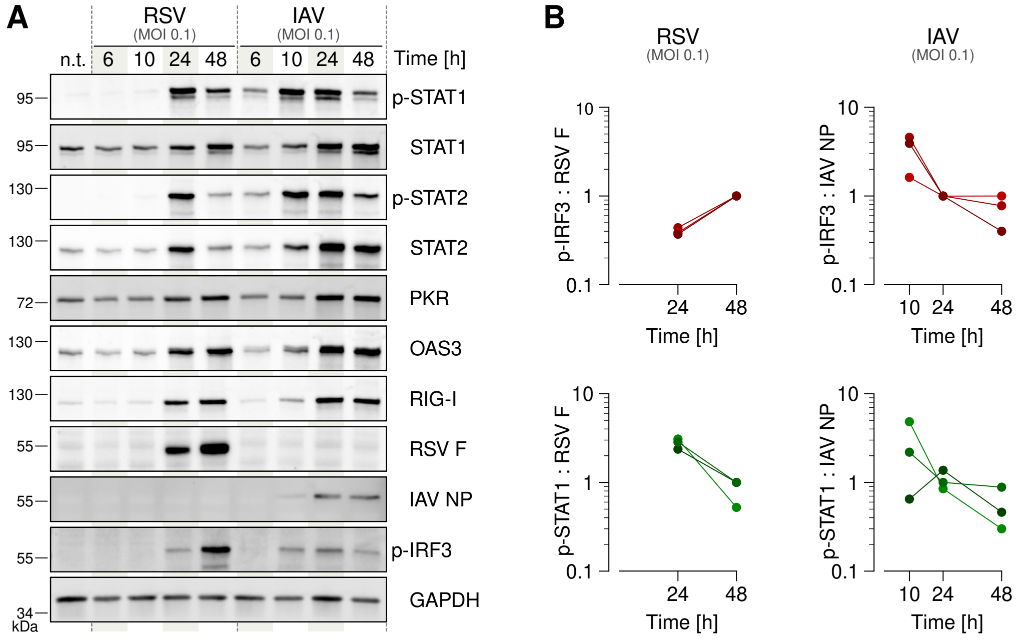

**Fig. S1**: **Viral infection with IAV and RSV triggers and terminates STAT1 and STAT2 activity.** (**A**) Western blots show p-STAT1/2, three considered ISGs (PKR, OAS3, RIG-I), p-IRF3, and viral proteins in A549 WT cells infected with RSV or IAV, each at an MOI of 0.1. Representative blots from 3 independent experiments are shown. See Source Data for corresponding uncropped blots. (**B**) Ratios of quantified and normalized levels of p-IRF3 -to- virus and p-STAT1 -to- virus.

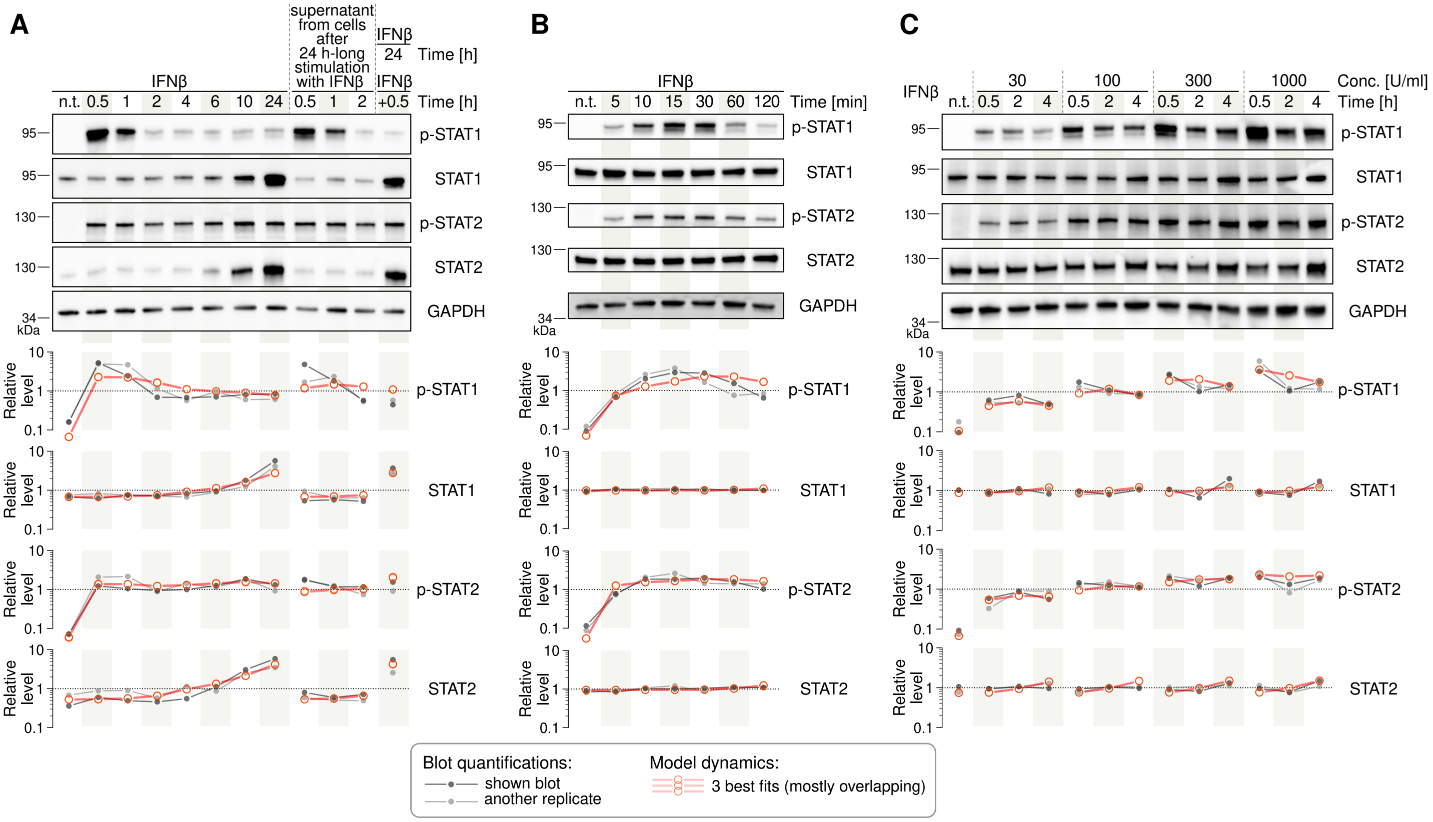

**Fig. S2**: **STAT1/2 signaling in response to stimulation with IFNβ.** (**A**) A549 WT cells stimulated with IFNβ (1000 U/ml) using the indicated protocol. Conditioned medium was collected from cell culture of cells stimulated with IFNβ for 24 h. (**B**) A549 WT cells stimulated with IFNβ (1000 U/ml) in indicated time points. (**C**) A549 WT cells stimulated with IFNβ at indicated concentrations for 0.5, 2, or 4 h.

Representative blots from 2 independent experiments are shown. Numerical solutions are continuous in time but, for the sake of comparison to experimental data, were extracted in experimental time points and connected with straight line segments to guide the eye. See Source Data for corresponding uncropped blots.

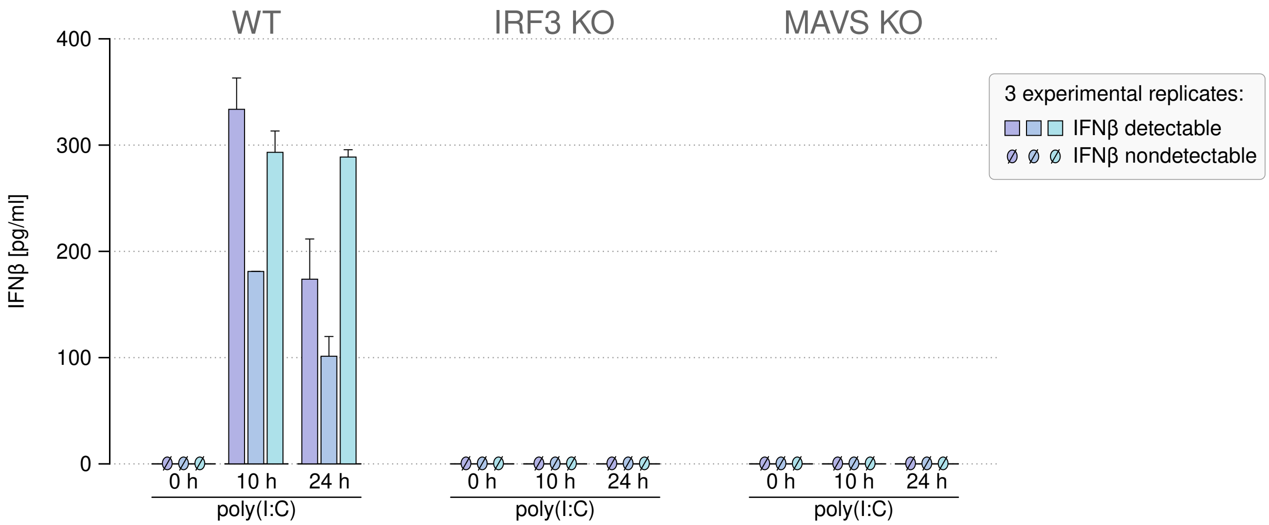

**Fig. S3**: **Secretion of IFNβ by A549 WT, IRF3 KO, and MAVS KO cells after poly(I:C).** ELISA measurements after stimulation with poly(I:C) of concentration 0.1 μg/ml in indicated time points. Error bars represent s.e.m. for two technical replicates in each experiment.

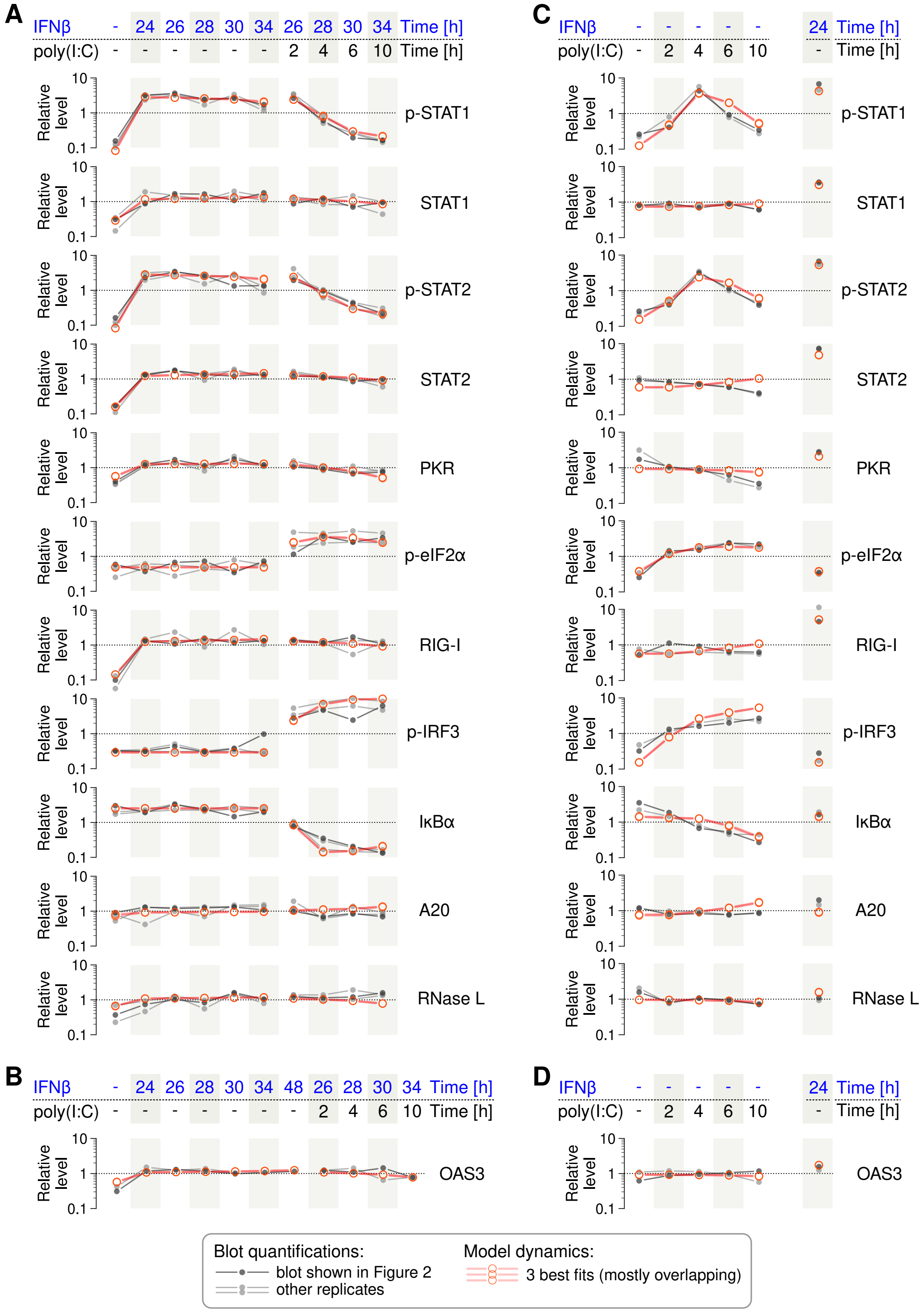

**Fig. S4**: **Comparison of experimental results from Fig. 2 to computational model predictions.** (**A**, **B**, **C**, **D**) Quantified Western blots from Fig. 2A, 2B, 2E, and 2F and their replicates juxtaposed with numerical solutions (continuous in time but, for the sake of comparison to experimental data, extracted in experimental time points and connected with straight line segments to guide the eye).

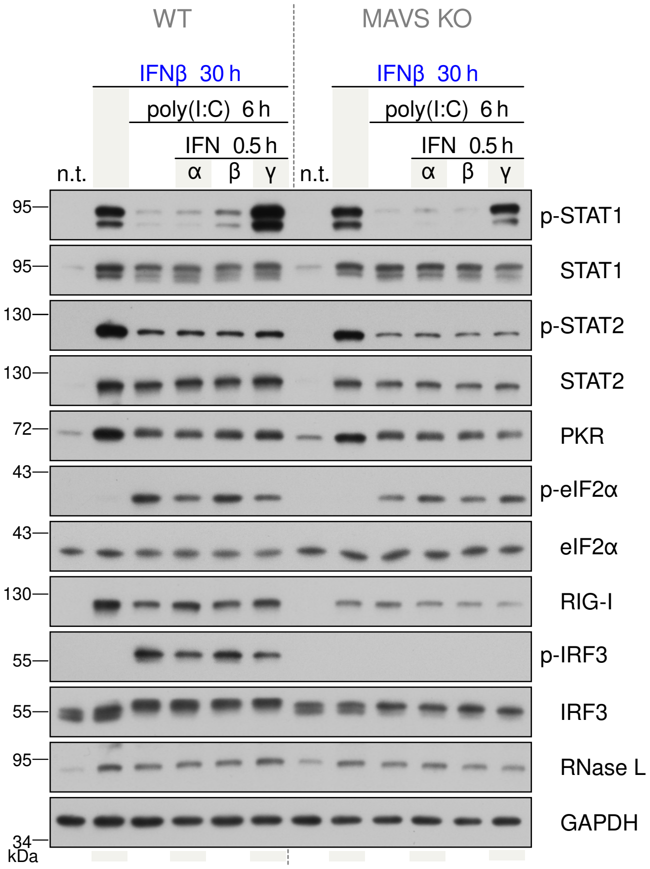

**Fig. S5**: **IFNβ-prestimulated A549 WT vs MAVS KO cells treated with poly(I:C) and restimulated with IFNα/β/γ.** Concentration of IFNβ: 1000 U/ml (prestimulation & restimulation), poly(I:C): 0.1 μg/ml, IFNα: 200 U/ml, IFNγ: 20 ng/ml. Graphical convention of stimulation protocols as in Fig. 3F in the main text. Representative blots from 2 independent experiments. Uncropped blots are shown in Source Data.

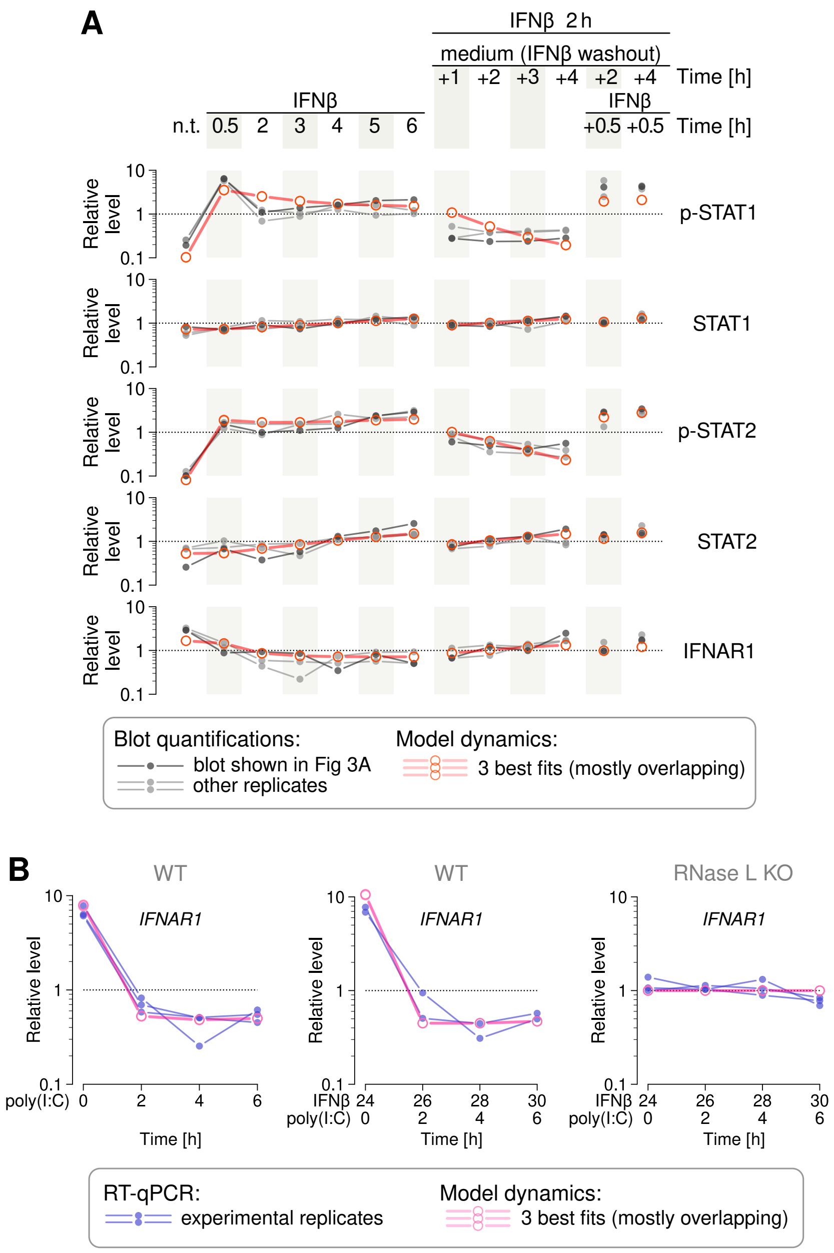

**Fig. S6**: **Comparison of experimental results from Fig. 3 to computational model predictions.** (**A**) Quantified Western blots from Fig. 3A juxtaposed with numerical solutions. (**B**) Digital PCR measurements from Fig. 3D juxtaposed with numerical solutions.

Numerical solutions are continuous in time but, for the sake of comparison to experimental data, were extracted in experimental time points and are connected with straight line segments to guide the eye.

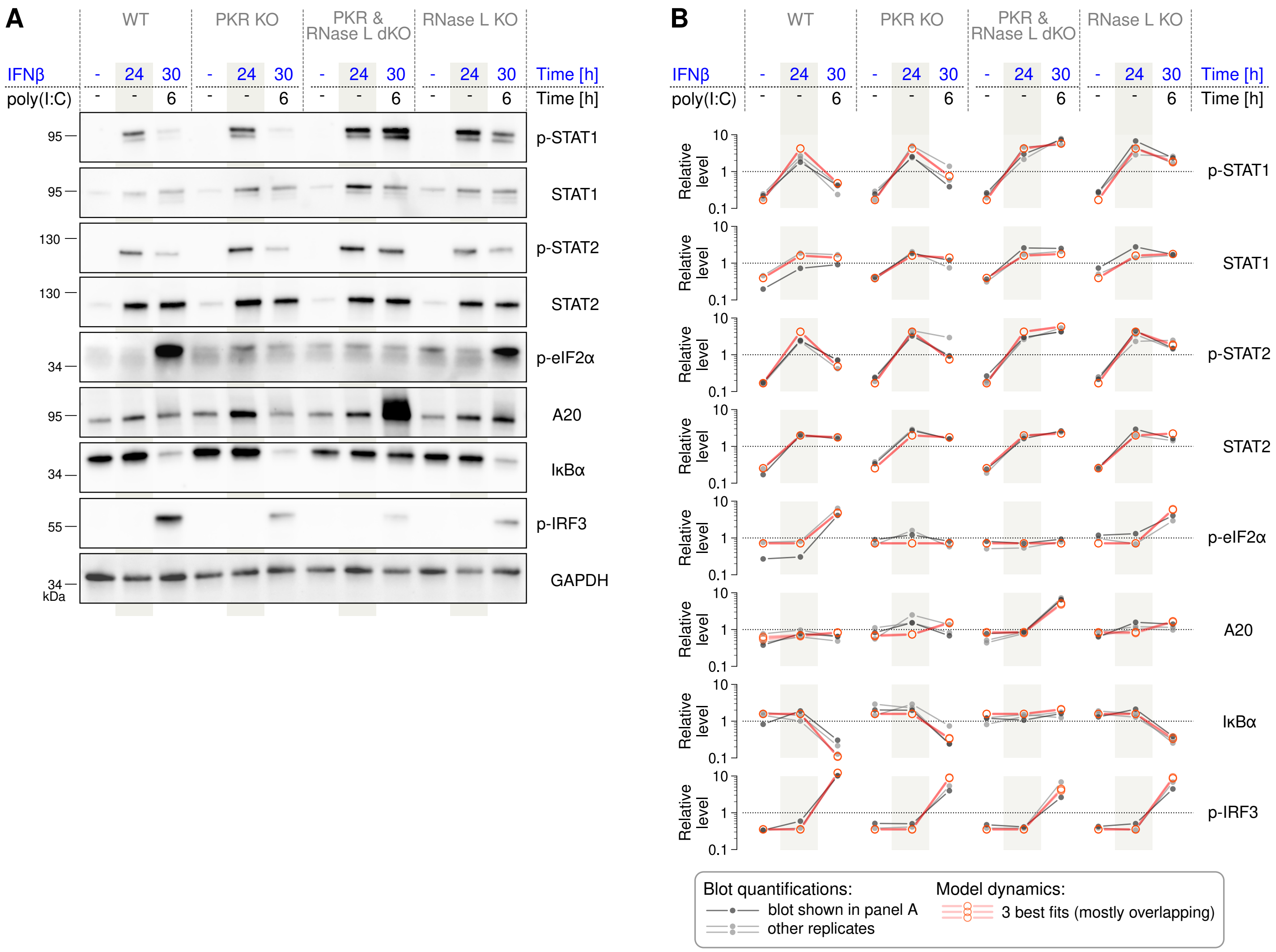

**Fig. S7**: **Response of A549 WT, PKR KO, PKR & RNase L double KO (dKO), and RNase L KO cells to IFNβ and poly(I:C).** (**A**) A549 WT, PKR KO, PKR & RNase L double KO, and RNase L KO cells were prestimulated with IFNβ (1000 U/ml) and then stimulated additionally with poly(I:C) (0.1 μg/ml). Representative blots from 3 independent experiments are shown. (**B**) Quantified Western blots juxtaposed with numerical solutions (continuous in time but, for the sake of comparison to experimental data, extracted in experimental time points and connected with straight line segments to guide the eye).

See Source Data for corresponding uncropped blots.

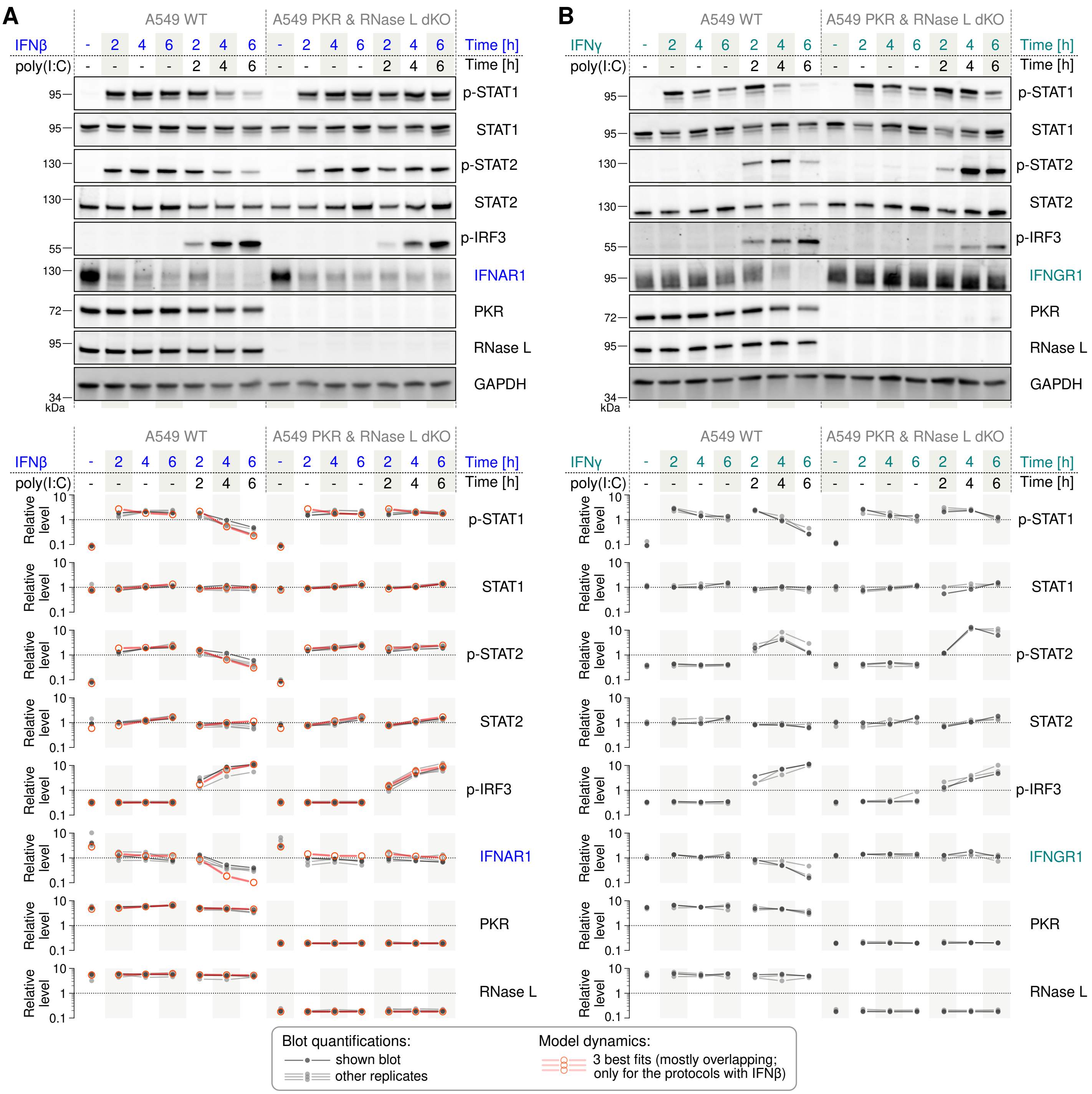

**Fig. S8**: **Response of A549 WT and PKR & RNase L double KO cells to simultaneous stimulation with IFNβ/γ and poly(I:C).** (**A**) A549 WT and PKR & RNase L double KO cells were stimulated with IFNβ (1000 U/ml) or with IFNβ (1000 U/ml) and simultaneously with poly(I:C) (0.1 µg/ml). Representative blots from 4 independent experiments are shown. Quantified Western blots juxtaposed with numerical solutions (continuous in time but, for the sake of comparison to experimental data, extracted in experimental time points and connected with straight line segments to guide the eye). (**B**) A549 WT and PKR & RNase L double KO cells were stimulated with IFNγ (20 ng/ml) or with IFNγ (20 ng/ml) and simultaneously with poly(I:C) (0.1 µg/ml). Representative blots from 3 independent experiments are shown. Blot quantifications are not juxtaposed with numerical solutions because the model does not include IFNGR1 (shown experiments were added in revision).

See Source Data for corresponding uncropped blots.

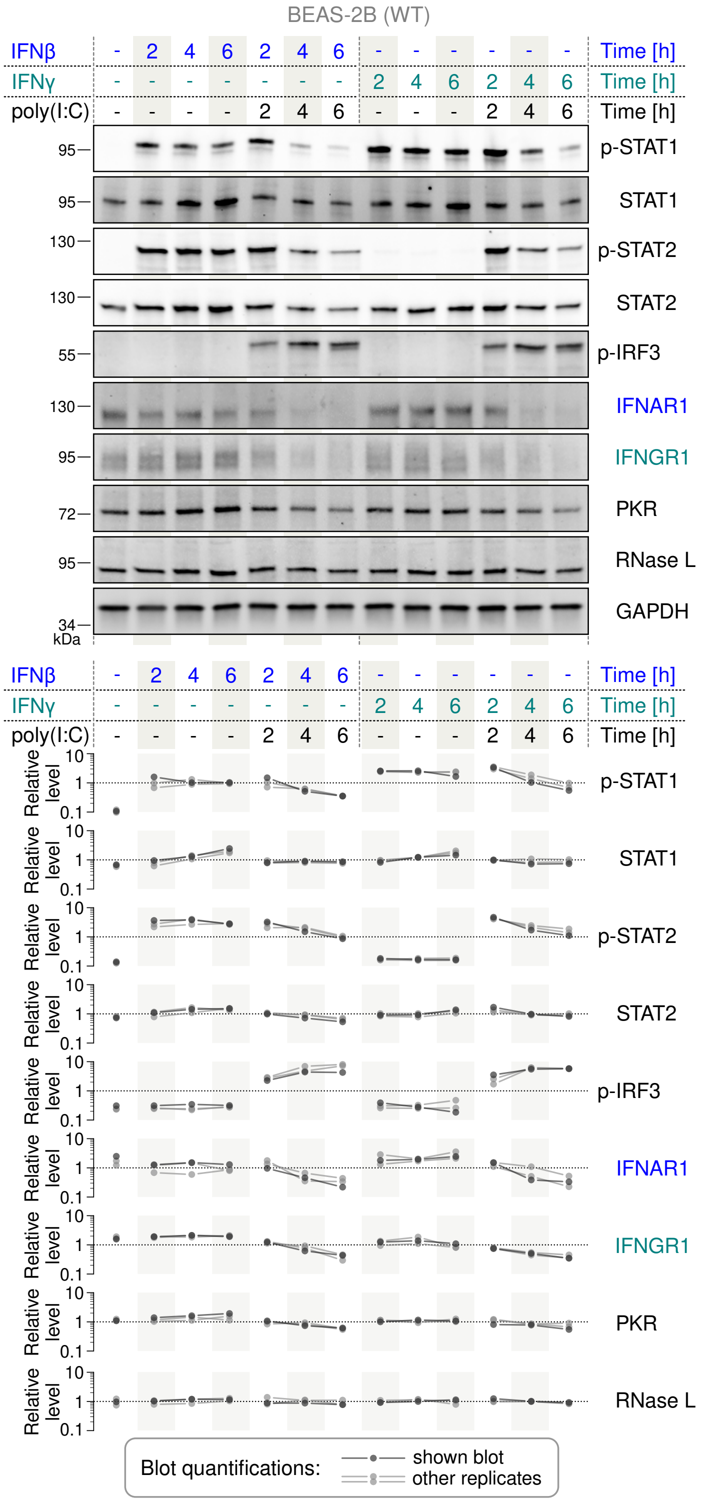

**Fig. S9**: **Response of BEAS-2B cells to simultaneous stimulation with IFNβ/γ and poly(I:C).** BEAS-2B WT cells were stimulated with IFNβ (1000 U/ml), with IFNβ (1000 U/ml) and simultaneously with poly(I:C) (0.1 µg/ml), with IFNγ (20 ng/ml), or with IFNγ (20 ng/ml) and simultaneously with poly(I:C) (0.1 µg/ml). Representative blots from 3 independent experiments are shown. Blot quantifications are not juxtaposed with numerical solutions because the model does not include IFNGR1 (shown experiments were added in revision). See Source Data for corresponding uncropped blots.

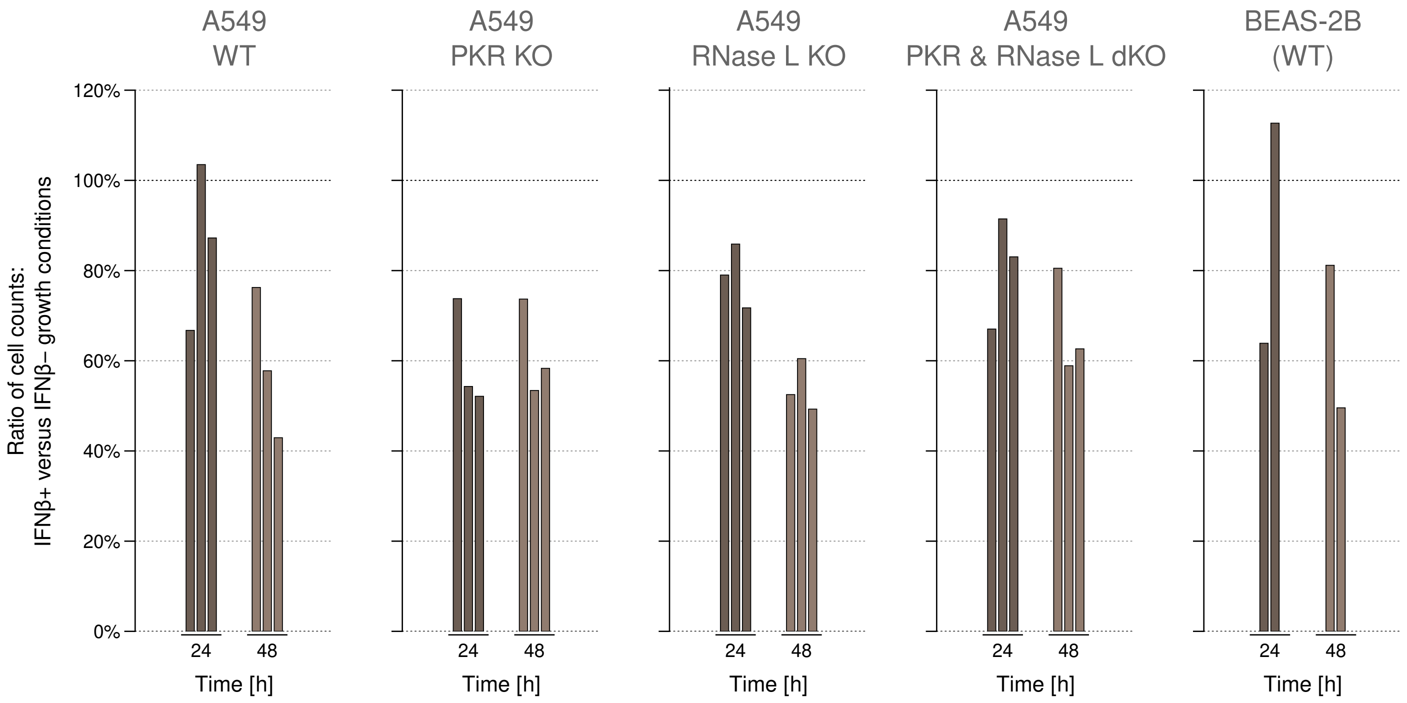

**Fig. S10**: **Antiproliferative effect of IFNβ stimulation on A549 WT, RNaseL KO, PKR KO and PKR & RNase L double KO cell lines as well as on BEAS-2B (WT) cells.** In IFNβ+ conditions, cells were stimulated with IFNβ at 1000 U/ml. Each bar represents an independent experiment.

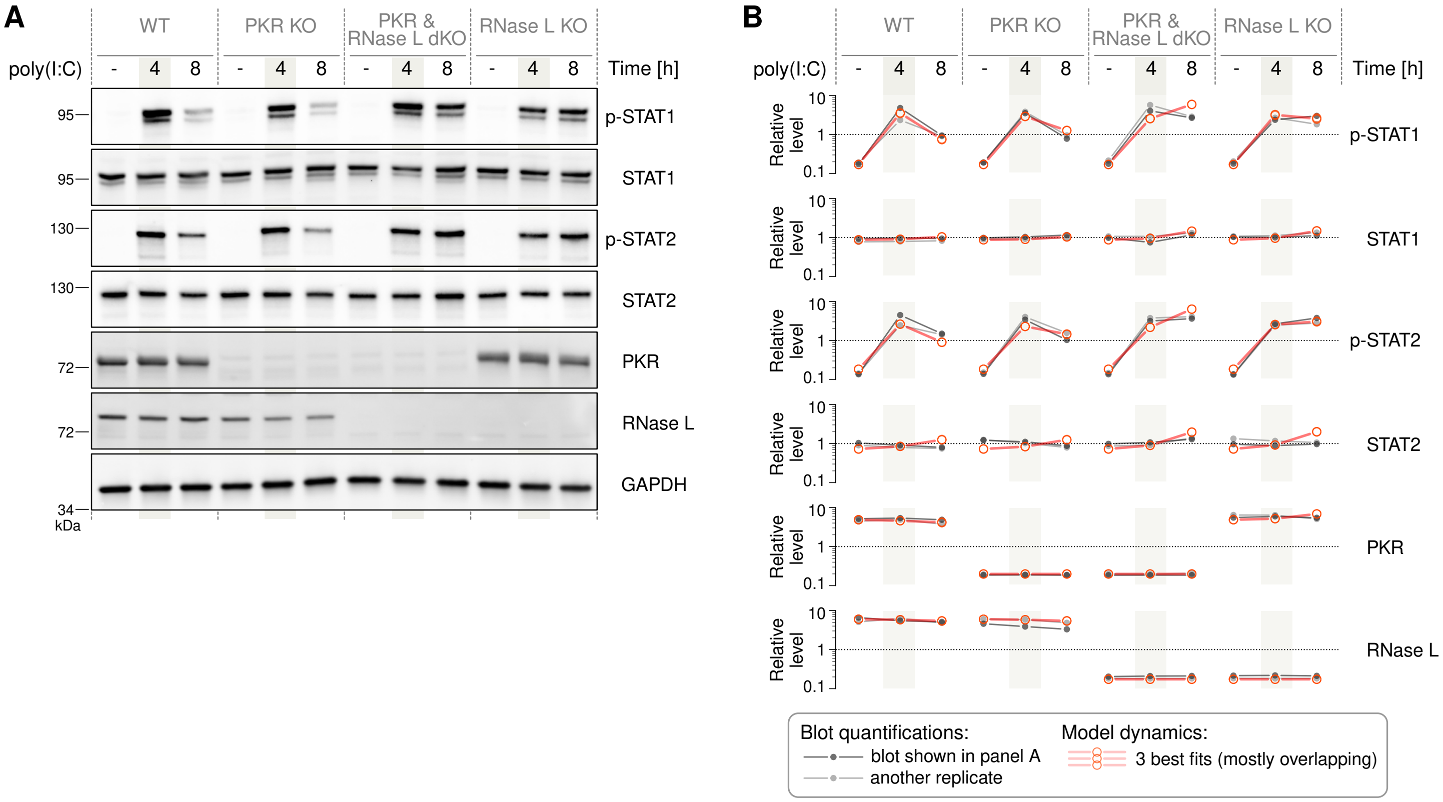

**Fig. S11**: **Response of A549 WT, PKR KO, PKR & RNase L double KO, and RNase L KO cells to poly(I:C). (A)** Concentration of poly(I:C) was 0.1 µg/ml. Representative blots from 2 independent experiments are shown. See Source Data for corresponding uncropped blots. **(B)** Quantified Western blots juxtaposed with numerical solutions (continuous in time but, for the sake of comparison to experimental data, extracted in experimental time points and connected with straight line segments to guide the eye).

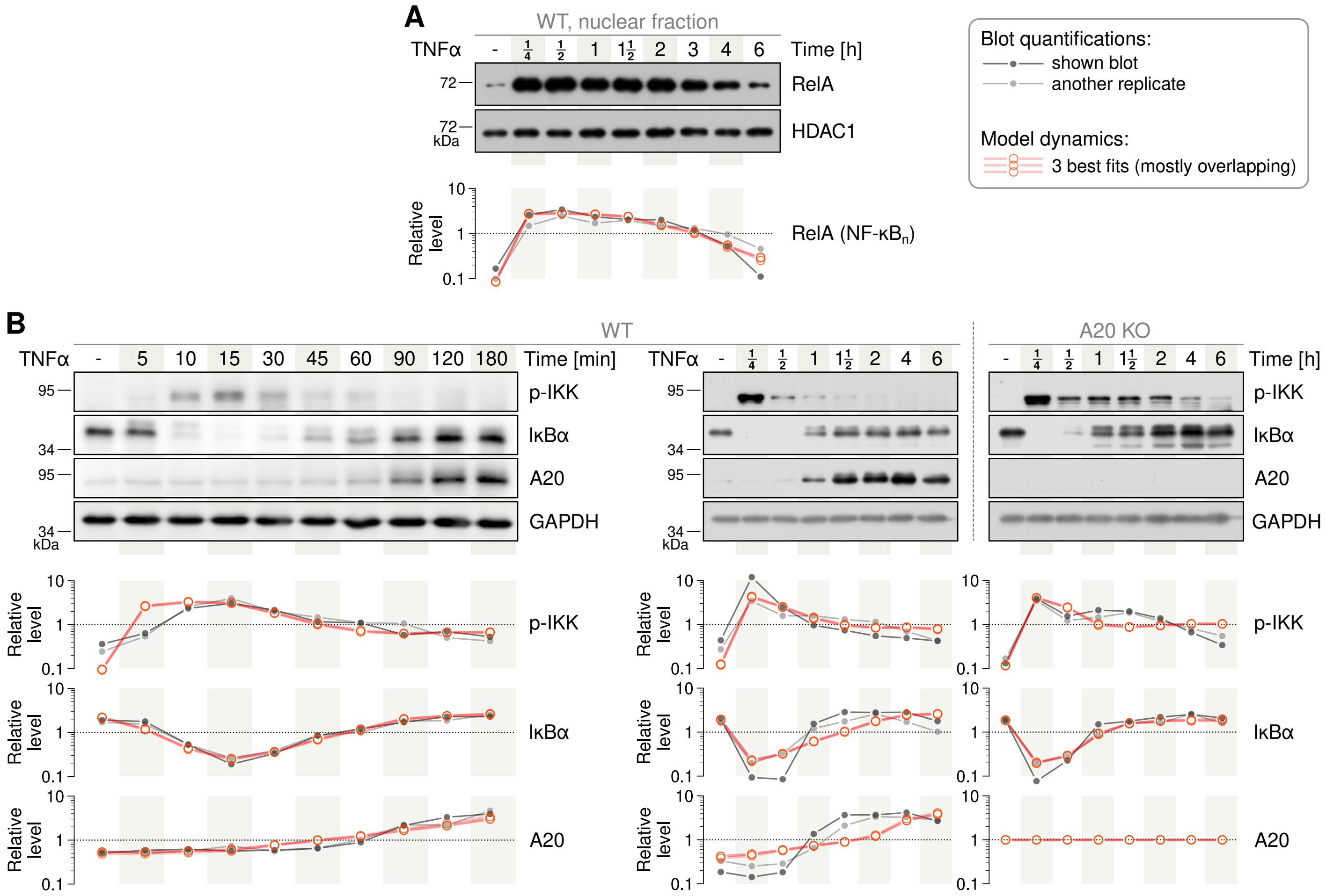

**Fig. S12**: **A549 WT and A20 KO cells stimulated with TNFα.** (**A**) Nuclear NF-κB in A549 WT cells stimulated with TNFα at 10 ng/ml. Representative blots from 2 independent experiments are juxtaposed with model trajectories. (**B**) A549 WT and *TNFAIP3* (A20) KO cells stimulated with TNFα at 10 ng/ml. Representative blots from 2 independent experiments are juxtaposed with model trajectories.

Uncropped blots are shown in Source Data.

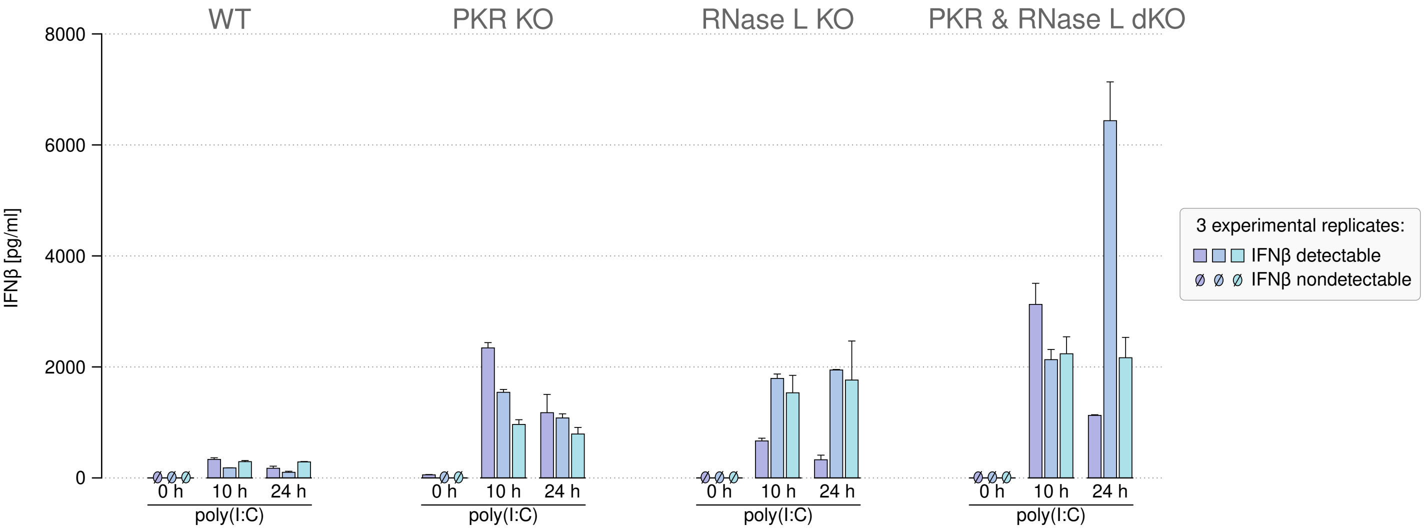

**Fig. S13**: **Secretion of IFNβ by A549 WT, PKR KO, RNase L KO, and PKR & RNase L double KO cells after stimulation with poly(I:C).** ELISA measurements of IFNβ secretion after stimulation with poly(I:C) (0.1 µg/ml) in indicated time points. Error bars represent s.e.m. for two technical replicates in each experiment.

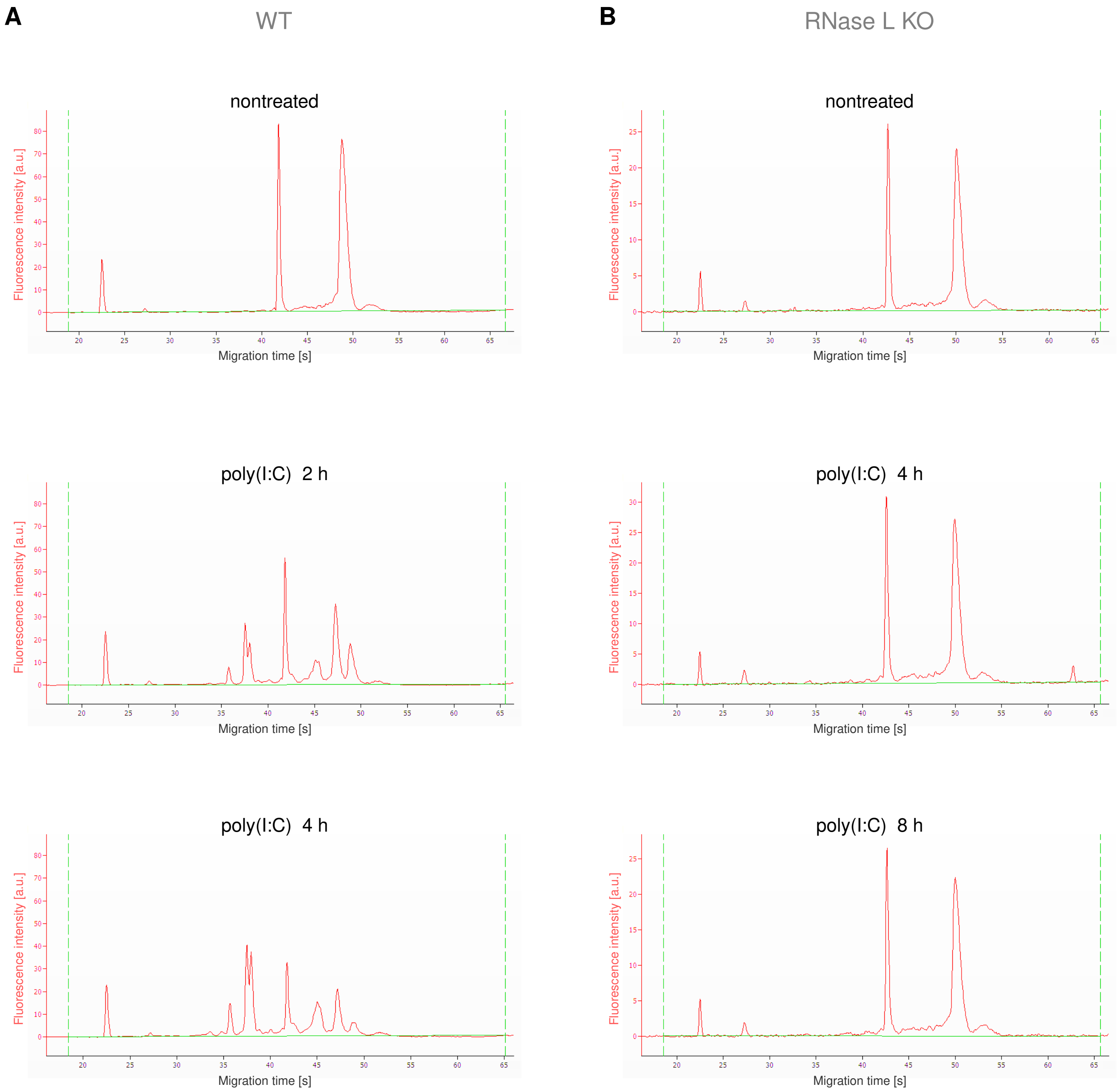

**Fig. S14**: **Degradation of rRNA in poly(I:C)-treated A549 (A) WT and (B) RNase L KO cells.** Concentration of poly(I:C) was 0.1 µg/ml. Representative results from 3 independent experiments.

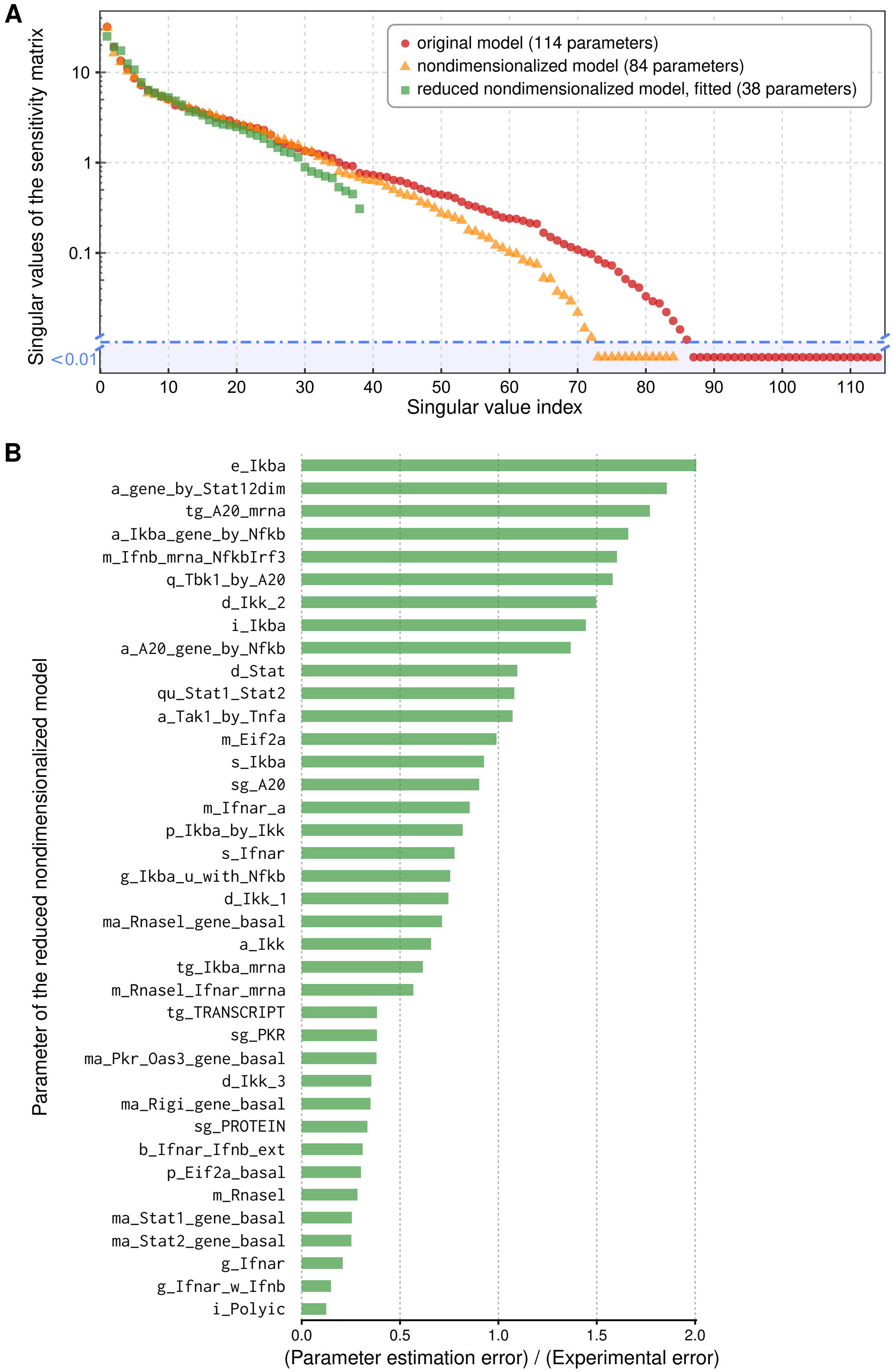

**Fig. S15**: **Singular values and parameter estimation errors.** (**A**) Singular values of the three considered models. (**B**) Ratio of the parameter estimation error to the experimental error in the final, fitted model.

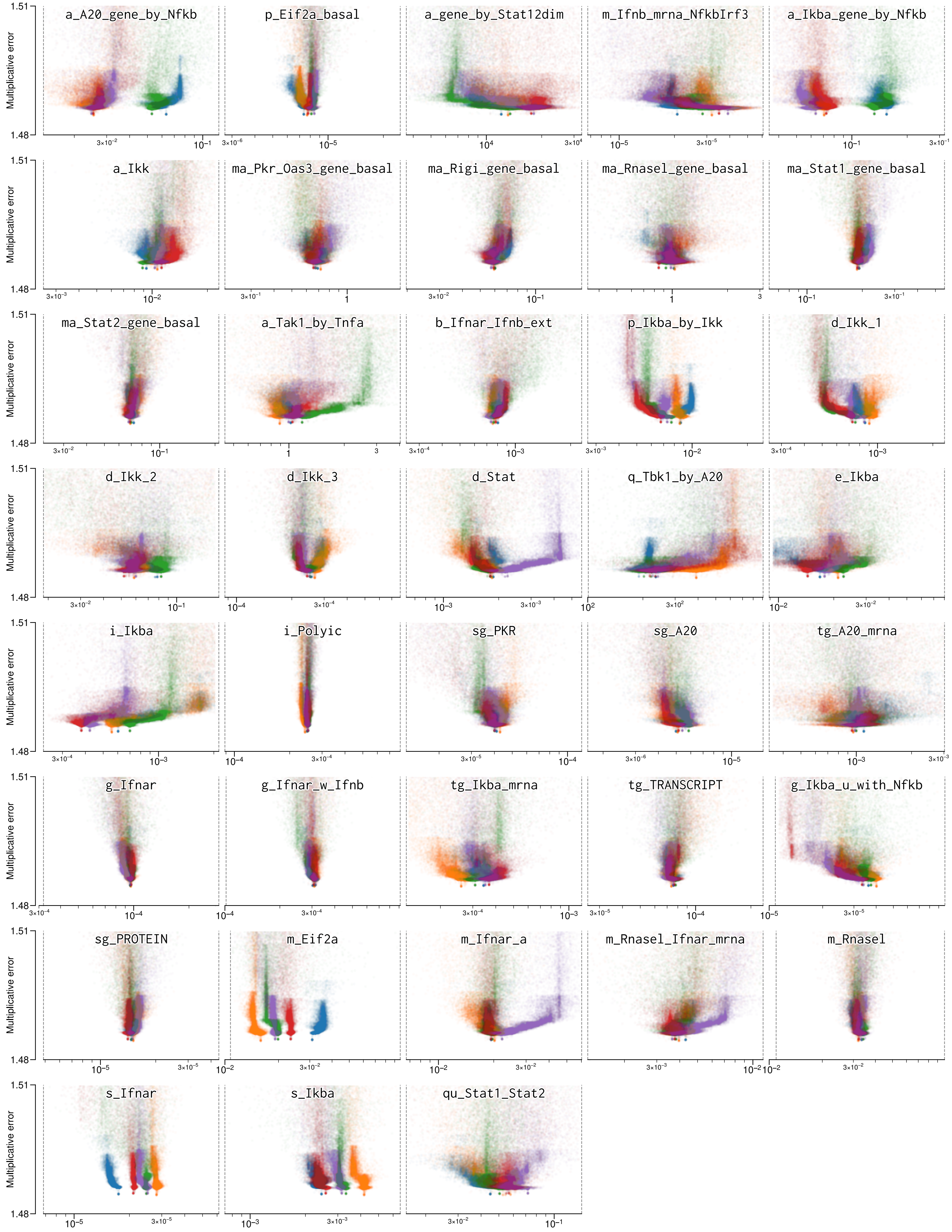

**Fig. S16**: **Convergence of parameter fitting.** Intermediate results from scatter search are refined with the Nelder–Mead (simplex) method. Parameter bounds are shown using vertical gray dashed lines.

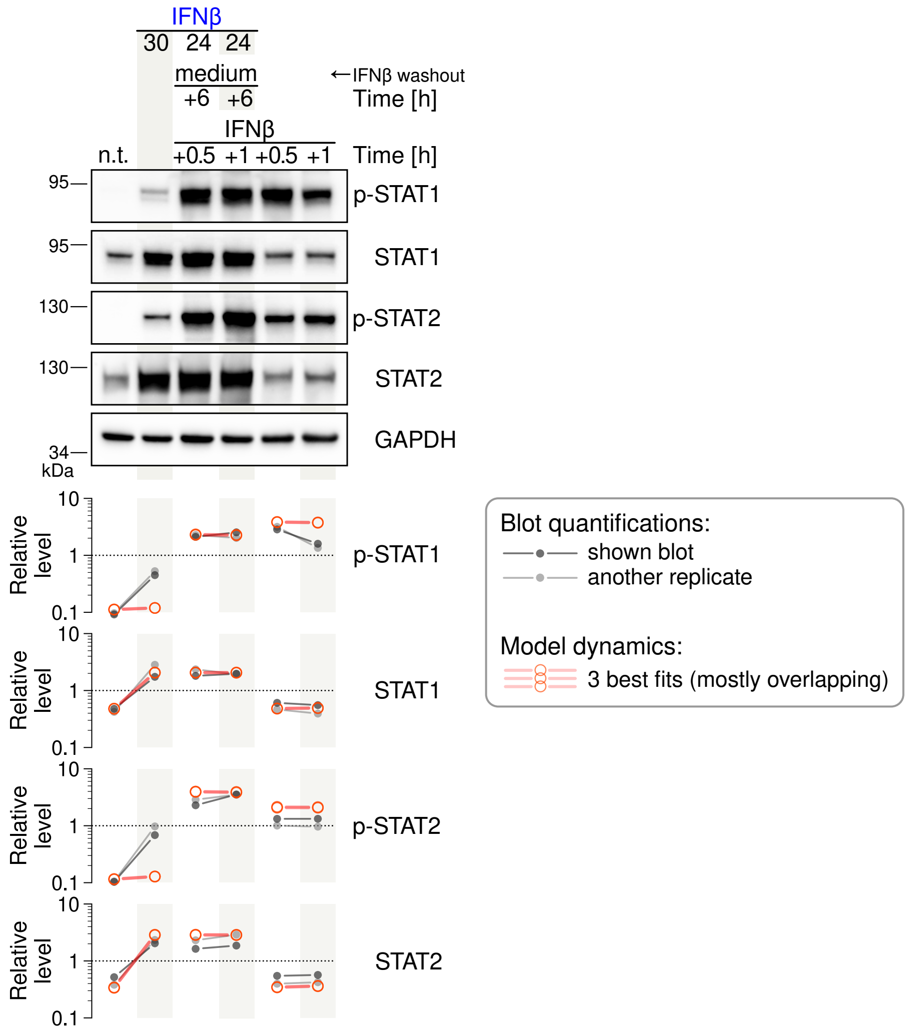

**Fig. S17**: **A549 WT cells stimulated and restimulated with IFNβ.** Concentration of IFNβ was 1000 U/ml. Graphical convention of stimulation protocols as in Fig. 3A in the main text. Representative blots from 2 independent experiments. Uncropped blots are shown in Source Data.

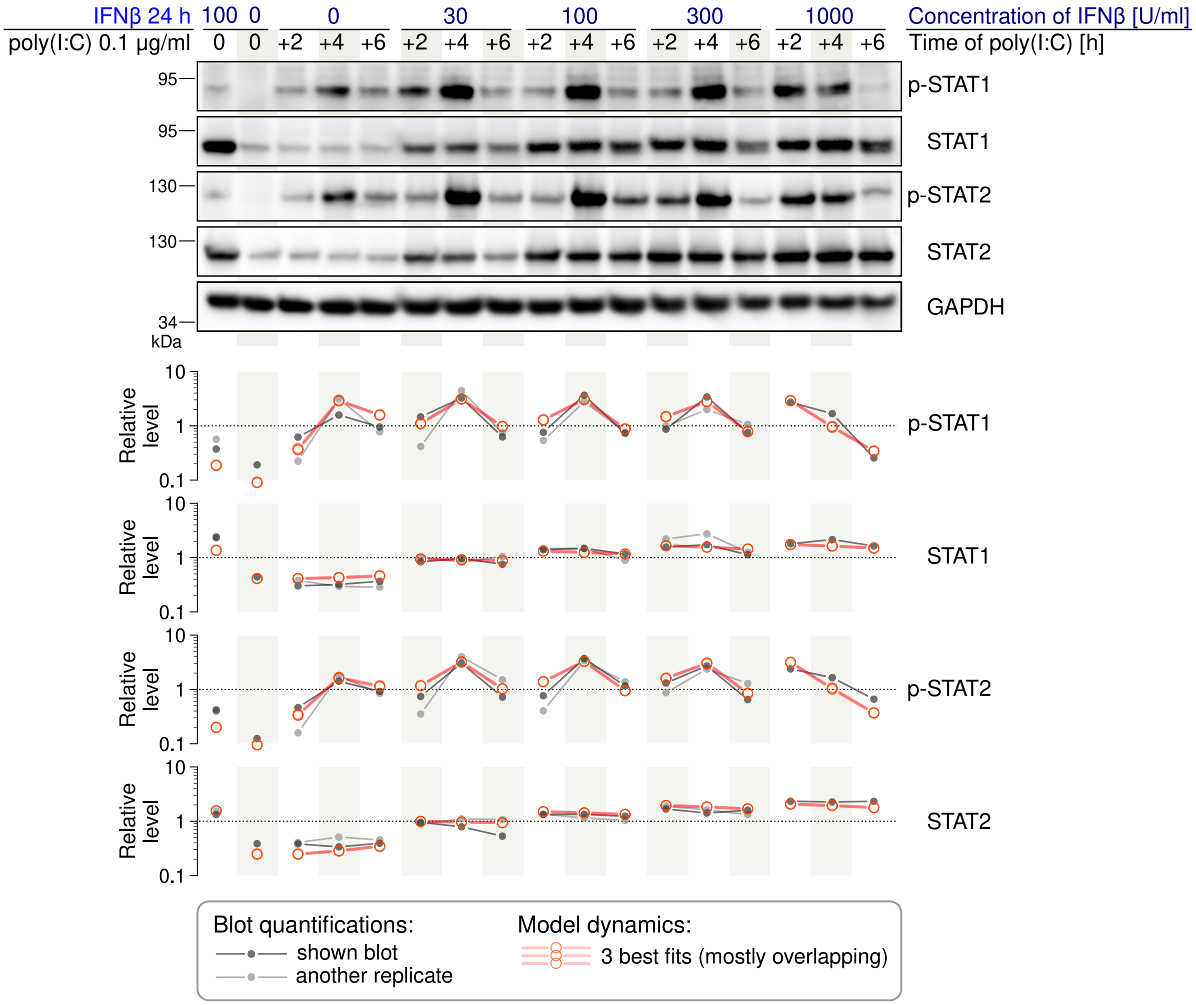

**Fig. S18**: **Poly(I:C)-induced activation of STAT1/2 in A549 WT cells prestimulated with IFNβ at various concentrations.** IFNβ-containing medium has not been displaced upon addition of poly(I:C). Representative blots from 2 independent experiments. Uncropped blots are available in Source Data.

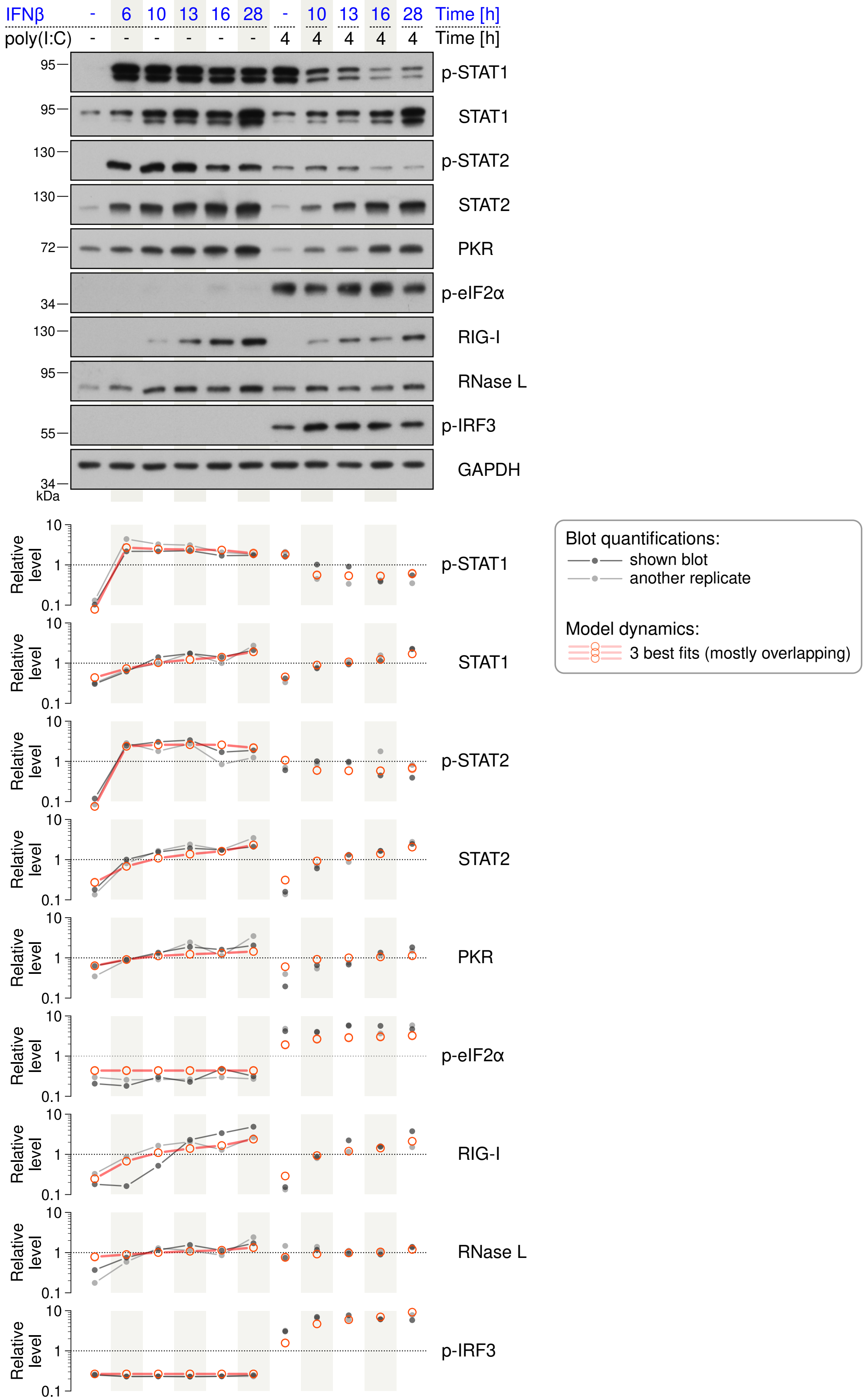

**Fig. S19**: **A549 WT cells prestimulated with IFNβ for different durations before stimulation with poly(I:C).** Concentrations: IFNβ 1000 U/ml, poly(I:C) 0.1 μg/ml. Representative blots from 2 independent experiments. Uncropped blots are available in Source Data.

**Table S1**: **Model parameters.** Kinetic rates are reported concisely according to the type of biochemical reaction: for reactions of type $\emptyset \xrightarrow{r} A,\ A \xrightarrow{r} \emptyset,\ A \xrightarrow{r} B,\ A+B \xrightarrow{r} C,\ C \xrightarrow{r} A+B$ just the rate $r$ is given, whereas for explicitly catalyzed reactions such as $E+S \xrightarrow{r} E+P,\ E \xrightarrow{r} E+P$ the rate is given as $[E] \times r$. Values of fitted parameters (dark gray) are given using 2 significant digits.

| **Process** | **Kinetic rate** | **Parameter values** |
|---|---|---|
| **Common parameters** | | |
| common rate of fast processes | k_FAST | k_FAST = 1 (assumed) |
| rate of transcription and/or transcript degradation | tg_TRANSCRIPT | tg_TRANSCRIPT = 0.000073 |
| rate of protein synthesis and/or protein degradation | sg_PROTEIN | sg_PROTEIN = 0.000015 |
| common rate for poly(I:C)-induced signaling processes | k_POLYIC | k_POLYIC = 0.00024 |
| inverse of a Michaelis–Menten-type constant in RNase L-driven transcript degradation | m_Rnasel | m_Rnasel = 0.031 |
| Michaelis–Menten constant in phospho-eIF2α-driven translation arrest | m_Eif2a | m_Eif2a = 0.016 |
| cytoplasm volume/nucleus volume | k_v | k_v = 5 (assumed) |

**The poly(I:C) module**

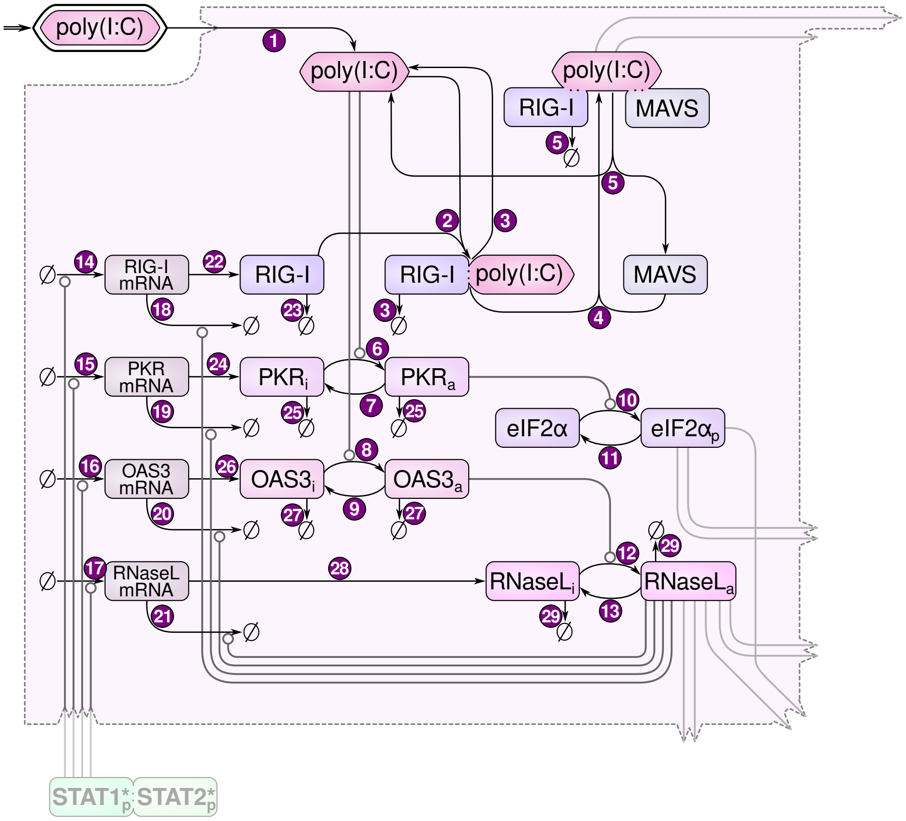

| | | |
|---|---|---|
| (1) internalization of poly(I:C) | i_Polyic | i_Polyic = k_POLYIC |

| | | |
|---|---|---|
| 2 binding of RIG-I and poly(I:C) | b_Rigi_Polyic | b_Rigi_Polyic = k_POLYIC |
| 3 RIG-I degradation in poly(I:C) : RIG-I | sg_Rigi | sg_Rigi = sg_PROTEIN |
| 4 binding of the poly(I:C) : RIG-I complex and MAVS | b_RigiPolyic_Mavs | b_RigiPolyic_Mavs = k_POLYIC |
| 5 RIG-I degradation in poly(I:C) : RIG-I : MAVS | sg_Rigi | sg_Rigi = sg_PROTEIN |
| 6 activation of PKR by poly(I:C) | a_Pkr_by_Polyic × [poly(I:C)] | a_Pkr_by_Polyic = k_POLYIC |
| 7 deactivation of PKR | d_Pkr | d_Pkr = k_POLYIC |
| 8 activation of OAS3 by poly(I:C) | a_Oas3_by_Polyic × [poly(I:C)] | a_Oas3_by_Polyic = k_POLYIC |
| 9 deactivation of OAS3 | d_Oas3 | d_Oas3 = k_POLYIC |
| 10 phosphorylation of eIF2α (basal and by active PKR) | p_Eif2a_basal + p_Eif2a_by_Pkr × $[\mathrm{PKR_a}]$ | p_Eif2a_basal = 0.0000077, p_Eif2a_by_Pkr = k_POLYIC |
| 11 dephosphorylation of phospho-eIF2α | q_Eif2a | q_Eif2a = k_POLYIC |
| 12 activation of RNase L by OAS3 | a_Rnasel_by_OAS3 × $[\mathrm{OAS3_a}]$ | a_Rnasel_by_OAS3 = k_POLYIC |
| 13 deactivation of RNase L | d_Rnasel | d_Rnasel = k_POLYIC |
| 14 RIG-I gene transcription | $\texttt{tg\_Isg\_mrna} \times \frac{\texttt{ma\_Rigi\_gene\_basal} + \texttt{a\_gene\_by\_Stat12dim} \times [\mathrm{STAT1_p : STAT2_p}]}{\texttt{ma\_Rigi\_gene\_basal} + \texttt{a\_gene\_by\_Stat12dim} \times [\mathrm{STAT1_p : STAT2_p}] + 1}$ | tg_Isg_mrna = tg_TRANSCRIPT, a_gene_by_Stat12dim = 13 000, ma_Rigi_gene_basal = 0.059, ma_Pkr_Oas3_gene_basal = 0.72, ma_Rnasel_gene_basal = 1.0, h_Pkr_gene = 1, h_Rnasel_gene = 1 |
| 15 PKR gene transcription | $\texttt{tg\_Isg\_mrna} \times \frac{\texttt{ma\_Pkr\_Oas3\_gene\_basal} + \texttt{a\_gene\_by\_Stat12dim} \times [\mathrm{STAT1_p : STAT2_p}]}{\texttt{ma\_Pkr\_Oas3\_gene\_basal} + \texttt{a\_gene\_by\_Stat12dim} \times [\mathrm{STAT1_p : STAT2_p}] + 1} \times \texttt{h\_Pkr\_gene}$ | |
| 16 OAS3 gene transcription | $\texttt{tg\_Isg\_mrna} \times \frac{\texttt{ma\_Pkr\_Oas3\_gene\_basal} + \texttt{a\_gene\_by\_Stat12dim} \times [\mathrm{STAT1_p : STAT2_p}]}{\texttt{ma\_Pkr\_Oas3\_gene\_basal} + \texttt{a\_gene\_by\_Stat12dim} \times [\mathrm{STAT1_p : STAT2_p}] + 1}$ | |
| 17 RNase L gene transcription | $\texttt{tg\_Isg\_mrna} \times \frac{\texttt{ma\_Rnasel\_gene\_basal} + \texttt{a\_gene\_by\_Stat12dim} \times [\mathrm{STAT1_p : STAT2_p}]}{\texttt{ma\_Rnasel\_gene\_basal} + \texttt{a\_gene\_by\_Stat12dim} \times [\mathrm{STAT1_p : STAT2_p}] + 1} \times \texttt{h\_Rnasel\_gene}$ | |
| 18 RIG-I mRNA degradation | $\texttt{tg\_Isg\_mrna} \times \frac{\texttt{m\_Rnasel} + [\mathrm{RNaseL_a}]}{\texttt{m\_Rnasel}}$ | tg_Isg_mrna = tg_TRANSCRIPT, m_Rnasel = 0.031 |
| 19 PKR mRNA degradation | | |
| 20 OAS3 mRNA degradation | | |
| 21 RNase L mRNA degradation | | |
| 22 RIG-I protein synthesis | sg_Rigi × [RIG-I mRNA] | sg_Rigi = sg_PROTEIN |
| 23 RIG-I protein degradation | sg_Rigi | |
| 24 PKR protein synthesis | sg_Pkr × [PKR mRNA] | sg_Pkr = 0.000043 |
| 25 PKR protein degradation | sg_Pkr | |
| 26 OAS3 protein synthesis | sg_Oas3 × [OAS3 mRNA] | sg_Oas3 = sg_PROTEIN |
| 27 OAS3 protein degradation | sg_Oas3 | |
| 28 RNase L protein synthesis | sg_Rnasel × [RNase L mRNA] | sg_Rnasel = sg_PROTEIN |
| 29 RNase L protein degradation | sg_Rnasel | |

### The NF-κB module

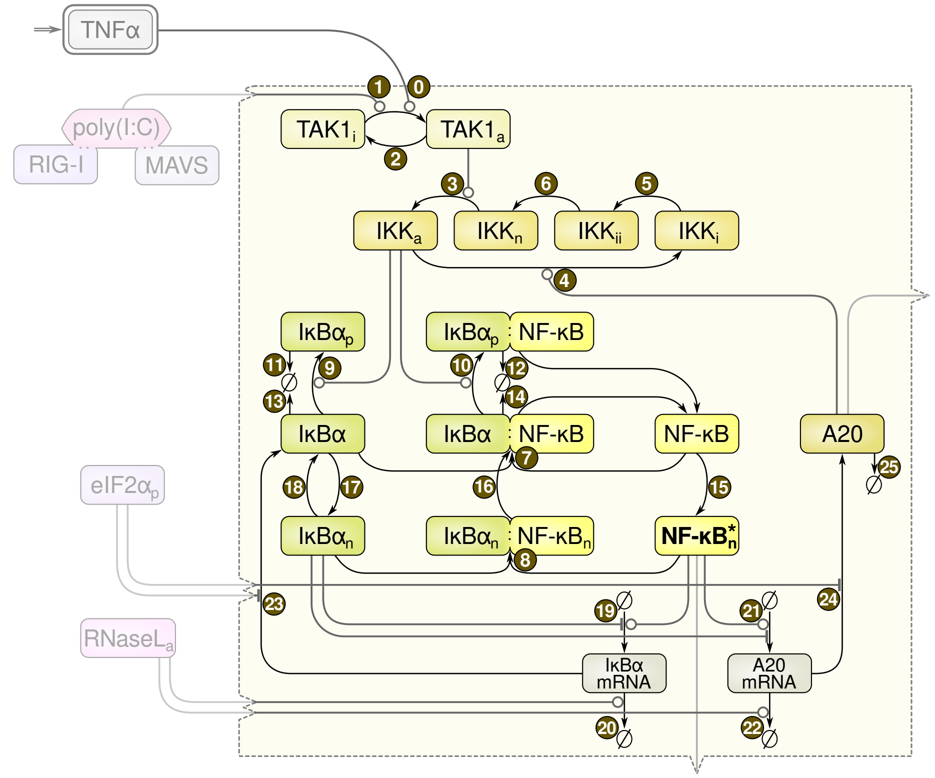

| | | |
|---|---|---|
| (0) activation of TAK1 by TNFα | a_Tak1_by_Tnfa × [TNFα] | a_Tak1_by_Tnfa = 0.89 |
| (1) activation of TAK1 by poly(I:C) : RIG-I : MAVS | a_Tak1_by_RigiMavs × [poly(I:C) : RIG-I : MAVS] | a_Tak1_by_RigiMavs = k_FAST |
| (2) deactivation of TAK1 | d_Tak1 | d_Tak1 = k_FAST |
| (3) activation of IKK by active TAK1 | a_Ikk × $[\mathrm{TAK1_a}]^2$ | a_Ikk = 0.011 |
| (4) deactivation of IKK | $\frac{\text{d\_Ikk\_1}}{\text{d\_Ikk\_2}} \times (\text{d\_Ikk\_2} + [\text{A20}])$ | d_Ikk_1 = 0.00088, d_Ikk_2 = 0.060 |
| (5), (6) transition of inactive IKK to(wards) the neutral state | d_Ikk_3 | d_Ikk_3 = 0.00027 |
| (7) binding of IκBα and NF-κB in the cytoplasm | b_Nfkb_Ikba_cyt | b_Nfkb_Ikba_cyt = k_FAST |
| (8) binding of IκBα and NF-κB in the nucleus | b_Nfkb_Ikba_nuc | b_Nfkb_Ikba_nuc = k_v × k_FAST |
| (9), (10) phosphorylation of IκBα by active IKK | p_Ikba_by_Ikk × $[\mathrm{IKK_a}]$ | p_Ikba_by_Ikk = 0.0084 |
| (11), (12) phospho-IκBα degradation | g_Ikba_p_any | g_Ikba_p_any = k_FAST |
| (13) nonbound nonphospho-IκBα degradation | g_Ikba_u_free | g_Ikba_u_free = sg_PROTEIN |
| (14) NF-κB-bound nonphospho-IκBα degradation | g_Ikba_u_with_Nfkb | g_Ikba_u_with_Nfkb = 0.000039 |
| (15) import of nonbound NF-κB into the nucleus | i_Nfkb | i_Nfkb = k_FAST |

| | | |
|---|---|---|
| 16 NF-κB : IκBα export from the nucleus | e_Nfkb_with_Ikba | e_Nfkb_with_Ikba = k_FAST |
| 17 nonbound nonphospho-IκBα import into the nucleus | i_Ikba | i_Ikba = 0.00056 |
| 18 nonbound nonphospho-IκBα export from the nucleus | e_Ikba | e_Ikba = 0.021 |
| 19 IκBα gene transcription | $\text{tg\_Ikba\_mrna} \times \frac{\text{a\_Ikba\_gene\_by\_Nfkb} \times [\text{NF-κB}_{\text{nuc free}}]}{\text{a\_Ikba\_gene\_by\_Nfkb} \times [\text{NF-κB}_{\text{nuc free}}] + \text{d\_Ikba\_gene\_by\_Ikba} \times [\text{IκBα}_{\text{nuc free}}] + \epsilon}$ | tg_Ikba_mrna = 0.00026, a_Ikba_gene_by_Nfkb = 0.068 × k_FAST, d_Ikba_gene_by_Ikba = k_FAST |
| 20 IκBα mRNA degradation | $\text{tg\_Ikba\_mrna} \times \frac{\text{m\_RnaseL} + [\text{RNaseL}_a]}{\text{m\_RnaseL}}$ | tg_Ikba_mrna = 0.00026, m_RnaseL = 0.031 |
| 21 A20 gene transcription | $\text{tg\_A20\_mrna} \times \frac{\text{a\_A20\_gene\_by\_Nfkb} \times [\text{NF-κB}_{\text{nuc free}}]}{\text{a\_A20\_gene\_by\_Nfkb} \times [\text{NF-κB}_{\text{nuc free}}] + \text{d\_A20\_gene\_by\_Ikba} \times [\text{IκBα}_{\text{nuc free}}] + \epsilon}$ | tg_A20_mrna = 0.00094, a_A20_gene_by_Nfkb = 0.026 × k_FAST, d_A20_gene_by_Ikba = k_FAST |
| 22 A20 mRNA degradation | $\text{tg\_A20\_mrna} \times \frac{\text{m\_RnaseL} + [\text{RNaseL}_a]}{\text{m\_RnaseL}}$ | tg_A20_mrna = 0.00094, m_RnaseL = 0.031 |
| 23 IκBα protein synthesis | $\text{s\_Ikba} \times [\text{IκBα mRNA}] \times \frac{\text{m\_eIF2a}}{\text{m\_eIF2a} + [\text{eIF2a}_p]}$ | s_Ikba = 0.0043, m_Eif2a = 0.016 |
| 24 A20 protein synthesis | $\text{sg\_A20\_mrna} \times [\text{A20 mRNA}] \times \frac{\text{m\_eIF2a}}{\text{m\_eIF2a} + [\text{eIF2a}_p]}$ | sg_A20_mrna = 0.0000050, m_Eif2a = 0.016 |
| 25 A20 protein degradation | sg_A20_mrna | sg_A20 = 0.0000050 |

### The IRF3 module

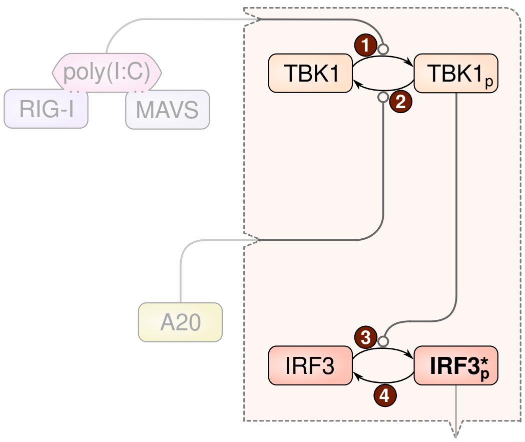

| | | |
|---|---|---|
| 1 TBK1 phosphorylation induced by poly(I:C) : RIG-I : MAVS | p_Tbk1_by_RigiMavs × [poly(I:C) : RIG-I : MAVS] | p_Tbk1_by_RigiMavs = k_FAST |
| 2 deactivation of TBK1 | q_Tbk1 + q_Tbk1_by_A20 × [A20] | q_Tbk1 = k_FAST, q_Tbk1_by_A20 = 390 × k_FAST |
| 3 activation of IRF3 by active TBK1 | $\text{p\_Irf3\_by\_Tbk1} \times [\text{TBK1}_p]$ | p_Irf3_by_Tbk1 = k_POLYIC |
| 4 deactivation of IRF3 | q_Irf3 | q_Irf3 = k_POLYIC |

### The IFNβ module

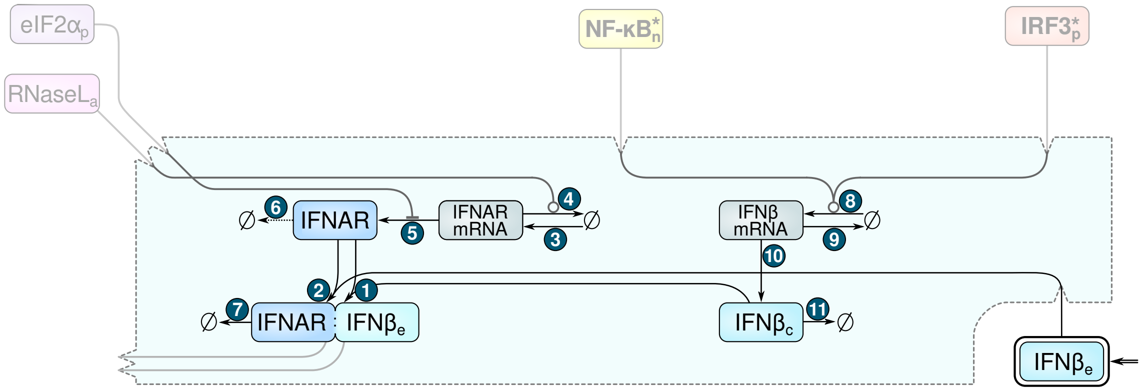

| | | |
|---|---|---|
| ① binding of IFNβ & IFNAR (autocrine) | b_Ifnar_Ifnb_cyt | b_Ifnar_Ifnb_cyt = k_FAST |
| ② binding of (external) IFNβ and IFNAR | b_Ifnar_Ifnb_ext | b_Ifnar_Ifnb_ext = 0.00075 |
| ③ IFNAR gene transcription | tg_Ifnar_mrna | tg_Ifnar_mrna = tg_TRANSCRIPT |
| ④ IFNAR mRNA degradation | $\texttt{tg\_Ifnar\_mrna} \times \frac{\texttt{m\_Rnasel\_Ifnar\_mrna} + [\text{RNaseL}_a]}{\texttt{m\_Rnasel\_Ifnar\_mrna}}$ | tg_Ifnar_mrna = tg_TRANSCRIPT, m_Rnasel_Ifnar_mrna = 0.0039 |
| ⑤ IFNAR protein synthesis | $\texttt{s\_Ifnar} \times [\text{IFNAR mRNA}] \times \frac{\texttt{m\_eIF2a}}{\texttt{m\_eIF2a} + [\text{eIF2a}_p]}$ | s_Ifnar = 0.000029, m_Eif2a = 0.016 |
| ⑥ unbound IFNAR degradation | g_Ifnar | g_Ifnar = 0.000098 |
| ⑦ IFNβ-bound IFNAR degradation | g_Ifnar_w_Ifnb | g_Ifnar_w_Ifnb = 0.00031 |
| ⑧ IFNβ gene transcription induced by NF-κB$_{\text{nuc free}}$ and IRF3$_p$ | $\texttt{tg\_Ifnb\_mrna} \times \frac{[\text{NF-κB}_{\text{nuc free}}] \times [\text{IRF3}_p]}{[\text{NF-κB}_{\text{nuc free}}] \times [\text{IRF3}_p] + \texttt{m\_Ifnb\_mrna\_NfkbIrf3}}$ | tg_Ifnb_mrna = tg_TRANSCRIPT, m_Ifnb_mrna_NfkbIrf3 = 0.000032 |
| ⑨ IFNβ mRNA degradation | tg_Ifnb_mrna | tg_Ifnb_mrna = tg_TRANSCRIPT |
| ⑩ IFNβ protein synthesis<br>⑪ IFNβ protein degradation | sg_Ifnb | sg_Ifnb = sg_PROTEIN |

### The STAT1/2 module

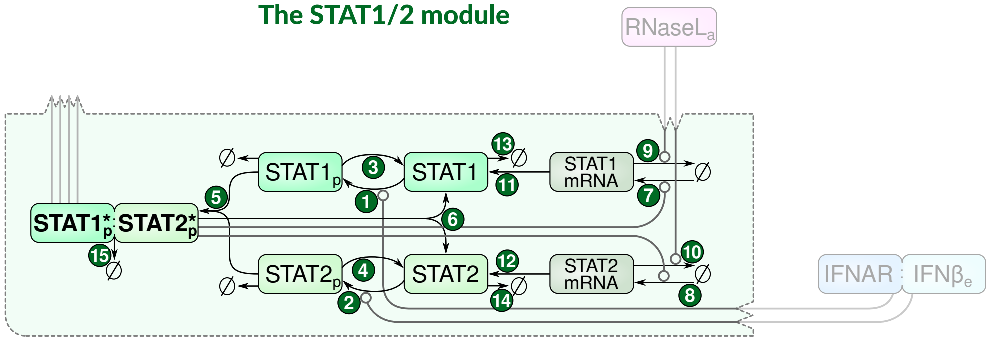

| | | |
|---|---|---|
| ①, ② STAT1/2 phosphorylation by IFNβ-activated IFNAR | $\texttt{p\_Stat} \times [\text{IFNAR}_a] \times \frac{\texttt{m\_Ifnar\_a}}{\texttt{m\_Ifnar\_a} + [\text{STAT1}_u]}$ | p_Stat = k_FAST, m_Ifnar_a = 0.020 |
| ③, ④ STAT1/2 dephosphorylation | q_Stat | q_Stat = 0.0018 |

| | | |
|---|---|---|
| (5) heterodimerization of phospho-STAT1 & phospho-STAT2 | b_Stat1_Stat2 | b_Stat1_Stat2 = k_FAST |
| (6) dephosphorylation & unbinding of p-STAT1 : p-STAT2 dimer | qu_Stat1_Stat2 | qu_Stat1_Stat2 = 0.052 |
| (7) STAT1 gene transcription | $\texttt{tg\_Stat\_mrna} \times \frac{\texttt{ma\_Stat1\_gene\_basal} + \texttt{a\_gene\_by\_Stat12dim} \times [\mathrm{STAT1_p : STAT2_p}]}{\texttt{ma\_Stat1\_gene\_basal} + \texttt{a\_gene\_by\_Stat12dim} \times [\mathrm{STAT1_p : STAT2_p}] + 1}$ | tg_Stat_mrna = tg_TRANSCRIPT, ma_Stat1_gene_basal = 0.18, a_gene_by_Stat12dim = 13 000 |
| (8) STAT2 gene transcription | $\texttt{tg\_Stat\_mrna} \times \frac{\texttt{ma\_Stat2\_gene\_basal} + \texttt{a\_gene\_by\_Stat12dim} \times [\mathrm{STAT1_p : STAT2_p}]}{\texttt{ma\_Stat2\_gene\_basal} + \texttt{a\_gene\_by\_Stat12dim} \times [\mathrm{STAT1_p : STAT2_p}] + 1}$ | tg_Stat_mrna = tg_TRANSCRIPT, ma_Stat2_gene_basal = 0.070, a_gene_by_Stat12dim = 13 000 |
| (9), (10) STAT1/2 mRNA degradation | $\texttt{tg\_Stat\_mrna} \times \frac{\texttt{m\_RnaseL} + [\mathrm{RNaseL_a}]}{\texttt{m\_RnaseL}}$ | tg_Stat_mrna = tg_TRANSCRIPT, m_RnaseL = 0.031 |
| (11), (12) STAT1/2 protein synthesis | sg_Stat | sg_Stat = sg_PROTEIN |
| (13), (14) STAT1/2 monomer degradation | | |
| (15) p-STAT1 : p-STAT2 dimer degradation | | |

**Table S2**: **Reagents, antibodies, and other resources.**

| Reagent/resource | Source or reference | Catalog #, RRID, DOI |
|---|---|---|
| ***Experimental models*** | | |
| *Homo sapiens*: A549 WT cell line | ATCC | CCL-185 |
| *Homo sapiens*: A549 *IRF3* KO cell line | This paper. | N/A |
| *Homo sapiens*: A549 *TNFAIP3* (A20) KO cell line | This paper. | N/A |
| *Homo sapiens*: A549 *MAVS* KO cell line | This paper. | N/A |
| *Homo sapiens*: A549 *EIF2AK2* (PKR) KO cell line | This paper. | N/A |
| *Homo sapiens*: A549 *RNASEL* (RNase L) KO cell line | Liu & Moss (2016) *J. Virol.* | 10.1128/jvi.00869-16 |
| *Homo sapiens*: A549 *EIF2AK2* (PKR) & *RNASEL* (RNase L) double KO (dKO) cell line | Liu & Moss (2016) *J. Virol.* | 10.1128/jvi.00869-16 |
| *Homo sapiens*: BEAS-2B | ATCC | CRL-9609 |
| *Homo sapiens*: HeLa cell line | ATCC | CCL-2 |
| *Canis familiaris*: MDCK (NBL-2) | ATCC | CCL-34 |
| ***Recombinant DNA*** | | |
| pCMV-Cas9-GFP (*IRF3* U6gRNA) | Sigma-Aldrich | HS 0000187830 |
| CMV-CAS9-GFP (*TNFAIP3* U6gRNA) | Sigma-Aldrich | HS 0000224185 |
| pCMV-Cas9-RFP (*EIF2AK2* U6gRNA) | Sigma-Aldrich | HS 0000128652 |
| pSpCas9(BB)-2A-GFP (PX458) | Addgene | 48138 |
| ***Viral strains*** | | |
| Human Respiratory Syncytial Virus; strain A2 | ATCC | VR-1540 |
| Influenza A virus (H1N1); A/PR/8/34; TC adapted | ATCC | VR-1469 |
| ***Antibodies*** [with dilutions for Western blot (WB) or immunofluorescence (IF)] | | |
| Rabbit monoclonal anti-NF-κB p65 (D14E12) XP [WB 1:1000, IF 1:1000] | Cell Signaling Technology | 8242, AB_10859369 |
| Rabbit monoclonal anti-phospho-IRF-3 $Ser^{396}$ (4D4G) [WB 1:1000] | Cell Signaling Technology | 4947, AB_823547 |
| Rabbit monoclonal anti-IRF-3 (D83B9) [WB 1:1000] | Cell Signaling Technology | 4302, AB_1904036 |
| Mouse monoclonal anti-IκBα (L35A5) [WB 1:1000] | Cell Signaling Technology | 4814, AB_390781 |
| Rabbit monoclonal anti-RIG-I (D14G6) [WB 1:1000] | Cell Signaling Technology | 3743, AB_2269233 |
| Rabbit monoclonal anti-phospho-STAT1 $Tyr^{701}$ (58D6) [WB 1:1000, IF 1:200] | Cell Signaling Technology | 9167, AB_561284 |
| Rabbit monoclonal anti-A20/TNFAIP3 (D13H3) [WB 1:1000] | Cell Signaling Technology | 5630, AB_10698880 |
| Mouse monoclonal anti-HDAC1 [WB 1:1000] | Cell Signaling Technology | 5356, AB_10612242 |
| Rabbit polyclonal anti-MAVS [WB 1:1000] | Cell Signaling Technology | 3993, AB_823565 |
| Rabbit monoclonal anit-phospho-Stat2 ($Tyr^{690}$) (D3P2P) [WB 1:1000] | Cell Signaling Technology | 88410, AB_2800123 |
| Rabbit monoclonal anti-Stat2 (D9J7L) [WB 1:1000] | Cell Signaling Technology | 72604, AB_2799824 |
| Rabbit polyclonal anti-phospho-eIF2α ($Ser^{51}$) [WB 1:1000] | Cell Signaling Technology | 9721, AB_330951 |
| Mouse monoclonal anti-eIF2α (L57A5) [WB 1:1000] | Cell Signaling Technology | 2103, AB_836874 |

| Reagent/resource | Source or reference | Catalog #, RRID, DOI |
|---|---|---|
| Rabbit monoclonal anti-eIF2α [WB 1:1000] | Cell Signaling Technology | 9722, AB_2230924 |
| Rabbit monoclonal anti-phospho-IKKα/β (Ser176/180) (16A6) [WB 1:1000] | Cell Signaling Technology | 2697, AB_2079382 |
| Mouse monoclonal anti-IRF-3 (D-3) [IF 1:500] | Santa Cruz Biotechnology | sc-376455, AB_11151578 |
| Mouse monoclonal anti-PKR (B-10) [WB 1:1000] | Santa Cruz Biotechnology | sc-6282, AB_628150 |
| Mouse monoclonal anti-RNase L (E-9) [WB 1:1000] | Santa Cruz Biotechnology | sc-74405, AB_2181661 |
| Mouse monoclonal anti-OAS1 (F-3) [WB 1:1000] | Santa Cruz Biotechnology | sc-374656, AB_10991311 |
| Goat polyclonal anti-Influenza A Virus [IF 1:1000] | Abcam | ab20841, AB_775660 |
| Mouse monoclonal anti-Influenza A Virus (C43) [IF for virus quantification 1:1000, WB 1:1000] | Abcam | ab128193, AB_11143769 |
| Goat polyclonal anti-Respiratory Syncytial Virus [IF 1:1000] | Abcam | ab20745, AB_777677 |
| Mouse monoclonal anti-RSV fusion glycoprotein [IF for virus quantification 1:1000, WB 1:1000] | Abcam | ab43812, AB__777676 |
| Rabbit monoclonal Recombinant Anti-Interferon α/β receptor 1 [WB 1:1000] | Abcam | ab124764, AB_10972855 |
| Rabbit polyclonal anti-IFNGR1 [WB 1:1000] | Cell Signaling | 34808, AB_2799061 |
| Goat polyclonal anti-STAT2 [WB 1:1000] | R&D Systems | PAF-ST2, AB_2240134 |
| Mouse monoclonal anti-STAT1 [WB 1:1000] | BD Biosciences | 610116, AB_397522 |
| Mouse monoclonal anti-GAPDH (6C5) [WB 1:10 000] | Merck | MAB374, AB_2107445 |
| hFAB™ Rhodamine anti-GAPDH Primary Antibody [WB 1:10 000] | Bio-Rad | 12004168 |
| StarBright Blue 700 Goat Anti-Mouse IgG [WB 1:10 000] | Bio-Rad | 12004159, AB_2884948 |
| OAS2 Monoclonal Antibody (OTI6E6) [WB 1:1000] | Thermo Fisher Scientific | MA5-26552, AB_2724898 |
| OAS3 Polyclonal Antibody [WB 1:1000] | Thermo Fisher Scientific | PA5-59539, AB_2644933 |
| Donkey polyclonal anti-rabbit IgG (H+L), Alexa Fluor® 488 conjugate [IF 1:1000] | Thermo Fisher Scientific | A-21206, AB_141708 |
| Donkey polyclonal anti-mouse IgG (H+L), Alexa Fluor® 555 conjugate [IF 1:1000] | Thermo Fisher Scientific | A-31570, AB_2536180 |
| Donkey polyclonal anti-goat IgG (H+L), Alexa Fluor® 633 conjugate [IF 1:1000] | Thermo Fisher Scientific | A-21082, AB_141493 |
| Goat anti-Rabbit IgG (H+L) Secondary Antibody, DyLight 800 4X PEG [WB 1:10 000] | Thermo Fisher Scientific | SA5-35571, AB_2556775 |
| Goat anti-Mouse IgG (H+L) Secondary Antibody, DyLight 800 4X PEG [WB 1:10 000] | Thermo Fisher Scientific | SA5-35521, AB_2556774 |
| Goat polyclonal anti-rabbit immunoglobulins/HRP [WB 1:10 000] | Agilent | P0448, AB_2617138 |
| Goat anti-mouse immunoglobulins/HRP [WB 1:10 000] | Agilent | P0447, AB_2617137 |
| Polyclonal Rabbit Anti-Goat Immunoglobulins/HRP [WB 1:10 000] | Agilent | P0449, AB_2617143 |
| ***Oligonucleotides and other sequence-based reagents: gRNA*** | | |
| gRNA targeting AATACATACCGTCAGAAGCAGG in *EIF2AK2* | This paper. | N/A |
| gRNA targeting GTAGATACAACTGACCCTGT in *MAVS* | This paper. | N/A |
| gRNA targeting ACACCTCTCCGGACACCAATGG in *IRF3* | This paper. | N/A |
| gRNA targeting GTCATAGCCGAGAACAATGGGG in *TNFAIP3* | This paper. | N/A |

| Reagent/resource | Source or reference | Catalog #, RRID, DOI |
|---|---|---|
| ***Oligonucleotides and other sequence-based reagents: TaqMan Gene Expression Assays*** | | |
| Hs01066116_m1: *IFNAR1* (single tube) | Thermo Fisher Scientific | 4331182 |
| Hs00988304_m1: *IFNGR1* (single tube) | Thermo Fisher Scientific | 4331182 |
| Hs00153283_m1 *NFKBIA* (array card) | Thermo Fisher Scientific | 4342249 |
| Hs00234713_m1 *TNFAIP3* (array card) | Thermo Fisher Scientific | 4342249 |
| Hs00985639_m1 *IL6* (array card) | Thermo Fisher Scientific | 4342249 |
| Hs00174103_m1 *CXCL8* (array card) | Thermo Fisher Scientific | 4342249 |
| Hs00705164_s1 *SOCS1* (array card) | Thermo Fisher Scientific | 4342249 |
| Hs01077958_s1 *IFNB1* (array card) | Thermo Fisher Scientific | 4342249 |
| Hs00533665_m1 *IFIT2* (array card) | Thermo Fisher Scientific | 4342249 |
| Hs00982282_m1 *CCL5* (array card) | Thermo Fisher Scientific | 4342249 |
| Hs01013996_m1 *STAT1* (array card) | Thermo Fisher Scientific | 4342249 |
| Hs00169345_m1 *EIF2AK2* (array card) | Thermo Fisher Scientific | 4342249 |
| Hs01013123_m1 *STAT2* (array card) | Thermo Fisher Scientific | 4342249 |
| Hs00973637_m1 *OAS1* (array card) | Thermo Fisher Scientific | 4342249 |
| Hs01061436_m1 *DDX58* (array card) | Thermo Fisher Scientific | 4342249 |
| Hs01921425_s1 *ISG15* (array card) | Thermo Fisher Scientific | 4342249 |
| Hs00223420_m1 *IFIH1* (array card) | Thermo Fisher Scientific | 4342249 |
| Hs01675197_m1 *IFIT1* (array card) | Thermo Fisher Scientific | 4342249 |
| Hs00971960_m1 *IRF1* (array card) | Thermo Fisher Scientific | 4342249 |
| Hs01547283_m1 *IRF3* (array card) | Thermo Fisher Scientific | 4342249 |
| Hs00158114_m1 *IRF5* (array card) | Thermo Fisher Scientific | 4342249 |
| Hs00185375_m1 *IRF7* (array card) | Thermo Fisher Scientific | 4342249 |
| Hs99999905_m1 *GAPDH* (array card) | Thermo Fisher Scientific | 4342249 |
| Hs99999909_m1 *HPRT1* (array card) | Thermo Fisher Scientific | 4342249 |
| Hs99999901_s1 18S rRNA (array card) | Thermo Fisher Scientific | 4342249 |
| ***Chemicals, enzymes and other reagents*** | | |
| Polyinosinic–polycytidylic acid potassium salt | Sigma-Aldrich | P9582 |
| Human Interferon α | PeproTech | 300-02AA |
| Human Interferon β 1a | PBL Assay Science | 11415-1 |
| IFN-γ Recombinant Human Protein | Thermo Fisher Scientific | PHC4031 |
| eBioscience Brefeldin A Solution (1000X) | Thermo Fisher Scientific | 00-4506-51 |
| DAPI | Sigma-Aldrich | D8417 |
| Mowiol 4-88 | Sigma-Aldrich | 81381 |
| ***Other*** | | |
| RNA Nano Chips | Agilent Technologies | 5067-1521 |
| PureLink RNA Mini Kit | Thermo Fisher Scientific | 12183020 |
| High-Capacity cDNA Reverse Transcription Kit with RNase Inhibitor | Thermo Fisher Scientific | 4374966 |

| Reagent/resource | Source or reference | Catalog #, RRID, DOI |
| --- | --- | --- |
| ***Other** – continued* | | |
| VeriKine Human Interferon-Beta ELISA Kit | PBL Assay Science | 41410 |
| Lipofectamine LTX Reagent with PLUS Reagent | Thermo Fisher Scientific | 15338100 |
| LookOut Mycoplasma qPCR Detection Kit | Merck | 200-664-3 |
| ***Equipment*** | | |
| Agilent 2100 Bioanalyzer | Agilent Technologies | G2939BA |
| ChemiDoc MP Imaging System | Bio-Rad | 12003154 |
| Nucleofector 2b Device | Lonza | AAB-1001 |
| Multiskan GO plate reader | Thermo Fisher Scientific | |
| QuantStudio 3D Digital PCR System | Thermo Fisher Scientific | |
| QuantStudio 12K Flex Real-Time PCR System with the Array Card block | Life Technologies | |
| Leica TCS SP5 | Leica Microsystems | |
| ***Software*** | | |
| BIONETGEN | Harris *et al.* (2016) *Bioinformatics* | 10.1093/bioinformatics/btw469 |
| RULEBENDER | Smith *et al.* (2012) *BMC Bioinformatics* | 10.1186/1471-2105-13-S8-S3 |
| PYBIONETFIT | Mitra *et al.* (2019) *iScience* | 10.1016/j.isci.2019.08.045 |
