## Supplementary material for "Non-self RNA rewires IFNβ signaling: A mathematical model of the innate immune response": Source Data: Uncropped Western blot images and uncropped full-resolution immunostaining images.

featuring the article

### Contents

**Source Data @ Figure 1**
Uncropped full-resolution microscopy images for Figures 1A and 1B.

**Source Data @ Figure 2**
Uncropped blots for Figures 2A, 2B, 2E, 2F.
Uncropped full-resolution microscopy images for Figures 2C and 2D.

**Source Data @ Figure 3**
Uncropped blots for Figures 3A, 3B, 3F.
Uncropped full-resolution microscopy images for Figure 3C.

**Source Data @ Figure 4**
Uncropped blots for Figure 4A.

**Source Data @ Figure S1**
Uncropped blots for Figure S1A.

**Source Data @ Figure S2**
Uncropped blots for Figures S2A, S2B, S2C.

**Source Data @ Figure S5**
Uncropped blots.

**Source Data @ Figure S7**
Uncropped blots for Figure S7A.

**Source Data @ Figure S8**
Uncropped blots for Figure S8A, S8B.

**Source Data @ Figure S9**
Uncropped blots.

**Source Data @ Figure S11**
Uncropped blots @ Figure S11A.

**Source Data @ Figure S12**
Uncropped blots.

**Source Data @ Figure S17**
Uncropped blots.

**Source Data @ Figure S18**
Uncropped blots.

**Source Data @ Figure S19**
Uncropped blots.

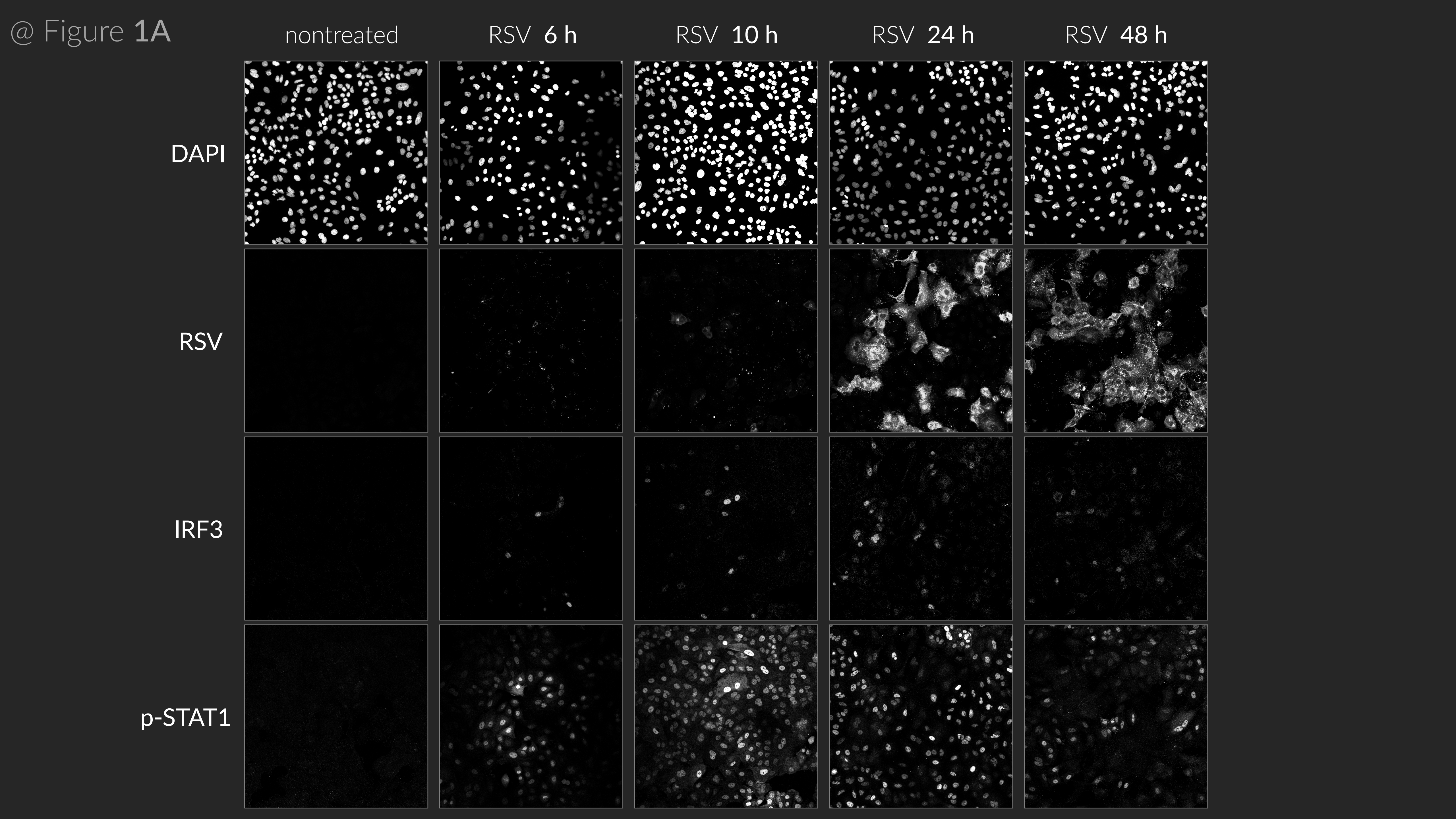
@ Figure 1A
nontreated
RSV 6 h
RSV 10 h
RSV 24 h
RSV 48 h
DAPI
RSV
IRF3
p-STAT1

@ Figure 1B

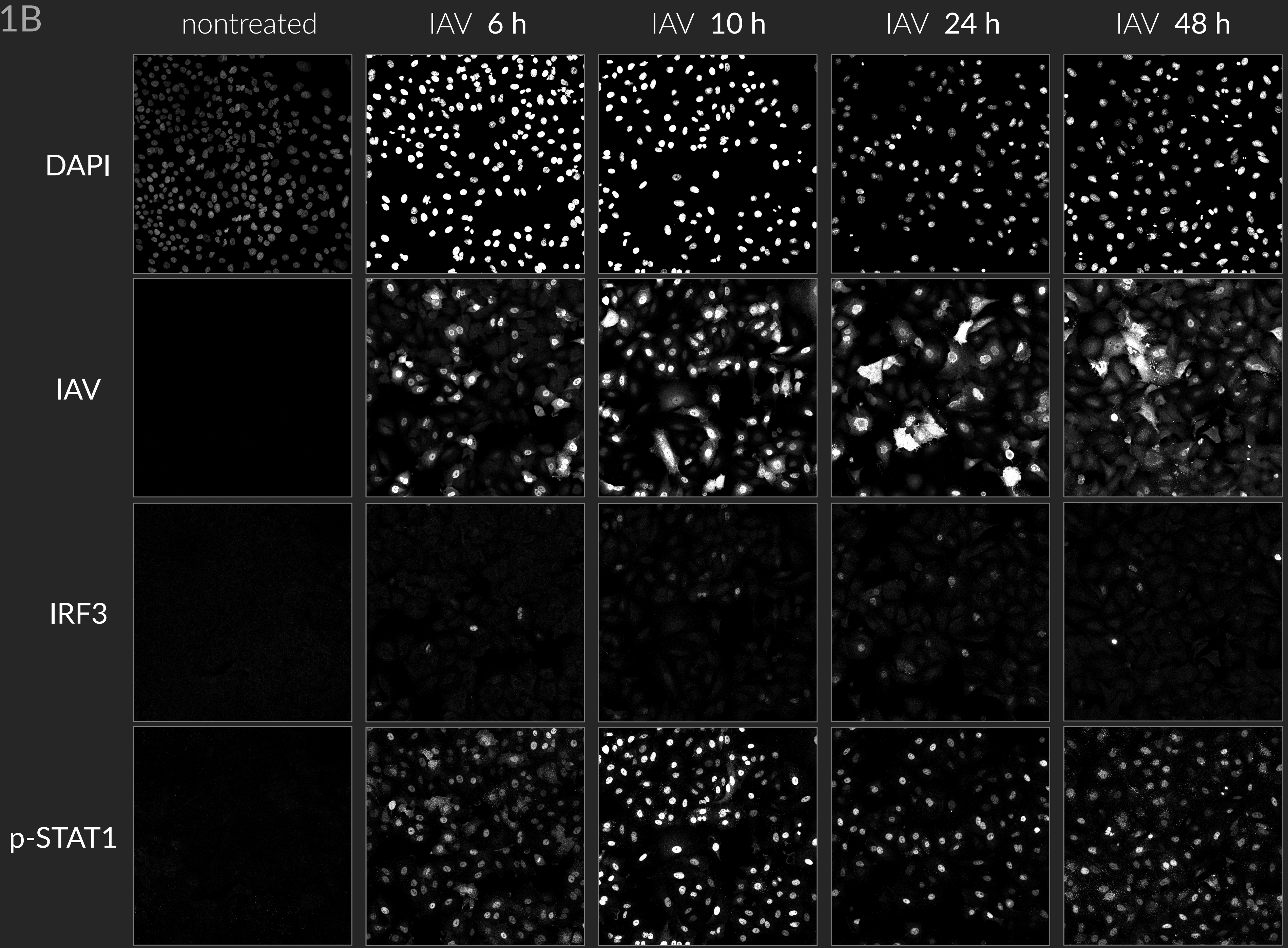

nontreated
IAV 6 h
IAV 10 h
IAV 24 h
IAV 48 h
DAPI
IAV
IRF3
p-STAT1

@ Figure **2A**

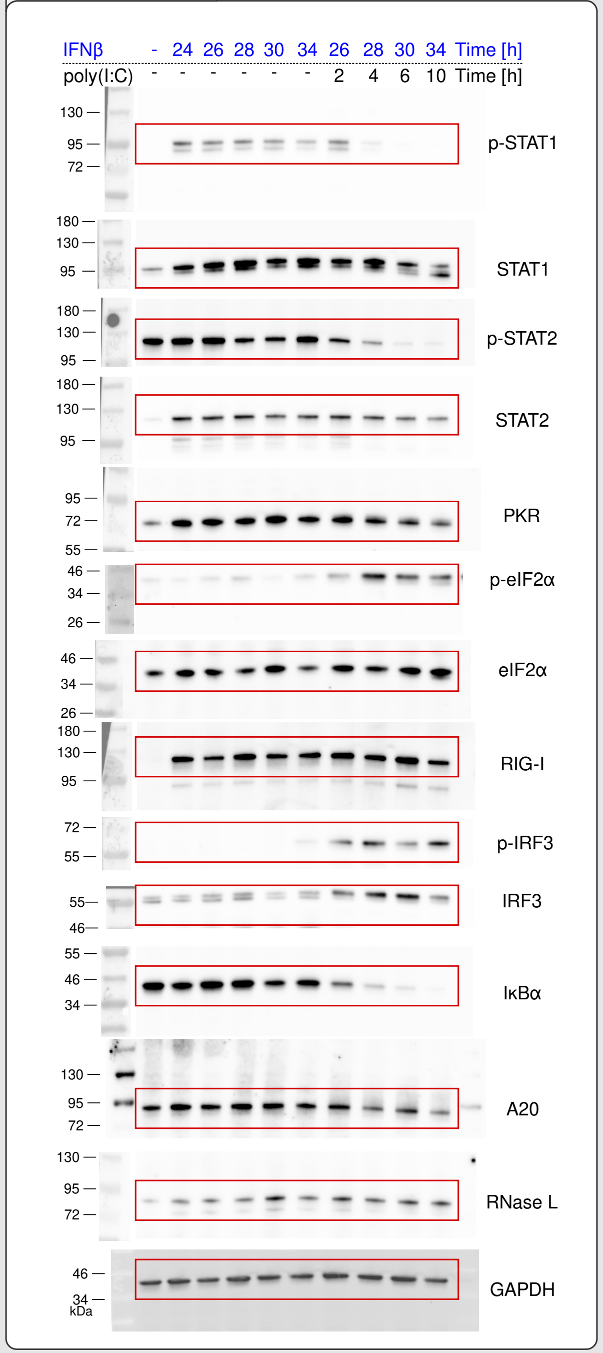


@ Figure **2B**

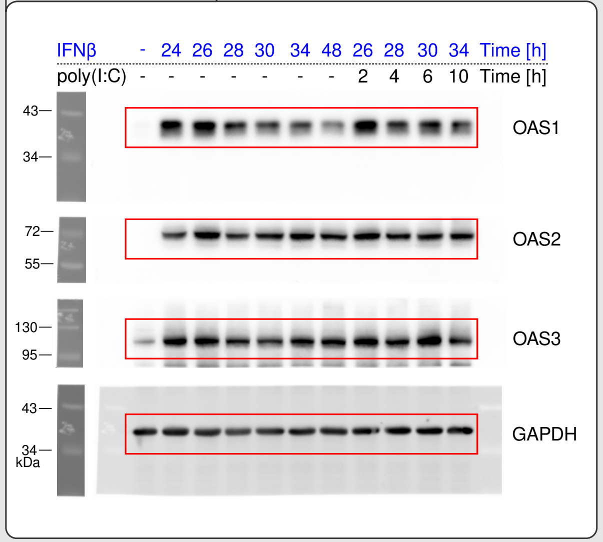


@ Figure **2E**

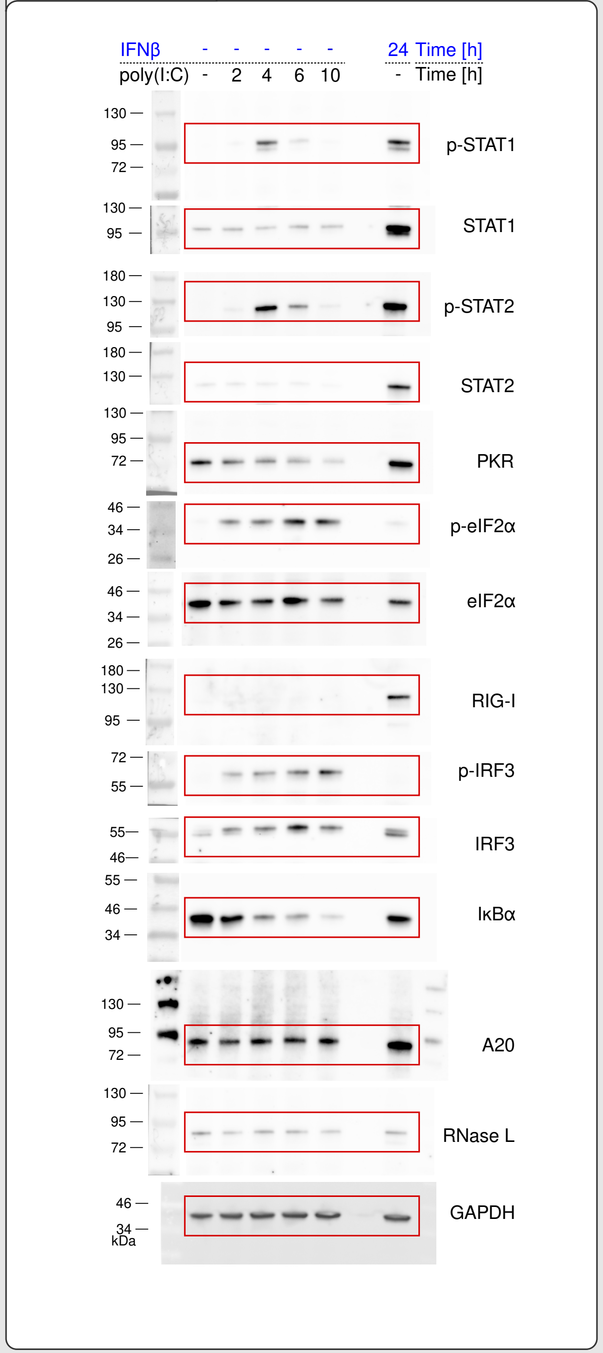


@ Figure **2F**

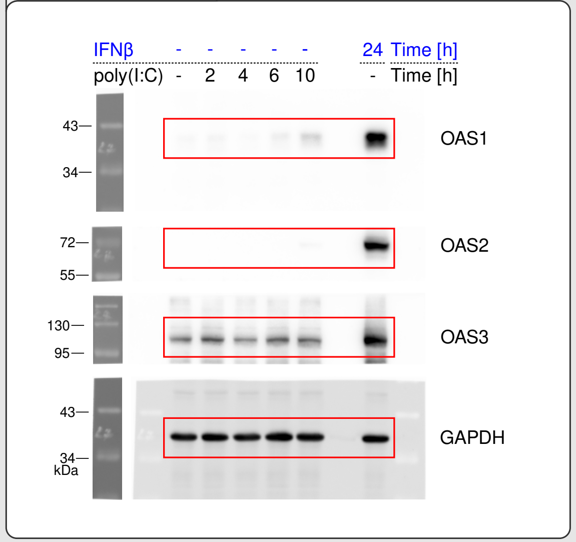

@ Figure **2C**

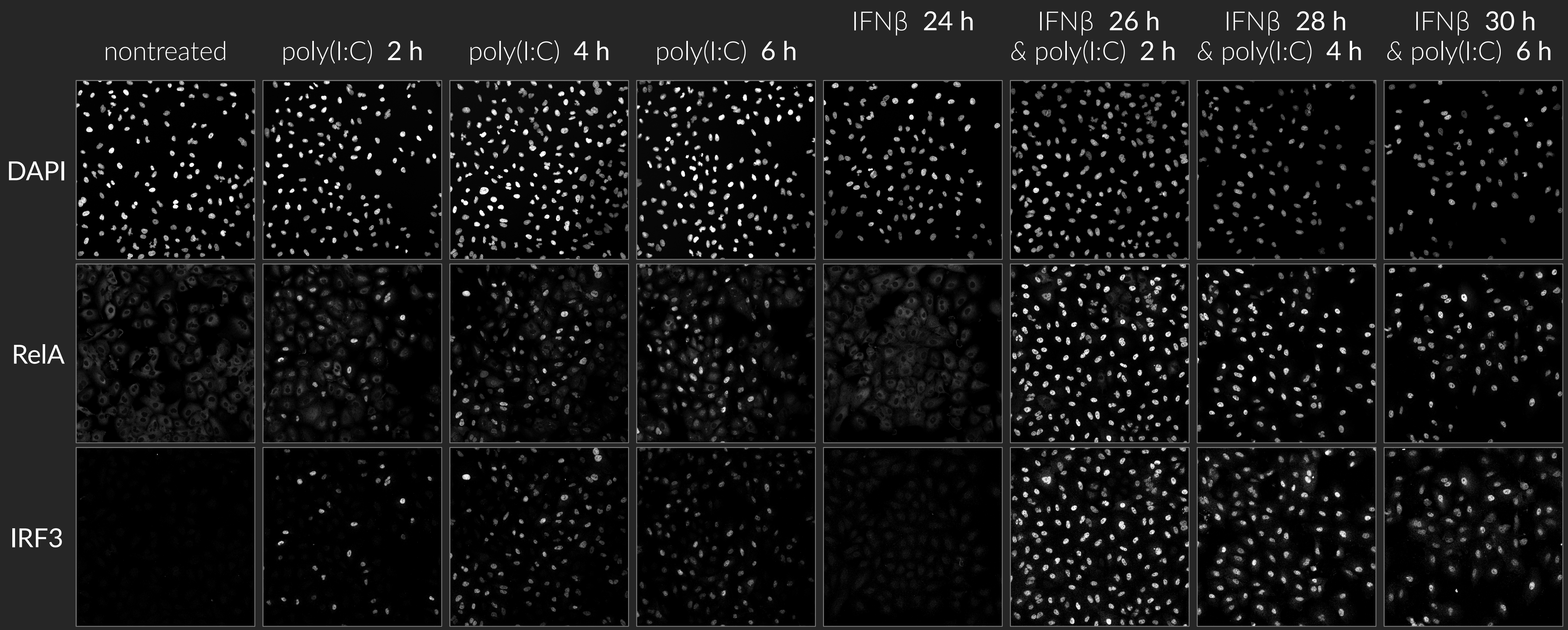
nontreated
poly(I:C) 2 h
poly(I:C) 4 h
poly(I:C) 6 h
IFNβ 24 h
IFNβ 26 h & poly(I:C) 2 h
IFNβ 28 h & poly(I:C) 4 h
IFNβ 30 h & poly(I:C) 6 h
DAPI
RelA
IRF3

@ Figure 2D

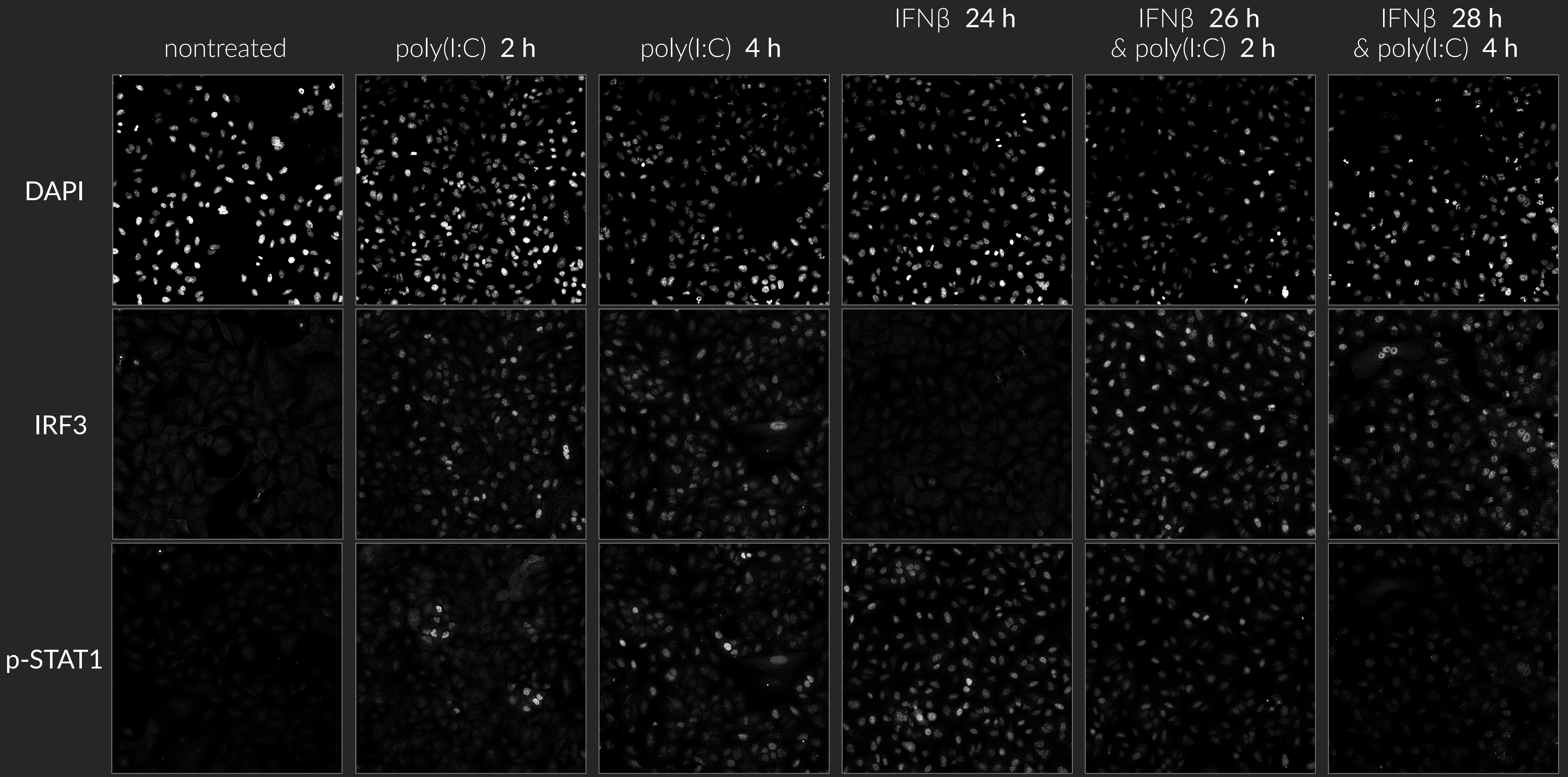
nontreated
poly(I:C) 2 h
poly(I:C) 4 h
IFNβ 24 h
IFNβ 26 h
& poly(I:C) 2 h
IFNβ 28 h
& poly(I:C) 4 h
DAPI
IRF3
p-STAT1

@ Figure **3A**

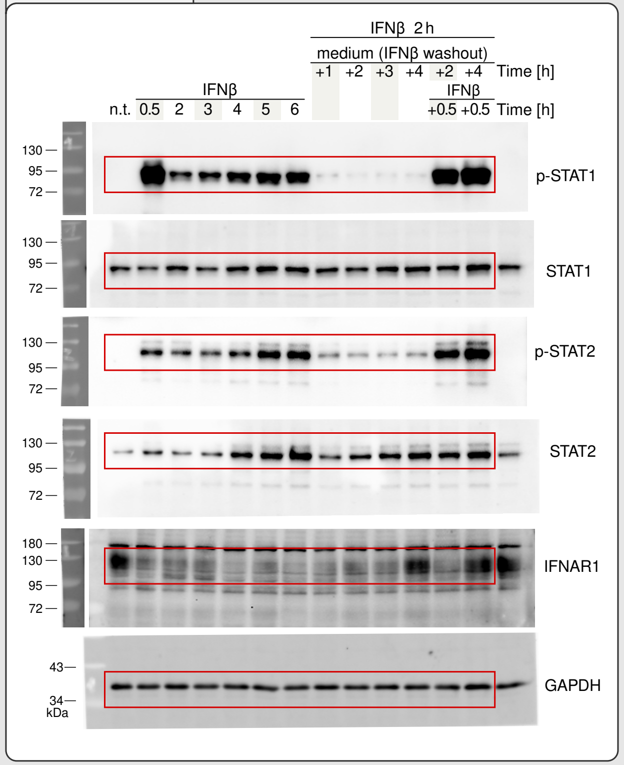


@ Figure **3B**

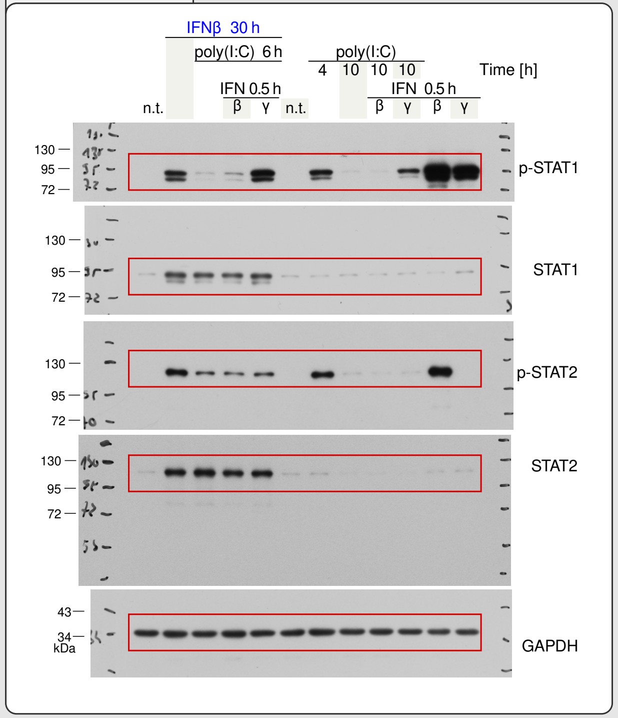


@ Figure **3F**

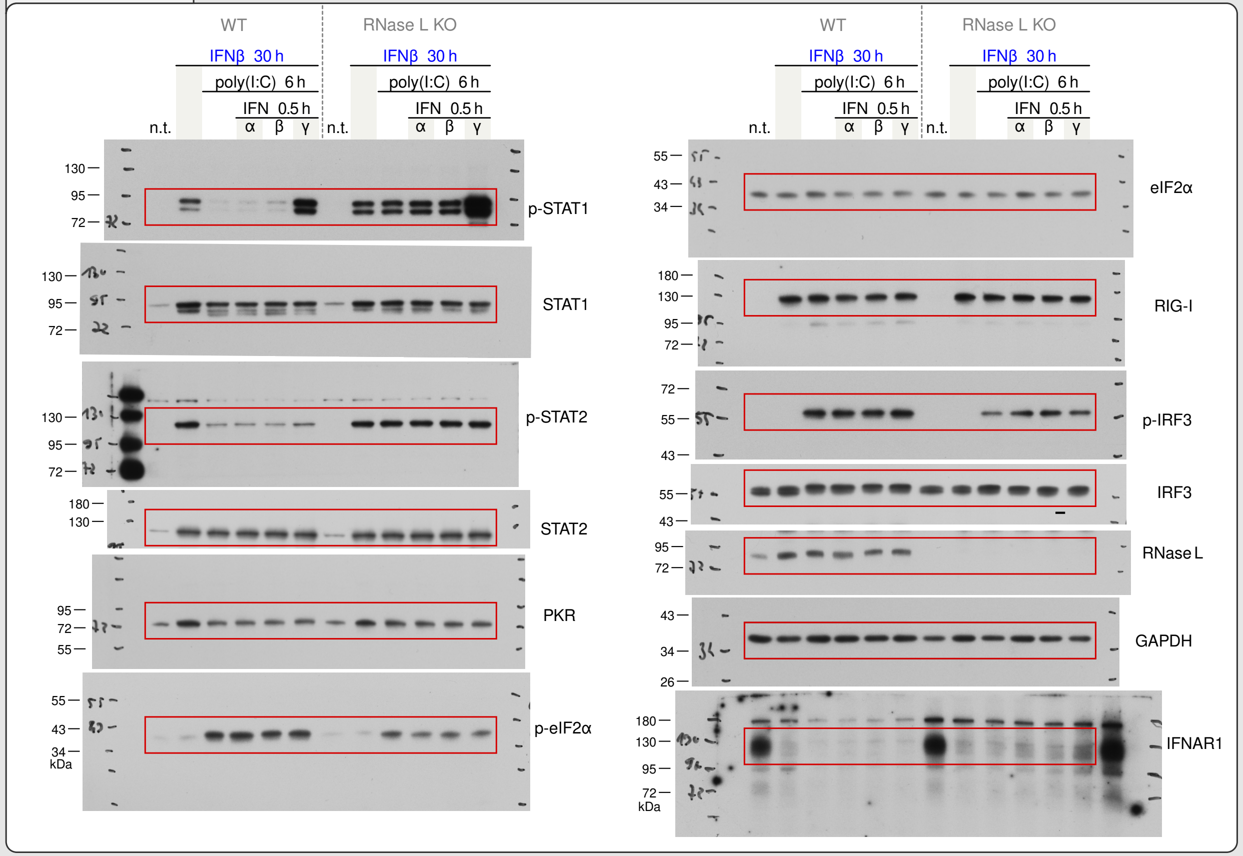

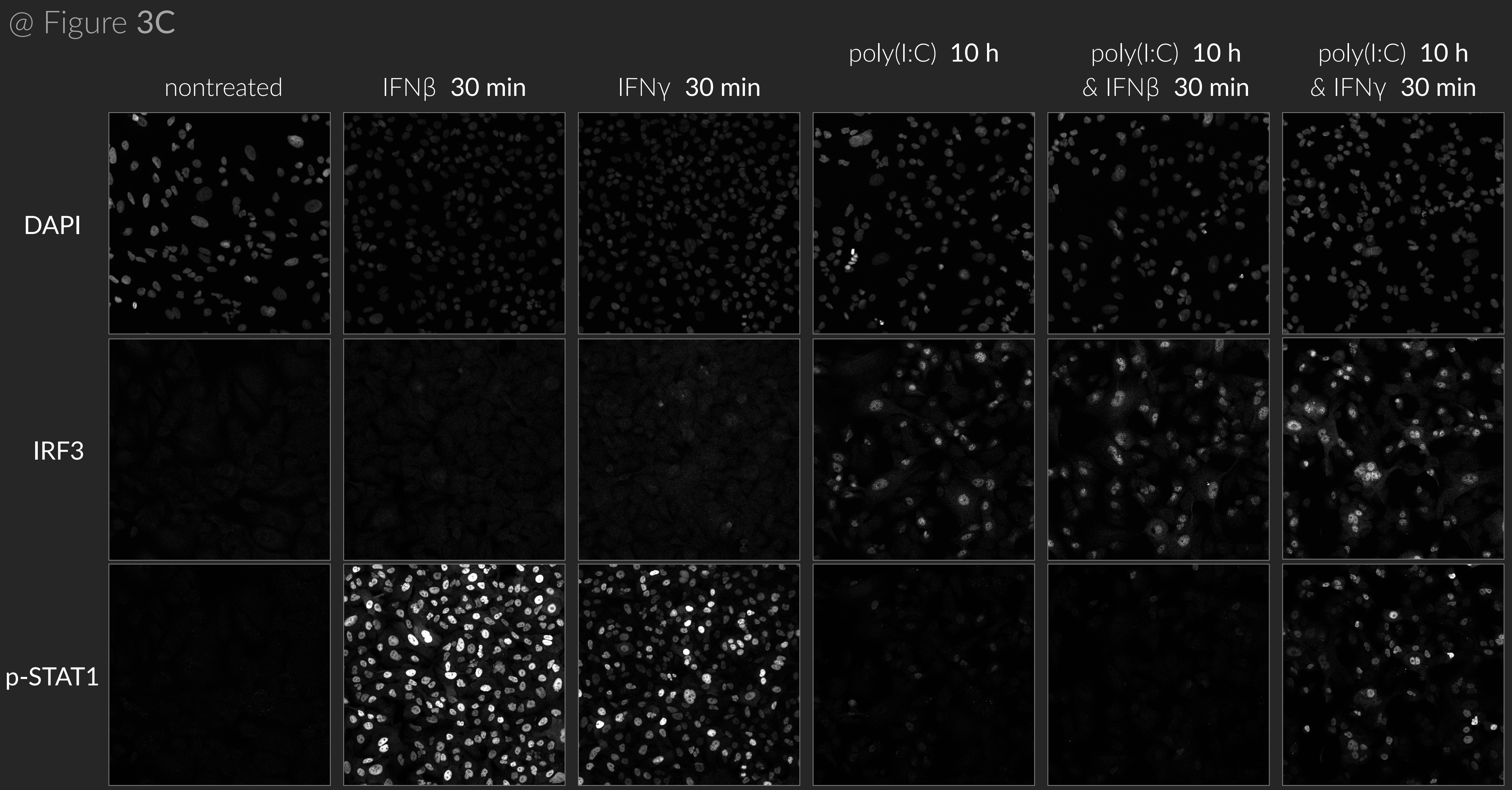
@ Figure 3C
nontreated
IFNβ 30 min
IFNγ 30 min
poly(I:C) 10 h
poly(I:C) 10 h
& IFNβ 30 min
poly(I:C) 10 h
& IFNγ 30 min
DAPI
IRF3
p-STAT1

@ Figure **4A**

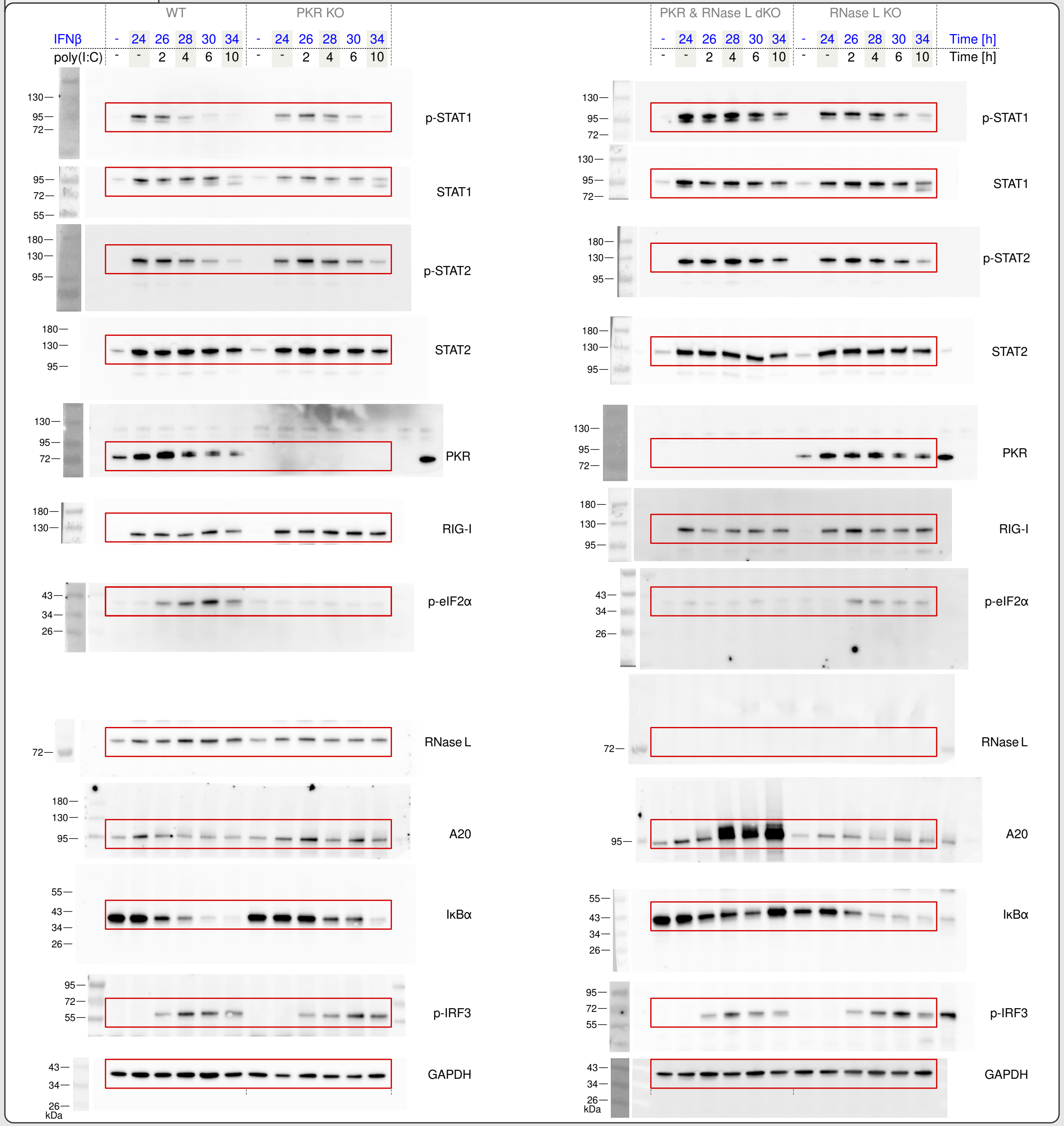
WT
PKR KO
PKR & RNase L dKO
RNase L KO
IFNβ
poly(I:C)
Time [h]
p-STAT1
STAT1
p-STAT2
STAT2
PKR
RIG-I
p-eIF2α
RNase L
A20
IκBα
p-IRF3
GAPDH
kDa

@ Figure **S1A**

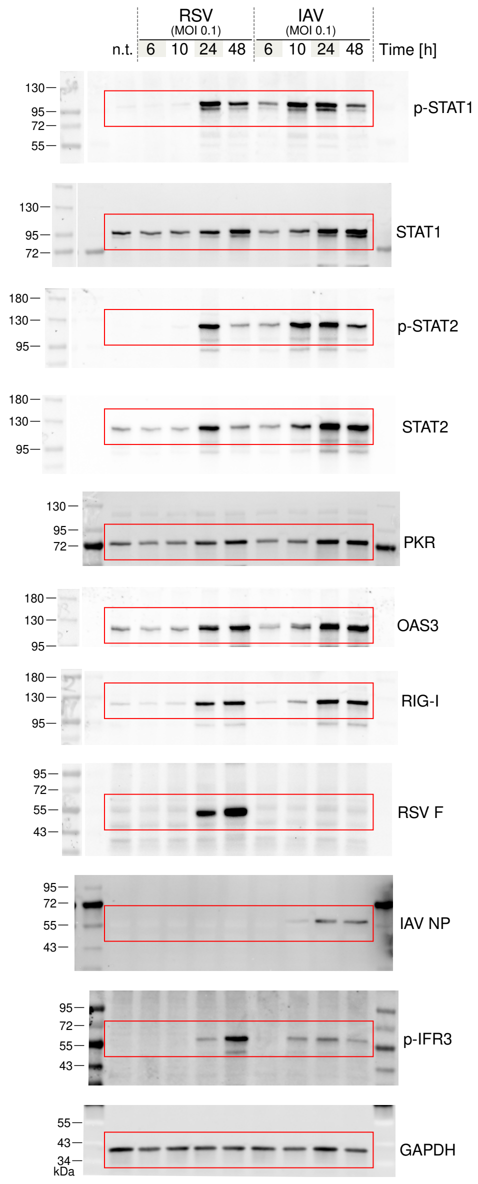

RSV
(MOI 0.1)
IAV
(MOI 0.1)
n.t.
6 10 24 48
6 10 24 48
Time [h]
p-STAT1
STAT1
p-STAT2
STAT2
PKR
OAS3
RIG-I
RSV F
IAV NP
p-IFR3
GAPDH
kDa

@ Figure **S2A**

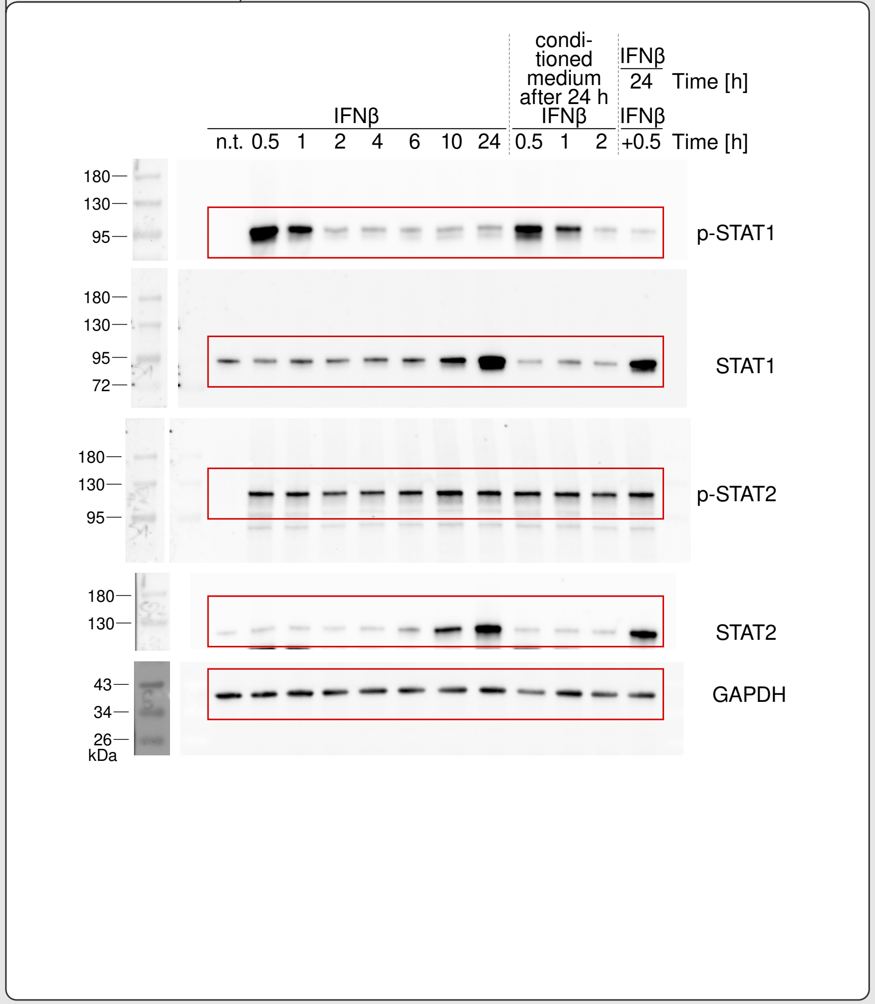


@ Figure **S2B**

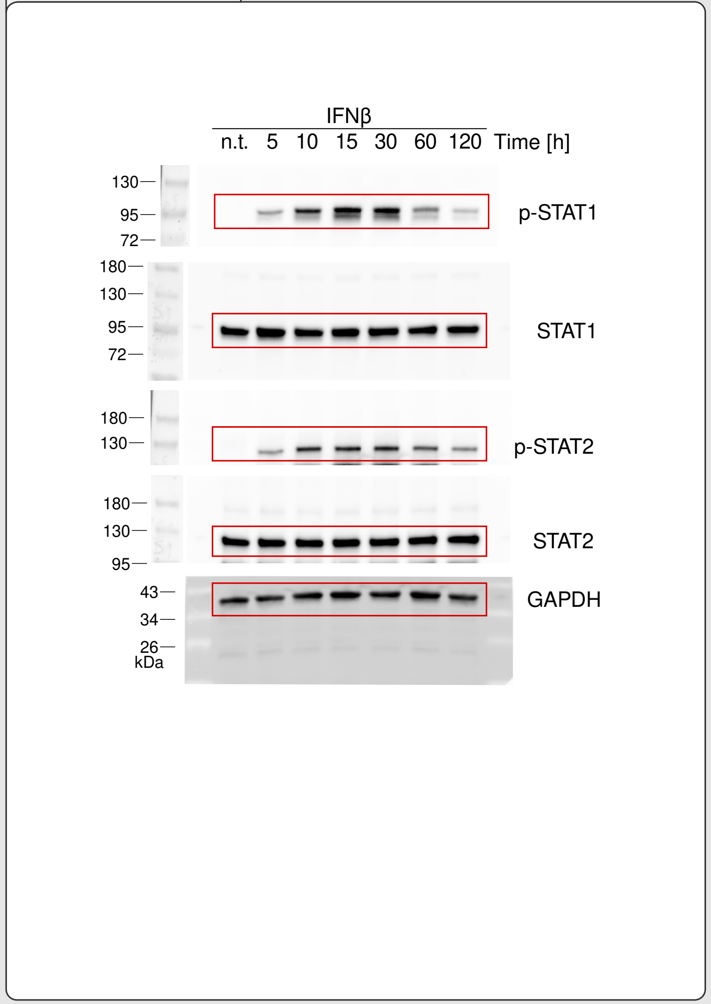


@ Figure **S2C**

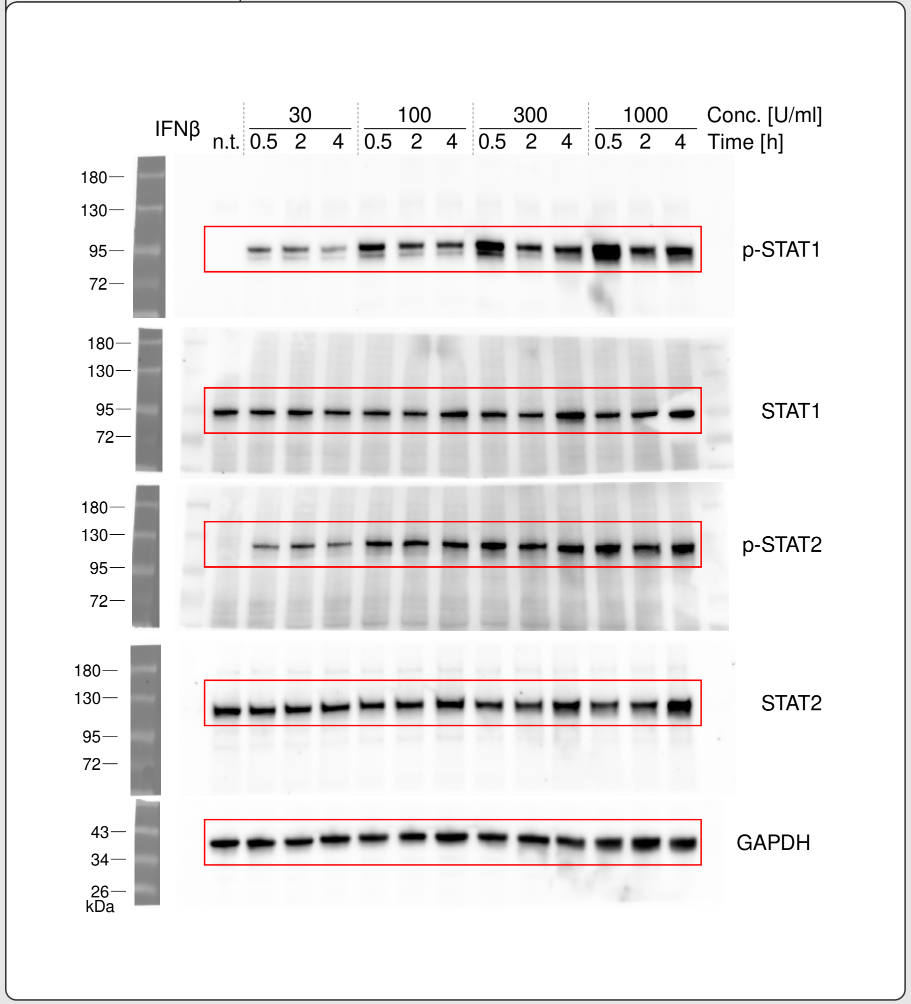

@ Figure **S5**

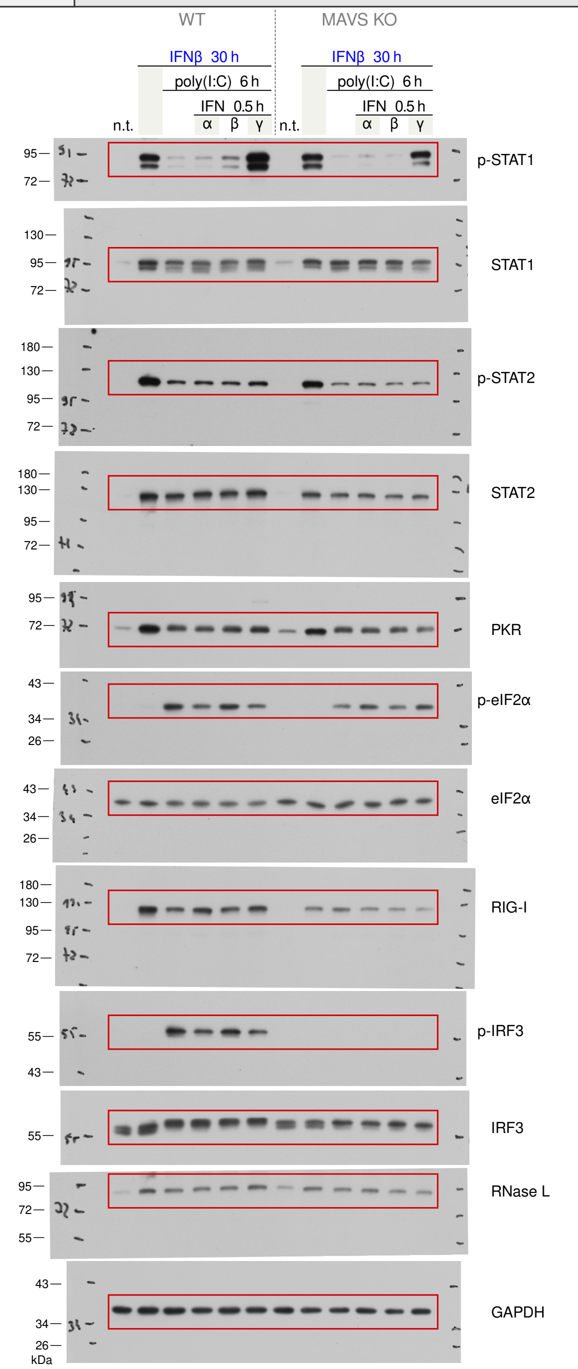
WT
MAVS KO
IFNβ 30 h
poly(I:C) 6 h
IFN 0.5 h
n.t.
α
β
γ
p-STAT1
STAT1
p-STAT2
STAT2
PKR
p-eIF2α
eIF2α
RIG-I
p-IRF3
IRF3
RNase L
GAPDH
kDa

@ Figure **S7A**

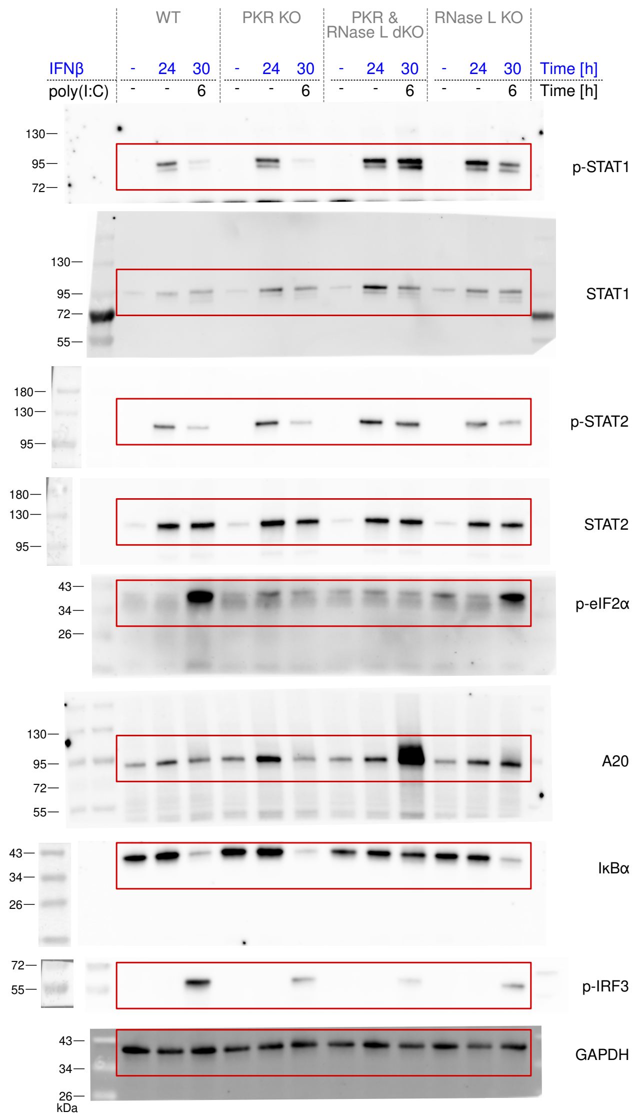

WT
PKR KO
PKR & RNase L dKO
RNase L KO
IFNβ - 24 30 - 24 30 - 24 30 - 24 30 Time [h]
poly(I:C) - - 6 - - 6 - - 6 - - 6 Time [h]
p-STAT1
STAT1
p-STAT2
STAT2
p-eIF2α
A20
IκBα
p-IRF3
GAPDH
kDa

@ Figure **S8A**

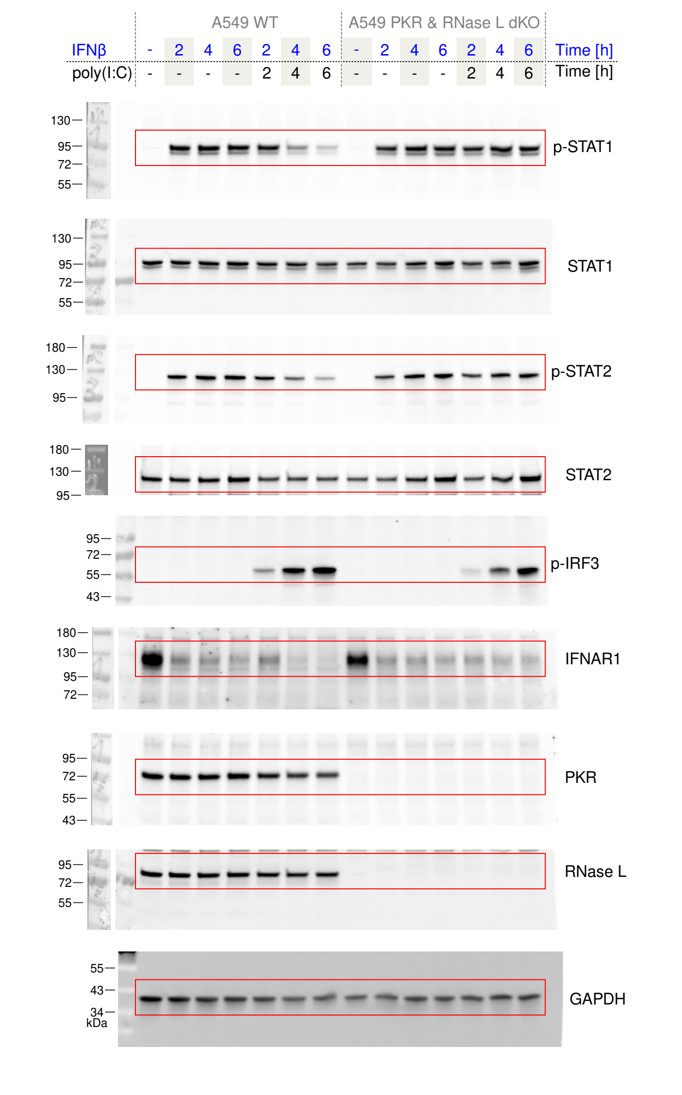


@ Figure **S8B**

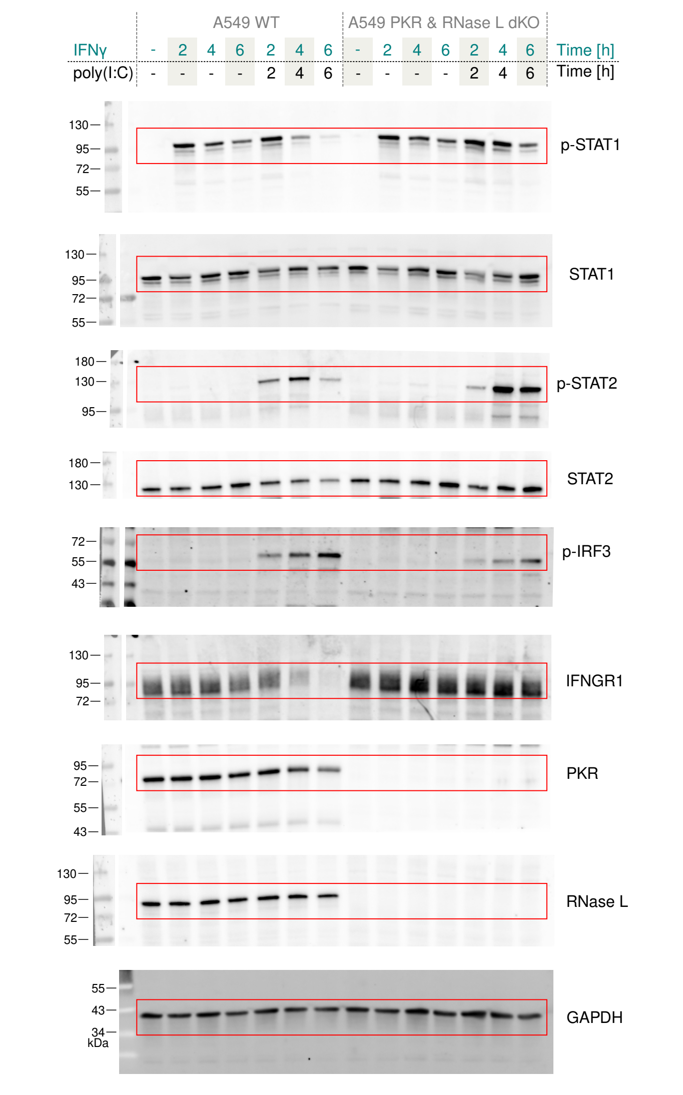

@ Figure **S9**

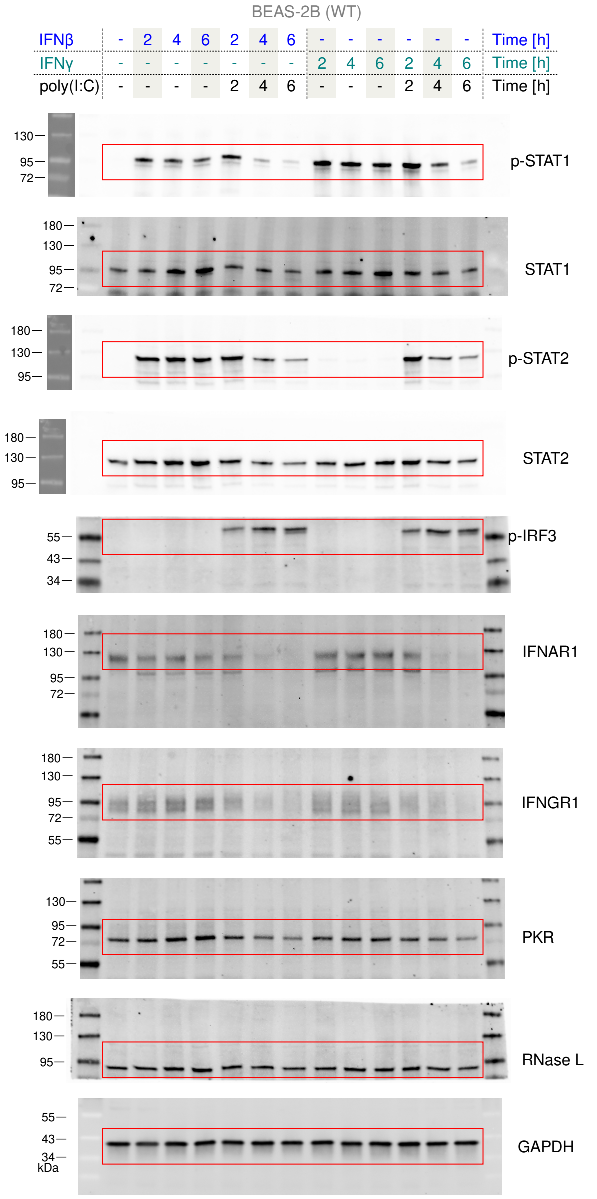
BEAS-2B (WT)
IFNβ - 2 4 6 2 4 6 - - - - - - Time [h]
IFNγ - - - - - - - 2 4 6 2 4 6 Time [h]
poly(I:C) - - - - 2 4 6 - - - 2 4 6 Time [h]
p-STAT1
STAT1
p-STAT2
STAT2
p-IRF3
IFNAR1
IFNGR1
PKR
RNase L
GAPDH
kDa

@ Figure **S11A**

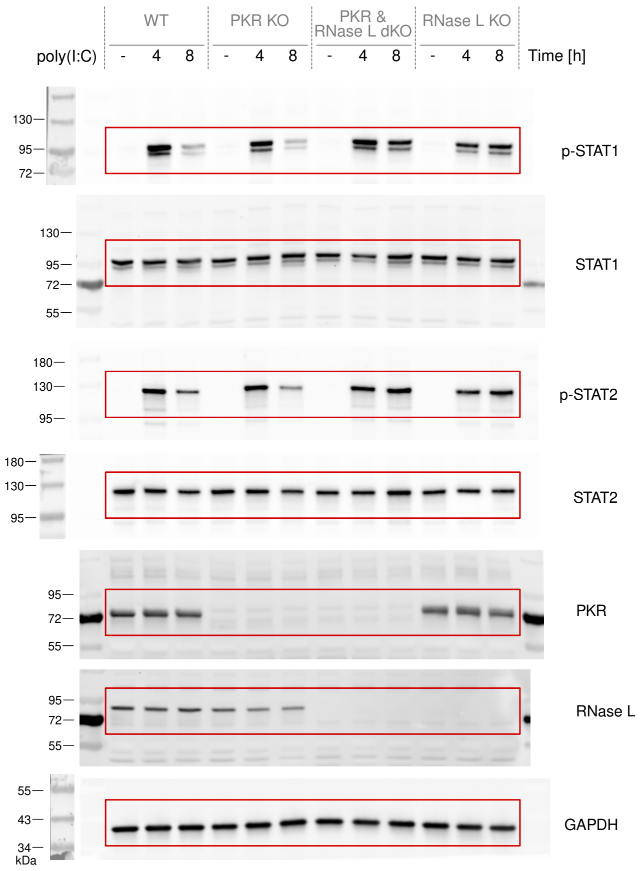
WT
PKR KO
PKR & RNase L dKO
RNase L KO
poly(I:C)
- 4 8 - 4 8 - 4 8 - 4 8
Time [h]
p-STAT1
STAT1
p-STAT2
STAT2
PKR
RNase L
GAPDH
kDa

@ Figure **S12A**

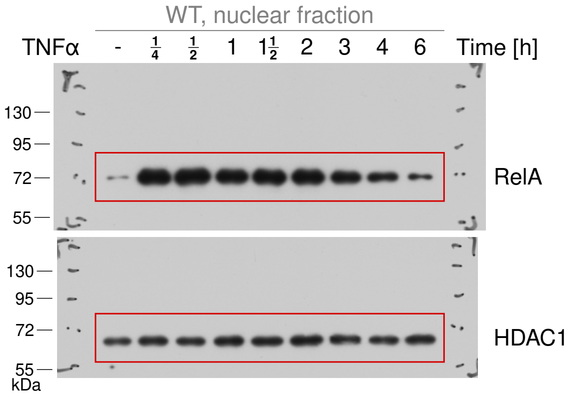


@ Figure **S12B**

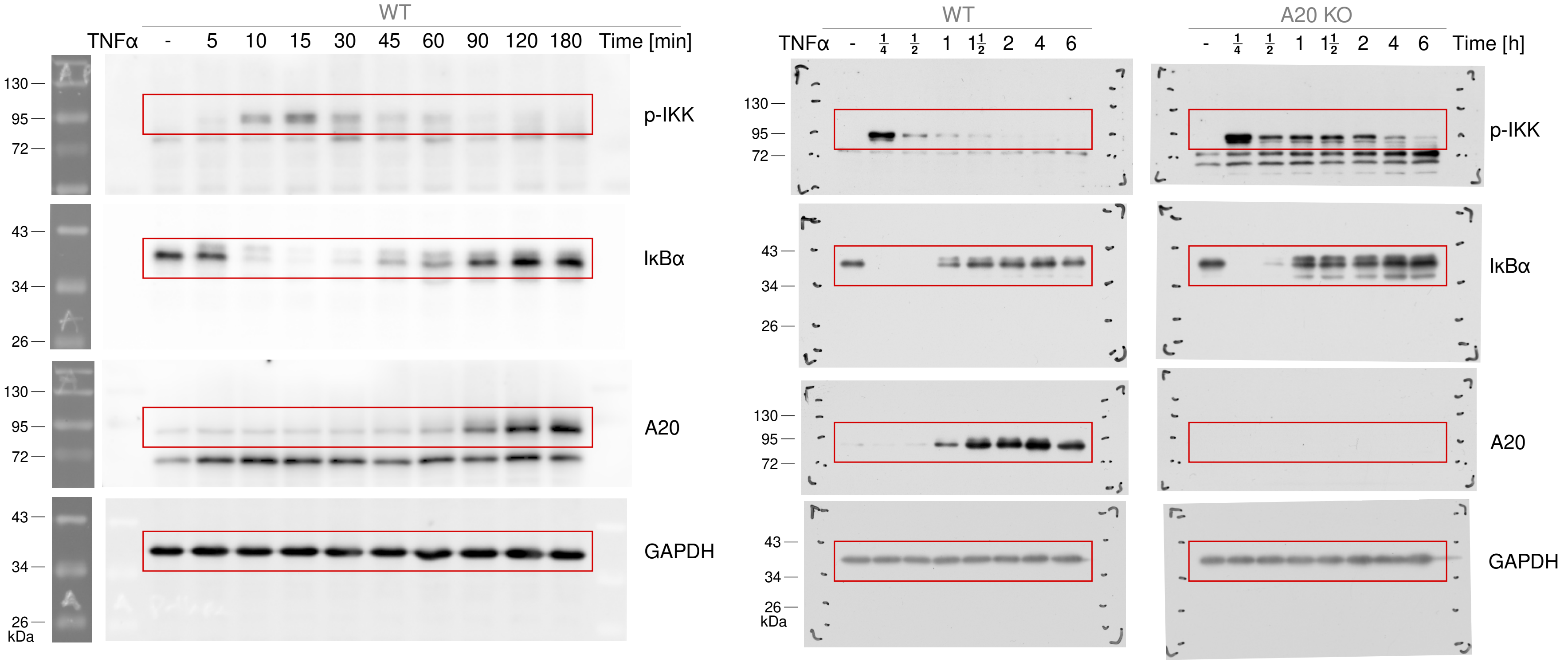

@ Figure **S17**

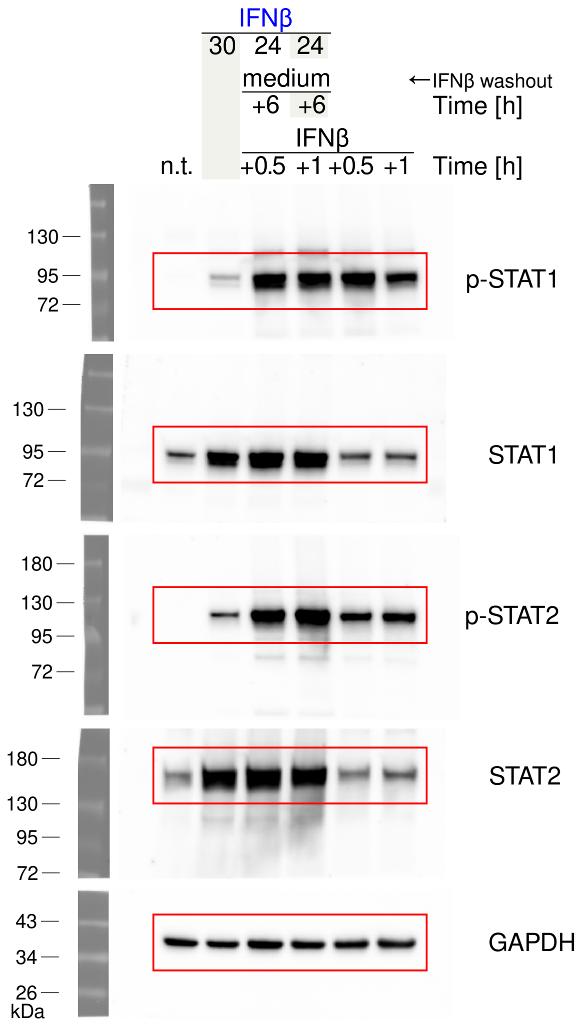
IFNβ
30 24 24
medium
+6 +6
←IFNβ washout
Time [h]
IFNβ
n.t.
+0.5 +1 +0.5 +1
Time [h]
130
95
72
p-STAT1
130
95
72
STAT1
180
130
95
72
p-STAT2
180
130
95
72
STAT2
43
34
26
kDa
GAPDH

@ Figure **S18**

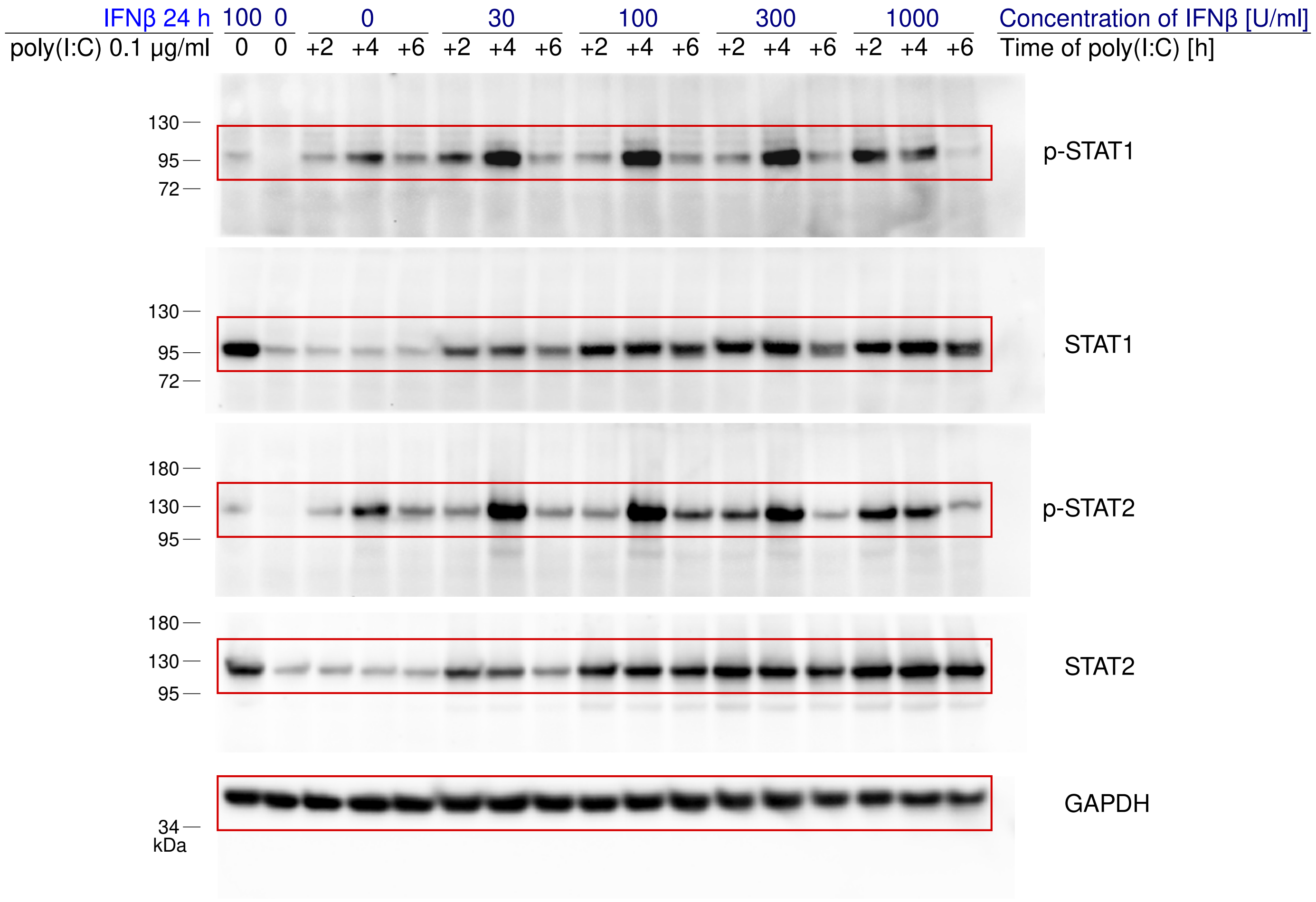

IFNβ 24 h 100 0 0 30 100 300 1000 Concentration of IFNβ [U/ml]
poly(I:C) 0.1 µg/ml 0 0 +2 +4 +6 +2 +4 +6 +2 +4 +6 +2 +4 +6 +2 +4 +6 Time of poly(I:C) [h]
130
95
72
p-STAT1
130
95
72
STAT1
180
130
95
p-STAT2
180
130
95
STAT2
GAPDH
34
kDa

@ Figure S19

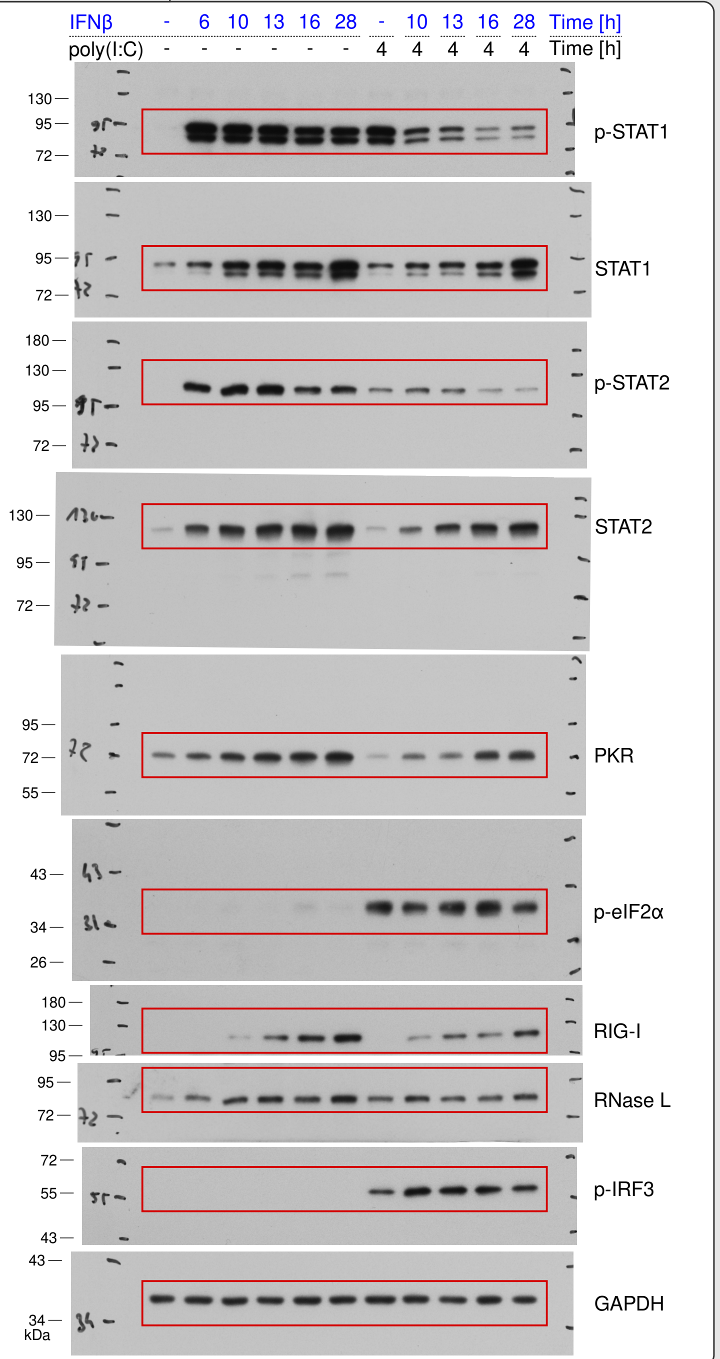
IFNβ - 6 10 13 16 28 - 10 13 16 28 Time [h]
poly(I:C) - - - - - - 4 4 4 4 4 Time [h]
p-STAT1
STAT1
p-STAT2
STAT2
PKR
p-eIF2α
RIG-I
RNase L
p-IRF3
GAPDH
kDa
130
95
72
180
55
43
34
26
